# High Axial Resolution Is Necessary for Quantitative Two-Photon Calcium Imaging of Neuronal Populations

**DOI:** 10.64898/2026.07.28.741086

**Authors:** Ha Yun Anna Yoon, Umaima Afifa, Génesis Ferrer Imbert, Adam S. Charles, Na Ji

## Abstract

Two-photon calcium imaging is a standard tool for measuring neuronal population activity *in vivo*, yet how axial resolution, sensor expression strategy, and analysis pipeline jointly affect data accuracy remains poorly characterized. Here, we imaged the same L2/3 neurons in mouse primary visual cortex at five axial resolutions (3.6–21.0 μm), spanning current two-photon systems from benchtop microscopes to large-field-of-view and miniaturized designs. We expressed cytosolic, transgenic, and soma-targeted GCaMP variants and applied five analysis pipelines. Reducing axial resolution systematically attenuated ΔF/F_0_, corrupted visual responsiveness and orientation tuning classifications, and biased pairwise correlations. No pipeline corrected these resolution-dependent artifacts, and pipeline choice alone produced quantitatively divergent results even at the highest resolution. Soma-targeted sensors mitigated but did not eliminate these artifacts. Our findings demonstrate that high axial resolution is necessary for accurate quantitative population imaging, and that robust separation of somatic from neuropil signals remains an unresolved challenge.

**In Brief:** Imaging the same cortical neurons across five axial resolutions and five analysis pipelines, Ji and colleagues show that lower resolution corrupts neuronal tuning and population correlations. No pipeline corrects these artifacts, and pipeline choice yields divergent results even at high resolution. Soma-targeted sensors mitigate but do not eliminate these failures.

## Introduction

Point-scanning two-photon fluorescence microscopy (2PFM) is the most widely used technique for *in vivo* imaging, capable of interrogating structure and function deep within intact, optically opaque tissues^1,2^. In 2PFM, two-photon (2P) absorption confines fluorescence generation to a small excitation volume created by focusing the excitation laser with a microscope objective of high numerical aperture (NA)^3,4^. The excitation focus is scanned across the lateral (xy) focal plane of the objective, and the fluorescence intensity recorded at each position maps the distribution and brightness of fluorophores within a thin optical section of the tissue. The thickness of this optical section is set by the axial size of the excitation focus, which is in turn determined by the excitation NA.

In neuroscience, this optical sectioning capability has established 2PFM as a central tool for functional population imaging, enabling cellular-resolution activity measurements of large neuronal populations deep within scattering brain tissue *in vivo*. In typical 2P calcium imaging, the cytosol of neuronal somata and neuropil is labeled with calcium sensors, fluorophores whose brightness changes in response to variations in intracellular calcium concentration caused by action potential firing. With fluorophores distributed throughout the tissue volume, achieving reliable measurement of neuronal calcium responses depends critically on the axial extent of the excitation focus. Early applications of 2PFM for population calcium imaging often used a microscope objective of 0.8 NA^5–7^, which can form an excitation focus with a theoretical axial full width at half maximum (aFWHM) of <3 μm (e.g., 2.1 μm for 900 nm excitation light^4,8^). However, practical aFWHMs often exceed this ideal due to underfilling of the objective, microscope aberrations, or sample-induced aberrations^3,9,10^. In superficial cortical layers (e.g., layer 2/3, L2/3, of the mouse cortex), well-designed microscopes typically maintain an effective aFWHM, or axial resolution, of 5 μm or better^4,10–13^. Given that the somata of mouse L2/3 neurons are typically 10 to 20 μm in size^14^, such an aFWHM is sufficient to achieve cellular resolution.

More recently, however, developments in 2PFM have increasingly traded high spatial resolution for improvements in other performance metrics. By reducing the excitation NA, one can more easily design scanning optics and microscope objectives to increase the field of view (FOV)^11,15–27^, modulate the divergence of the excitation light for fast axial scanning^28–39^, or miniaturize 2PFM for head-mounted imaging in freely moving animals^40–62^ (**Table S1**). Among these systems, aFWHMs up to 41.5 μm were used for 2PFM, considerably larger than the ∼5 μm typical of conventional 2PFMs. With such an axially elongated excitation focus, individual neurons can still be resolved laterally, but cellular resolution is lost along the axial dimension.

This loss of axial resolution has important consequences for how calcium signals are interpreted. In calcium imaging, resolution, a microscope’s ability to distinguish signals from two closely spaced objects, determines how accurately the activity of a neuron of interest can be isolated from transients in its surrounding structures, including neuropil (i.e., axons and dendrites of neurons) and adjacent cell bodies. An axially elongated excitation focus causes a loss of resolution and exacerbates neuropil contamination^63^ and signal mixing, which, if not properly corrected, leads to inaccurate measurements of neuronal activity.

In parallel with these hardware trends and in response to increasingly large datasets, methods for calcium imaging data analysis have also advanced. While many studies still manually segment regions of interest (ROIs) corresponding to somata, analysis pipelines have been developed for automatic cell detection, fluorescence extraction, neuropil subtraction, and, in some cases, demixing of signals from overlapping neurons^64–69^. These methods were developed and tested on high-NA calcium imaging datasets. However, they are increasingly being applied to lower-resolution data (**Table S1**), a regime in which their performance has yet to be systematically validated.

Given the increasing popularity of lower-resolution 2PFM systems, it has become essential to understand how reducing axial resolution impacts the accuracy of *in vivo* calcium imaging and whether existing analysis pipelines can mitigate the resulting errors. We acquired calcium imaging data at different aFWHMs from the same L2/3 neurons in the mouse primary visual cortex (V1) expressing either GCaMP6s throughout soma and neuropil (“cytosolic GCaMP6s”) or soma-targeted jGCaMP8s. We then analyzed these data using five different analysis pipelines applied to either manually segmented or automatically detected ROIs.

We found that for both cytosolic and soma-targeted sensor expression, reducing axial resolution degraded structural image quality, attenuated somatic calcium transient magnitudes, and exacerbated neuropil contamination, leading to errors in the functional classification and network activity characterization of V1 neurons that none of the pipelines were able to eliminate. Network statistics such as pairwise orientation tuning or activity correlation over distance were highly sensitive to the choice of analysis pipeline, even for calcium data acquired at the highest axial resolution, indicating that careful attention to neuropil-induced artifacts is warranted in such analyses. Notably, neurons expressing soma-targeted calcium sensors were substantially dimmer than those expressing cytosolic sensors but were more resilient to neuropil contamination artifacts. In conclusion, high axial resolution is necessary for accurate *in vivo* calcium imaging of neuronal populations when using cytosolic calcium sensors. For lower-resolution microscopes, restricting sensor expression to the somata or nuclei, combined with proper choice of analysis pipeline, may help reduce these artifacts.

## Results

### Effect of aFWHM on the signal of beads of different sizes

To systematically investigate the effect of axial resolution on signal intensity, we varied the excitation NA and axial resolution of a custom-built 2P fluorescence microscope^70^ using a motorized beam reducer upstream of the microscope (**Figure 1a**) to underfill the back pupil plane of the objective lens with 920 nm excitation light (**Methods**).

**Figure 1.**
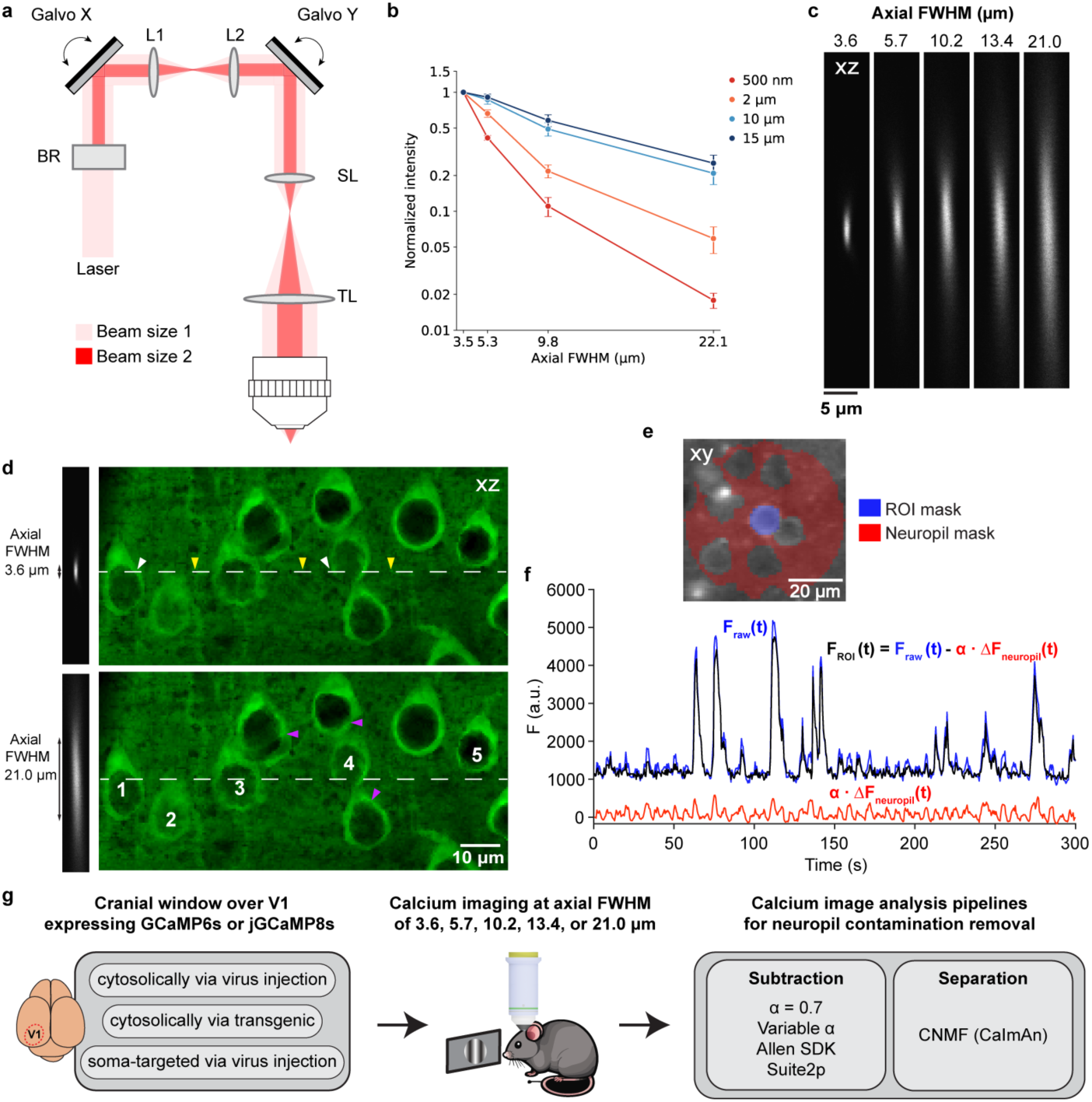
2PFM signal, axial resolution, and calcium image analysis pipelines. **(a)** Optical schematic showing two excitation beam sizes (light red and red). A motorized beam reducer (BR) underfills the objective back pupil, reducing excitation NA and axial resolution. **(b)** Fluorescence signals from beads of different diameters versus axial FWHM (30 beads per aFWHM), normalized by the square of post-objective excitation power and to the signal measured at 3.5-µm aFWHM. Error bars: s.e.m. **(c)** Axial images of a 2-µm-diameter bead with five axial FWHM (aFWHM) values. Images are individually normalized. **(d)** Axial bead images with aFWHM of 3.6 µm and 21.0 µm, respectively, alongside a simulated brain volume. White dashed line: focal center; white, yellow, purple arrowheads: neurons within the 3.6-µm focus, neuropil within the 3.6-µm focus, and additional neurons within the 21.0-µm focus, respectively. **(e)** Example ROI mask (blue) and corresponding neuropil mask (red, 30-µm radius from the ROI centroid, excluding pixels associated with other neurons). **(f)** Neuropil subtraction: the neuropil transient, ΔF_neuropil_(t), was scaled by α = 0.7 and subtracted from the raw ROI fluorescence trace, F_raw_(t), to obtain F_ROI_(t)=F_raw_(t) - ⍺ΔF_neuropil_(t). **(g)** Experimental workflow. GCaMP6s was expressed in V1 neurons by virus injection or transgenically, and soma-targeted jGCaMP8s was expressed by virus injection. 2P calcium imaging of the same fields of view in V1 was then performed at five aFWHM values in head-fixed awake mice. Data was then analyzed using different pipelines, with neuropil contamination removed by subtraction or CNMF.

First, we measured how aFWHM affected the maximal pixel values, i.e., signals, in axial images of fluorescent beads with diameters of 0.5, 2, 10, and 15 µm (normalized to the signal at 3.5-µm aFWHM and the same post-objective excitation power, **Figure 1b**, **Table S2,** N = 30 beads per condition), corresponding to neuronal structures ranging in size from synaptic specializations to cell bodies. The signal of 0.5-µm-diameter beads decreased drastically with the degradation of resolution: as aFWHM increased from 3.5 µm to 5.3, 9.8, and 22.1 µm, the signal decreased by 2.4×, 9.0×, and 55.5×, respectively. This decrease became more gradual with increasing bead size. For 2-µm beads, the signal decreased by 16.9× from 3.5- to 22.1-µm aFWHM. For 10- and 15-µm beads, the signal decreased much more gradually: as aFWHM increased from 3.5 to 9.8 µm, the signal decreased by 2.0× and 1.7×, respectively, and as it increased further to 22.1 µm, the signal decreased by 4.8× and 3.9×, respectively.

These observed trends can be understood by considering, first, how focal intensity and the intensity distribution within the focal volume vary with NA and, second, what determines the fluorescence signal at each pixel. A high-NA focus concentrates excitation power into a small volume, yielding a high peak intensity and rapid drop-off in intensity away from the focal center. Conversely, decreasing NA reduces peak intensity but expands the excitation volume both axially and laterally. The fluorescence signal at each image pixel is proportional to the number of fluorophores within the focal volume and the squared excitation intensity at their locations, and thus depends on the size and position of the fluorescent object relative to the focal volume.

For beads imaged with a diffraction-limited excitation focus, the maximal pixel value is recorded when the bead’s center coincides with the focal center. Beads of 0.5-µm diameter were contained within the excitation focus under all five conditions. As NA decreases and aFWHM increases, the number of fluorophores within the focus stays constant but the excitation intensity experienced by these fluorophores decreases, leading to the observed monotonic decay of maximal pixel values (**Figure 1b**).

For 2-µm-diameter beads, the number of fluorophores within the excitation focus first increases with the lateral expansion of the focus, then plateaus once the bead is fully contained within the excitation focus at larger aFWHM values. However, this increase in fluorophore number is insufficient to compensate for the drop in peak excitation intensity at larger aFWHMs, so pixel values also decrease monotonically with aFWHM, though less precipitously than for submicron beads.

For large beads of 10- and 15-µm diameter, increasing aFWHM initially recruits more fluorophores into the excitation volume. At intermediate aFWHMs (e.g., 5.3 µm), this gain in fluorophores can mostly compensate for the drop in peak excitation intensity, leading to a slight drop in signal (1.2× and 1.1× for 10-µm and 15-µm beads, respectively). At 9.8- and 22.1-µm aFWHMs, however, the reduction in peak intensity becomes too large to be compensated by the increased number of excited fluorophores, resulting in a moderate decrease in signal.

### Larger aFWHMs cause more severe neuropil contamination and signal mixing

The bead measurements above demonstrate that high-NA excitation is clearly advantageous for maximizing signal when imaging structures such as spines and boutons. However, signal from objects of cell-body dimensions declines only modestly as aFWHM increases, and this robustness raises the question of which axial resolution is appropriate for *in vivo* calcium imaging of neuronal populations.

To address this question, we considered five aFWHM values of 3.6, 5.7, 10.2, 13.4, and 21.0 µm, measured from axial images of a 2-µm-diameter fluorescent bead (**Figure 1c**), spanning the range most commonly reported in the literature (**Table S1**). (The actual axial resolution is slightly better than these values, owing to our use of relatively large, 2-µm-diameter beads.)

Considered in isolation, the bead data appear to favor larger aFWHMs: for neuronal cell bodies, which are typically 10-20 µm in diameter, increasing aFWHM from 3.6 µm to 10.2 µm reduces somatic signal by only half. On this basis, one could justify enlarging aFWHM and accepting a moderate loss of signal in exchange for gains in other performance metrics, such as imaging speed, FOV, volumetric coverage, or system portability (e.g., for miniaturized microscopes).

Signal magnitude, however, is not the only consideration. An axially elongated excitation focus broadens the volume from which fluorescence is generated at each pixel, increasing the likelihood that signal from a soma of interest is contaminated by surrounding neuropil or mixed with signal from axially adjacent cell bodies. The bead measurements alone cannot capture these effects, because beads are sparse and isolated, whereas neurons are embedded in a dense, fluorescent neuropil.

To illustrate this point, we juxtaposed axial bead images of 3.6 µm and 21.0 µm aFWHM with axial views of neurons simulated by NAOMi^71^ using an average cell body diameter of 11.8 µm (**Figure 1d**). With 3.6-µm-aFWHM excitation, fluorescence at each pixel predominantly originated from a single source (white arrowheads: cell body; yellow arrowheads: neuropil; **Figure 1d**). In contrast, with 21.0-µm-aFWHM excitation, fluorescence rarely originated from a single source: somatic signals were contaminated by substantial neuropil (Cells 1, 2, and 5; **Figure 1d**) and/or, for neurons in the center of the focal volume, mixed with signals from axially neighboring neurons (purple arrowheads, Cells 3 and 4; **Figure 1d**).

### Commonly used calcium image analysis pipelines for neuropil subtraction and signal demixing

Calcium transients from somata often differ from those of the surrounding neuropil^63,72^. If not properly corrected, this neuropil contamination can lead to inaccurate measurements of somatic activity. Various approaches^64–69,73–76^ have been developed to mitigate neuropil contamination in 2P calcium imaging data.

The most conceptually straightforward method is neuropil subtraction, in which an estimate of neuropil contamination is subtracted from the somatic signal. Because the neuropil estimate is measured with the excitation focus entirely within neuropils laterally adjacent to the soma (see below), whereas the somatic measurement contains both soma and neuropil within its focal volume, the neuropil estimate is scaled by a coefficient ⍺ before subtraction. Several approaches have been used to estimate this coefficient.

#### Hand-segmented ROIs and neuropil subtraction

In a common approach, an ROI is hand-segmented to encompass the cell body (blue pixels, **Figure 1e**), and pixel values within this ROI are averaged to obtain F_raw_ for each image frame. From the calcium imaging time series, the fluorescence trace over time, F_raw_(t), is then extracted (blue trace, **Figure 1f**). If neuropil contamination is ignored, a baseline fluorescence value F_0,raw_ is determined from F_raw_(t), followed by calculation of ΔF_raw_(t)=F_raw_(t) - F_0,raw_ and finally ΔF_raw_(t)/F_0,raw_, which represents the calcium response of the ROI.

To correct for neuropil contamination, a neuropil mask is generated for each ROI that includes pixels surrounding the ROI while excluding pixels that fall within any cell body (red pixels, **Figure 1e**). A neuropil fluorescence trace F_neuropil_(t) is then extracted by averaging the pixel values within the neuropil mask for each frame. A baseline F_0, neuropil_ is determined from F_neuropil_(t), and ΔF_neuropil_(t) = F_neuropil_(t) − F_0, neuropil_ can be calculated.

In the literature, the neuropil-corrected calcium trace is often calculated as F_ROI_(t)=F_raw_(t) - ⍺F_neuropil_(t) with ⍺ set to 0.7^69,74,76–78^. However, we found that, for samples with strong neuropil expression, as is often the case in population calcium imaging, subtracting F_neuropil_(t) can lead to artifactually small F_0_ values and erroneously large ΔF/F_0_. We therefore subtracted ΔF_neuropil_(t) instead of F_neuropil_(t). For an example neuron from a dataset acquired at 3.6-µm aFWHM (**Figure 1f**), neuropil subtraction using F_ROI_(t)=F_raw_(t) - ⍺ΔF_neuropil_(t) with ⍺ = 0.7 reduced the amplitudes of small calcium transients in the ROI trace (black trace, **Figure 1f**), presumably arising from contamination by neuropil activity (red trace, **Figure 1f**).

Although ⍺ is often set as a constant for all datasets, methods have also been developed to determine ⍺ for individual neurons. The rationale is that, depending on the position of the excitation focus relative to the cell body, neuropil contribution can vary across cells. For example, ⍺ can be determined for each ROI via linear regression between its neuropil fluorescence and the lowest 5% of raw somatic fluorescence^73^ (“Variable ⍺”), or by minimizing the cross-validated error under a smoothness constraint^75,79^ (“Allen SDK”). We thus applied three neuropil-subtraction methods to hand-segmented ROIs: Variable ⍺, Allen SDK, and a fixed ⍺ of 0.7 (“⍺ = 0.7”).

#### Automatically detected ROIs and neuropil subtraction by Suite2p

Suite2p^65^, a popular calcium image analysis pipeline, automatically detects cellular ROIs and generates masks for their surrounding neuropil. Instead of simply averaging pixel values within the ROI mask to obtain F_raw_(t), before averaging, Suite2p weights the pixel values by their variance explained by the extracted trace. It also subtracts F_neuropil_(t), rather than ΔF_neuropil_(t), using F_ROI_(t)=F_raw_(t) - ⍺F_neuropil_(t) with a default ⍺ value of 0.7.

#### Automatically detected ROIs, neuropil contamination removal, and signal demixing by CNMF

In addition to subtraction-based methods, analysis pipelines have also been developed to remove neuropil contamination and demix activity from overlapping neurons^64,66–69^. Methods based on constrained non-negative matrix factorization (CNMF) represent the spatiotemporal activity of each ROI as the outer product of a spatial component encoding the neuronal footprint and a temporal component describing its activity trace. Calcium imaging time series are then modeled as the sum of the spatiotemporal activity of all ROIs plus non-sparse, low-rank terms that describe background fluorescence and neuropil activity. The inferred activity traces for each neuron have neuropil contamination already removed, and the activity of partially overlapping neurons can also be demixed.

Importantly, all of these approaches were initially developed for high-resolution calcium imaging data collected with 0.8, 1.0, or 1.05 NA objective lenses^64,69,73,76^, for which neuropil contamination is less severe and signals from individual neurons are more readily separable. Whether they effectively remove the stronger neuropil contamination or demix activity from overlapping neurons at larger aFWHM values has not been systematically investigated. Given the increasing adoption of lower-resolution 2PFM systems and the growing application of these analysis methods to datasets acquired at large aFWHM values (**Table S1**), there is an urgent need for a systematic characterization of imaging performance and a rigorous evaluation of commonly used calcium imaging analysis tools. Below, we performed calcium imaging in mouse V1 L2/3 across a range of aFWHM values, using neurons expressing either cytosolic GCaMP6s^74^ (via intracortical viral transduction or transgenically) or soma-targeted jGCaMP8s^80^, to determine whether these commonly applied approaches adequately remove neuropil contamination and permit quantitatively accurate characterization of neuronal activity (**Figure 1g**).

### Calcium imaging of V1 L2/3 neurons with virally transduced cytosolic GCaMP6s using different aFWHMs analyzed by hand segmentation and neuropil subtraction

Following a common experimental preparation, we expressed GCaMP6s in the cytosol of L2/3 neurons of wild-type mice by injecting AAV2/1.syn.GCaMP6s into V1, followed by cranial window implantation (see **Table S3** and **Methods** for details). After 3 weeks of expression and habituation to head-fixation, we imaged L2/3 neurons in awake mice during the presentation of drifting grating stimuli (10 trials, 72 s per trial, each trial consisting of 12 gratings drifting in one of 12 directions for 3 s, each preceded by a 3-s-long black screen; gratings were presented in pseudorandom sequence within each trial). Each 400 µm × 400 µm FOV was imaged five times, at aFWHMs of 3.6, 5.7, 10.2, 13.4, and 21.0 µm, at a 2 Hz frame rate and 1 µm pixel size (**Movie S1**). The five datasets were motion-corrected using NoRMCorre^81^ and registered to a common reference image: the motion-corrected and time-averaged image of the dataset acquired at 3.6-µm aFWHM. Pixel values reported in this manuscript were normalized by the post-objective power used during image acquisition.

#### Increasing aFWHM degrades both structural image quality and cellular calcium signals

Reducing axial resolution leads to a visible degradation of structural image quality. Time-averaged images of an example FOV across the five aFWHM conditions revealed a progressive loss of resolution and contrast (**Figure 2a**). At larger aFWHM values, fewer neurons showed the nucleus-excluded GCaMP6s expression than at 3.6-µm aFWHM, because the axially elongated focus increasingly excited fluorescent neuropil and the other cell bodies above and below the neuron. Additionally, as aFWHM increased, unlabeled blood vessel segments that were out of focus at 3.6-µm aFWHM encroached into the excitation volume and cast increasingly prominent shadows. Although some neurons remained resolved in these lateral images (e.g., ROI 1, **Figure 2a**), most became harder to discern at larger aFWHM values: some vanished into the shadows of blood vessels (e.g., ROI 2, **Figure 2a**), while others lost contrast against surrounding fluorescent structures (e.g., ROIs 3,4, **Figure 2a**).

**Figure 2.**
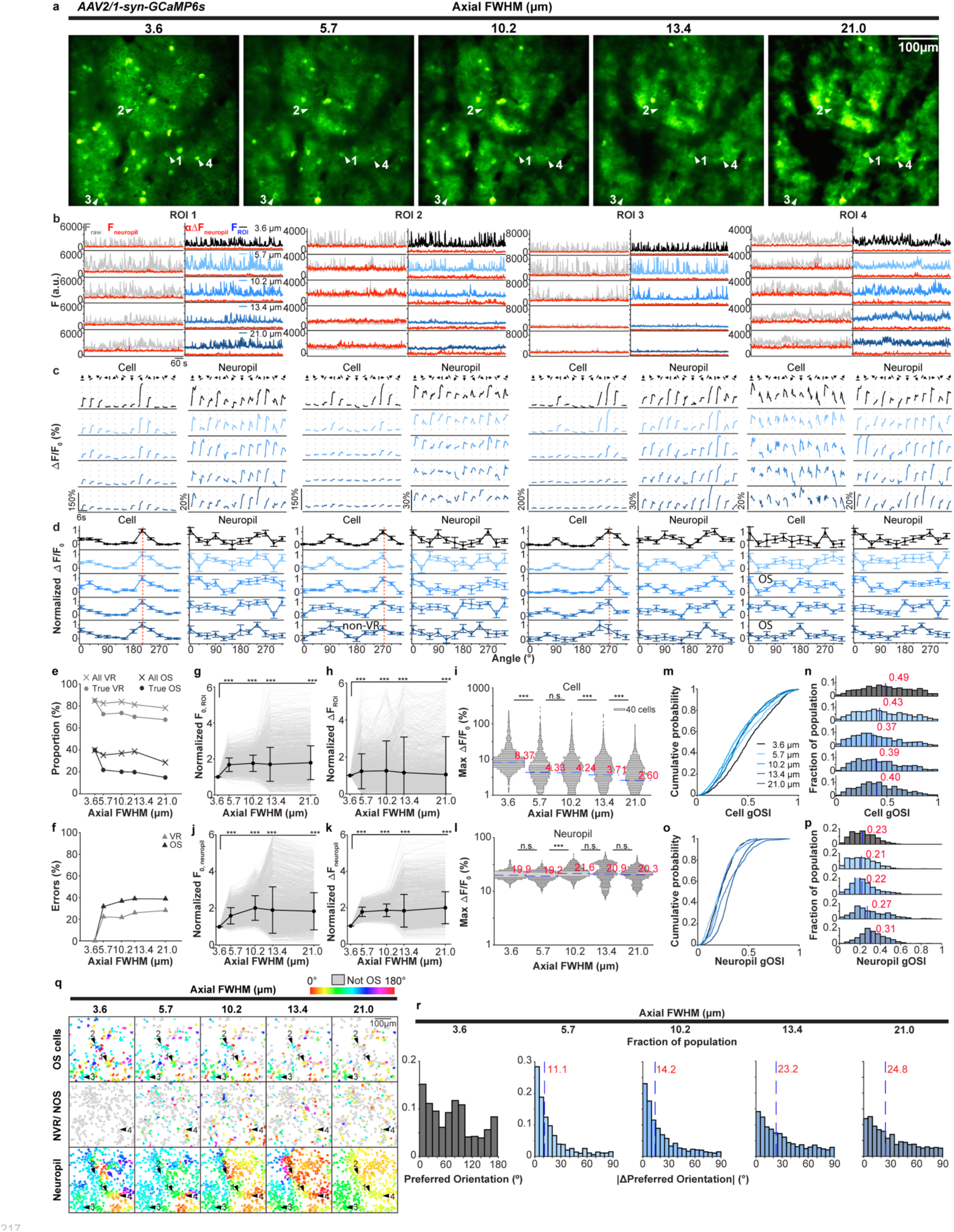
Degradation in axial resolution leads to increase in neuropil contamination and errors in functional characterization of neurons. **(a)** 2P fluorescence images of an example FOV of mouse V1 L2/3 neurons labeled with AAV2/1.syn.GCaMP6s acquired at five aFWHM values of 3.6, 5.7, 10.2, 13.4, and 21.0 µm. Arrowheads, example neurons: Cell 1 is classified as visually responsive (VR) and orientation-selective (OS) under all conditions; Cell 2 is misclassified as not VR as it moves into the shadow of the blood vessel at large aFWHM; Cell 3 exhibits changes in tuning curve and a shift in preferred orientation at large aFWHM; Cell 4 is misclassified as OS as aFWHM elongates. **(b)** Representative temporal traces of F_raw_ (gray), F_neuropil_ (red), αΔF_neuropil_ (red), and F_ROI_ (blue to black, color-coded by aFWHM) for Cell 1-4. α= 0.7. **(c)** Trial-averaged ΔF/F_0_ traces for Cell 1-4 and their neuropil. Gray dotted lines: onset of grating stimuli. **(d)** Tuning curves based on trial- and time-averaged ΔF/F_0_ during grating presentation for Cell 1-4 and their neuropil. Normalized individually to maximal ΔF/F_0_ values. Red dashed lines: preferred grating angles. **(e)** Out of 2,499 neurons, fraction of cells classified to be VR (gray cross) or OS (black cross) from images acquired with different aFWHM values; fraction of cells that are truly VR (gray dot) or OS (black dot). **(f)** Fraction of neurons that are misclassified in terms of VR (gray triangle) and OS (black triangle) properties. **(g)** F_0, ROI_ and **(h)** ΔF_ROI_ of 2,499 neurons for all aFWHM conditions, normalized to the value at 3.6 µm aFWHM (gray lines). Black dots: mean; error bars: s.d. Two-sample Kolmogorov-Smirnov (KS) test: ***p < 0.001. **(i)** For 2,122 neurons classified to be VR at 3.6-µm aFWHM, the distributions of their maximal trial-averaged ΔF/F_0_ across conditions. Red numbers and blue lines: median values. One-sided Wilcoxon rank sum (WRS) tests against previous aFWHM conditions: ***p < 0.001. **(j)** F_0, Neuropil_ and **(k)** ΔF_neuropil_ of neuropil associated with the 2,499 neurons. **(l)** Distributions of maximal trial-averaged ΔF/F_0_ for neuropil associated with 2,122 VR neurons. **(m,n)** For 934 neurons classified to be OS at 3.6-µm aFWHM, the cumulative distributions and histograms of their gOSI values; **(o, p)** Distributions of gOSI for their corresponding neuropil. Red numbers and blue lines: median gOSI values. **(q)** Color-coded preferred orientation (PO) maps for VR neurons in the example FOV in **(a)**. Top and middle rows: maps for cells classified to be OS and not OS at 3.6-µm aFWHM, respectively, color-coded by preferred orientation (gray for not OS). Bottom row: map for corresponding neuropil. **(r)** Leftmost histogram: distribution of preferred grating orientations of neurons classified to be OS at 3.6-µm aFWHM. Other histograms: distributions of changes in preferred orientation angles of these cells, relative to their preferred orientation angles at 3.6-µm aFWHM, at larger aFWHM values. Red numbers and blue lines: medians.

The loss of structural image quality was accompanied by a degradation of functional signals. In both single-trial and trial-averaged calcium movies (**Movie S1, S2**), the ability to discern individual active neurons was markedly reduced with increasing aFWHM. At 3.6-µm aFWHM, individual somata emerged as bright, sharply bounded cell bodies when activated by visual stimulation. At 5.7-µm aFWHM, the visibility of many cell bodies noticeably decreased. Above 10-µm aFWHM, the profiles of most somata became difficult to discern visually. At 21.0-µm aFWHM, most cells collapsed into a diffuse haze with little cellular definition.

#### Degradation of axial resolution leads to substantial errors in the functional characterization and classification of neurons

The qualitative loss of cellular contrast and signal-to-background ratio described above motivated a quantitative analysis of how reducing axial resolution affects calcium signals from the same neurons across aFWHM conditions (**Methods**). ROIs corresponding to neuronal somata were manually segmented from the reference image acquired at 3.6-µm aFWHM. The same ROI masks were then applied to datasets acquired at larger aFWHM values. For each ROI, we extracted the fluorescence trace and its corresponding neuropil signal at all five aFWHM values, and then performed neuropil subtraction with α = 0.7. Even for neurons whose structural and functional contrast was preserved across conditions (ROI1; F_raw_, F_ROI_, **Figure 2b**), calcium transient magnitudes decreased at larger aFWHM (ROI 1, Cell; trial-averaged ΔF/F_0_, **Figure 2c**). For neurons that became less visible at larger aFWHMs, trial-averaged calcium transients were severely attenuated when aFWHMs exceeded 10.2 µm (ROIs 2 and 3; F_raw_, F_ROI_, **Figure 2b**; Cell, trial-averaged ΔF/F_0_, **Figure 2c**).

We further characterized how increasing aFWHM impacted the functional classification of V1 neurons. We defined visually responsive (VR) neurons as those having significantly higher responses to drifting gratings than to dark screen (Student’s t-test, p < 0.05). We defined orientation-selective (OS) neurons as those having visual responses that varied significantly across the 12 drifting gratings (one-way ANOVA, p < 0.05), with their orientation tuning curves represented by the normalized trial-and time-averaged ΔF/F_0_ values at each grating orientation (Cell, normalized ΔF/F_0_, **Figure 2d**).

At 3.6-µm aFWHM, 85% of cells were classified as VR (2,122 out of 2,499 cells, 4 mice, 6 FOVs). As aFWHM increased, the percentage of VR ROIs decreased only moderately (gray crosses, **Figure 2e**), reaching 78% at 21.0-µm aFWHM. However, misclassification of visual responsiveness can include both false negatives, in which truly VR neurons appear unresponsive, and false positives, in which truly unresponsive neurons appear responsive. To distinguish these, we used the VR classification from the 3.6-µm aFWHM dataset as ground truth. At 21.0 µm aFWHM, the percentage of true VR neurons (i.e., those classified as VR at 3.6-µm aFWHM) that were correctly classified as VR decreased to 68% (gray dots, **Figure 2e**). Therefore, increasing the aFWHM from 3.6 µm to 21.0 µm resulted in 17% (85%-68%) of neurons misclassified as non-responsive (false negatives) and 10% (78%-68%) of neurons misclassified as VR (false positives). Overall, 27% of ROIs were misclassified in terms of visual responsiveness at the lowest resolution (gray triangles, **Figure 2f**).

The impact on classification of orientation selectivity, an important feature of visual cortical neurons^82^, was even more pronounced. At 3.6-µm aFWHM, 37% of ROIs were classified as OS (934 OS cells), while 29% of ROIs were classified as OS at 21.0-µm aFWHM (black crosses, **Figure 2e**). However, the percentage of true OS neurons (those classified as OS at 3.6-µm aFWHM) decreased to 15% at 21.0-µm aFWHM (black dots, **Figure 2e**). Therefore, increase of aFWHM from 3.6 to 21.0 µm resulted in 36% of ROIs having erroneous OS classification (black triangles, **Figure 2f**): 22% (37%-15%) of ROIs were falsely determined to be not OS, while 14% (29%-15%) of ROIs were mistakenly determined to be OS.

#### Degradation of axial resolution artifactually decreases calcium transient magnitudes

The misclassification at large aFWHMs was not driven by most ROIs becoming dimmer. Across the population (2,499 cells, 4 mice, 6 FOVs), average baseline brightness F_0, ROI_ increased significantly at larger aFWHMs (**Figure 2g**; normalized to the trial-averaged F_0,ROI_ at 3.6-µm aFWHM; see **Table S4** for statistical tests and p values). Under each aFWHM condition, the ROIs that were misclassified had distributions of baseline brightness F_0,ROI_ similar to those of correctly classified ROIs (**Figure S1a**), indicating that the misclassification did not arise from these neurons being dimmer or having noisier traces.

At 13.4- and 21.0-µm aFWHM, the mean baseline F_0,ROI_ remained higher than at 3.6-µm aFWHM. This differs from the bead experiments (c.f., **Figure 1b**), in which 10- or 15-µm-diameter beads were substantially dimmer at largest aFWHMs than at 3.6-µm aFWHM. Unlike isolated beads, which have no fluorescent neighbors above or below them, densely labeled brain tissue contains fluorescent structures above and below the cell body of interest, such as neuropil or the somata of other neurons. At larger aFWHMs, these structures were increasingly excited (e.g., **Figure 1d**) and contributed to the overall measured fluorescence signal. Because their fluorescence also varies with neuronal activity, they can contaminate responses from target cell bodies if not properly removed.

ΔF_ROI_ at larger aFWHMs also originates from fluorophores both inside and outside the cell body of interest. However, fluorescence changes associated with calcium activity from cell bodies typically have much larger magnitudes than those from neuropil (**Figure 1f**, ΔF_ROI_ traces in **Figure 2b**). Exciting neuropil at larger aFWHMs adds less to ΔF_ROI_ than to F_0, ROI_. As a result, ΔF_ROI_ increased more modestly with aFWHM than did F_0,ROI_ (**Figure 2h**). As a result, the calcium transient magnitude ΔF/F_0_ should decrease at large aFWHMs.

Indeed, for the 2,122 VR neurons identified at 3.6-µm aFWHM, maximal trial-averaged visually evoked ΔF/F_0_ (e.g., maxima of traces in **Figure 2c**) showed a monotonic decrease with increasing aFWHM (**Figure 2i; Table S5**). From 3.6- to 5.7-µm aFWHM, the median ΔF/F_0_ dropped by half; at 21.0-µm aFWHM, the median ΔF/F_0_ became 3.2-fold smaller. These population trends were also evident in the ΔF/F_0_ traces of the example ROIs (Cell responses, **Figure 2c**): whereas ROI 1’s ΔF/F_0_ reduction was moderate and its visual responsiveness was preserved, ROIs 2 and 3, whose cell bodies became obscured by blood vessel shadows or surrounding fluorescence at large aFWHMs, showed a precipitous drop in their maximal ΔF/F_0_, resulting in ROI 2 no longer satisfying the criterion for visual responsiveness at 21.0-µm aFWHM. By reducing the visually-triggered calcium transient magnitudes, imaging at lower resolution caused false negatives in VR classification.

#### Degradation of axial resolution increases neuropil contamination

We carried out similar analyses for the neuropil masks surrounding individual ROIs, whose pixel values arose from both neuronal processes and, at large aFWHMs, cell bodies that had been outside the excitation focus at 3.6-µm aFWHM. Consistent with a previous study reporting that neuropil exhibits visually evoked activity^72^, at 3.6-µm aFWHM, 99% of the neuropil mask regions exhibited visually evoked activity, with 34% exhibiting orientation selectivity.

In contrast to ROI masks, F_0, neuropil_ and ΔF_neuropil_ increased similarly at larger aFWHMs (**Figure 2j, k; Figure S1b**). Consequently, neuropil ΔF/F_0_ distributions across aFWHM conditions had similar median values (**Figure 2l, Table S5**). Reflecting the population trends, the neuropil surrounding ROI 1-4 had similar trial-averaged visually evoked calcium transients across aFWHMs (Neuropil, trial-averaged ΔF/F_0_, **Figure 2c**).

By geometry alone, at larger aFWHMs, the neuropil response constitutes a larger fraction of the activity measured from each ROI (e.g., **Figure 1d**). With the ROI ΔF/F_0_ decreasing with aFWHM while the neuropil mask ΔF/F_0_ remained largely constant (**Figure 2i,l**), imaging at larger aFWHMs exacerbates neuropil contamination.

#### Neuropil subtraction with ⍺ = 0.7 is insufficient for neuropil contamination removal at large aFWHMs and leads to errors in orientation-selectivity characterization

To accurately characterize the response properties of ROIs, contaminating neuropil activity needs to be fully removed from the ROI signals. However, as shown below, neuropil subtraction with ⍺ = 0.7 did not fully eliminate neuropil contamination, and the residual neuropil signal degraded the accuracy of orientation tuning characterization in V1 neurons.

For the 934 OS neurons identified at 3.6-µm aFWHM, the global orientation-selectivity index (gOSI) distribution (**Figure 2m,n**) was comparable to those reported in previous electrophysiology studies^83,84^ and high-resolution imaging studies^85–87^, with a median value of 0.49. At larger aFWHMs, however, the gOSI distributions shifted markedly toward smaller values (e.g., from a median of 0.49 at 3.6-µm to 0.40 at 21.0-µm aFWHM; **Figure 2m,n**; **Table S5**), indicating that neuropil subtraction with ⍺ = 0.7 does not fully remove neuropil contamination. Because the neuropil reflects the averaged activity of cellular processes from many neurons and is therefore more broadly tuned than cell bodies (**Figure 2o,p)**, residual neuropil contamination reduces the tuning of the somatic signal, lowering the apparent gOSI values of OS neurons. Consequently, uncorrected neuropil contamination led to errors in determining both the fraction of OS neurons (black crosses and dots, **Figure 2e,f**) and their orientation selectivity.

We observed a shift of the neuropil gOSI distribution toward higher values at 13.4- and 21.0-µm aFWHMs (**Figure 2p, Table S5**). This increase likely reflects the larger excitation volume at these aFWHMs, which can encompass dendritic or axonal branches, or even cell bodies, of nearby OS neurons, thereby making the neuropil response more tuned. Such tuned neuropil contamination can broaden tuning curves or shift an ROI’s preferred orientation toward that of the surrounding neuropil (e.g., ROI 3, **Figure 2d**; dashed red line indicates the preferred orientation at 3.6 µm). It can also cause non-VR neurons or non-OS cells at 3.6-µm aFWHM to artifactually acquire orientation selectivity: for example, ROI 4, a cell that was not OS at 3.6-µm aFWHM, was misclassified as OS at 10.2- and 21.0-µm aFWHMs.

These misclassifications can be visualized in the preferred orientation (PO) maps of an example FOV (top two rows, **Figure 2q**; for all FOVs, see **Figure S2**), which color-code ROIs based on their response properties: gray for non-VR or non-OS cells; colors for OS neurons, with the colors encoding the preferred orientation. Here, the preferred orientation (e.g., dashed red lines, **Figure 2d**) was determined by the vector sum of ROI responses across different grating directions^88^ (**Methods**). We similarly determined PO maps for the neuropil masks (color-coded in the corresponding ROI masks for easier visualization; bottom row, **Figure 2q**).

At 3.6-µm aFWHM, the PO map of OS neurons shows a characteristic “salt-and-pepper” organization^6^, while the neuropil PO maps reveal more spatially correlated, patch-like domains, consistent with a previous report^72^. As aFWHM increased, more OS neurons became non-OS (top row, gray ROIs, **Figure 2q**), whereas some non-OS or non-VR neurons were misclassified as OS (middle row, colored ROIs, **Figure 2q**). These spurious OS neurons tended to exhibit a preferred orientation similar to the local neuropil preference (bottom row, **Figure 2q**, **Figure S2**), consistent with these misclassifications being artifacts of neuropil contamination.

We examined how the POs of individual neurons that were classified as OS at 3.6-µm aFWHM (PO distribution, gray histogram, **Figure 2r**) varied with resolution degradation, by calculating the absolute changes in their PO angles (|ΔPO|, blue histograms, **Figure 2r**). When aFWHM increased to 5.7 µm, the changes in PO angles had a median of 11.1°; with further increases in aFWHM, the median continued to grow, reaching 24.8° at 21.0-µm aFWHM, indicating larger errors at larger aFWHMs.

Collectively, these findings demonstrate that reducing 2PFM spatial resolution compromises the accuracy of neuronal activity measurements by degrading somatic signal quality and increasing neuropil contamination. They further indicate that standard neuropil subtraction with ⍺ = 0.7 does not adequately remove neuropil contamination, resulting in widespread misclassification of neuronal functional properties.

### V1 neurons with transgenic pan-excitatory GCaMP6s expression suffer from similar artifacts caused by reduced axial resolution

In the experiments described above, GCaMP expression was transduced via intracortical virus injection, resulting primarily in labeling of neurons and neuropil near the injection site. With the development of recombinase-based driver and reporter lines^89,90^, calcium imaging experiments are increasingly performed in transgenic mice with pan-excitatory or pan-neuronal GCaMP expression, which produces strong neuropil fluorescence. To examine how this widespread sensor expression affects calcium imaging at reduced spatial resolution, we performed the same experiments and analyses in transgenic mice expressing GCaMP6s in excitatory neurons (Slc17a7-IRES2-Cre × Ai162D, **Table S4, Movie S3**).

As in the viral preparation, time-averaged images showed a progressive loss of spatial resolution and contrast with increasing aFWHM (**Figure 3a**). Likewise, the ability to visually identify individual active neurons in single-trial and trial-averaged calcium movies decreased rapidly as aFWHM increased (**Movie S3, S4**). Reduced resolution resulted in similar functional effects at the level of individual neurons (example fluorescence and activity traces, **Figure 3b,c**): some neurons retained their activity traces across aFWHMs (e.g., ROI1, **Figure 3a-d**); others became obscured by blood vessel shadows (e.g., ROI 2, **Figure 3a-d**); and many suffered from artifacts such as shifts in tuning preference (e.g., ROI 3, **Figure 3a-d**) or spurious orientation selectivity (e.g., ROI 4, **Figure 3a-d**).

**Figure 3.**
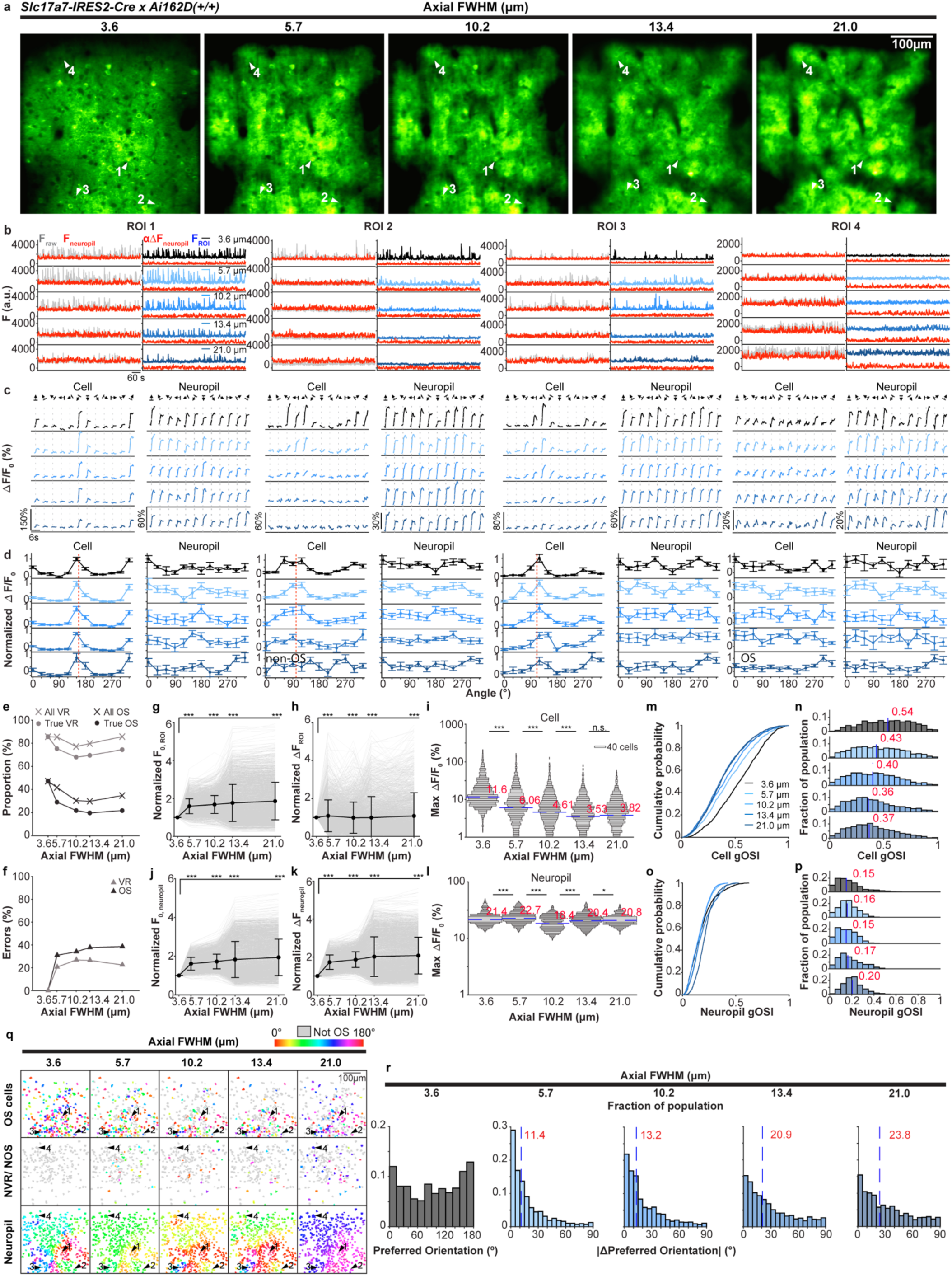
Degradation in axial resolution leads to similar errors for neurons that transgenically express GCaMP6s in the cytosol. **(a)** 2P fluorescence images of an example FOV of V1 L2/3 neurons of a transgenic mouse (Slc171a7-IRES2-Cre x Ai162D (+/+)) acquired at five aFWHM values of 3.6, 5.7, 10.2, 13.4, and 21.0 µm. Arrowheads, example neurons: Cell 1 is classified as VR and OS under all conditions; Cell 2 is misclassified as not OS as it moves into the shadow of the blood vessel at large aFWHMs; Cell 3 exhibits changes in tuning curve and a shift in preferred orientation at large aFWHMs; Cell 4 is misclassified as OS as aFWHM elongates. **(b)** Representative temporal traces of F_raw_ (gray), F_neuropil_ (red), αΔF_neuropil_ (red), and F_ROI_ (blue to black, color-coded by aFWHM) for Cell 1-4. α= 0.7. **(c)** Trial-averaged ΔF/F_0_ traces for Cell 1-4 and their neuropil. Gray dotted lines: onset of grating stimuli. **(d)** Tuning curves based on trial- and time-averaged ΔF/F_0_ during grating presentation for Cell 1-4 and their neuropil. Normalized individually to maximal ΔF/F_0_ values. Red dashed lines: preferred grating angles. **(e)** Out of 2,452 neurons, fraction of cells classified to be VR (gray cross) or OS (black cross) from images acquired with different aFWHM values; fraction of cells that are truly VR (gray dot) or OS (black dot). **(f)** Fraction of neurons that are misclassified in terms of VR (gray triangle) and OS (black triangle) properties. **(g)** F_0, ROI_ and **(h)** ΔF_ROI_ of 2,366 neurons for all aFWHM conditions, normalized to the value at 3.6 µm aFWHM (gray lines). Black dots: mean; error bars: s.d. Two-sample KS test: ***p < 0.001. **(i)** For 2,101 neurons classified to be VR at 3.6-µm aFWHM, the distributions of their maximal trial-averaged ΔF/F_0_ across conditions. Red numbers and blue lines: median values. One-sided WRS test against previous aFWHM conditions: ***p < 0.001. **(j)** F_0, neuropil_ and **(k)** ΔF_neuropil_ for neuropil associated with the 2,452 neurons. **(l)** Distributions of maximal trial-averaged ΔF/F_0_ for neuropil associated with 2,101 VR neurons. *p < 0.05, ***p < 0.001. **(m,n)** For 1,124 neurons classified to be OS at 3.6-µm aFWHM, the cumulative distributions and histograms of their gOSI values; **(o, p)** Distributions of gOSI for their corresponding neuropil. Red numbers and blue lines: median gOSI values. **(q)** Color-coded preferred orientation (PO) maps for VR neurons in the example FOV in **(a).** Top and middle rows: maps for cells classified to be OS and not OS at 3.6-µm aFWHM, respectively, color-coded by preferred orientation (gray for not OS). Bottom row: map for corresponding neuropil. **(r)** Leftmost histogram: distribution of preferred grating orientations of neurons classified to be OS at 3.6-µm aFWHM. Other histograms: distributions of changes in preferred orientation angles of these cells, relative to their preferred orientation angles at 3.6-µm aFWHM, at larger aFWHM values. Red numbers and blue lines: medians.

At the population level (3 mice, 6 FOVs, 2,452 cells; 2,101 VR and 1,124 OS cells at 3.6-µm aFWHM), many neurons were misclassified in datasets acquired at larger aFWHMs (**Figure 3e,f**). Under each aFWHM condition, the incorrectly classified ROIs had distributions of baseline brightness F_0,ROI_ similar to those of correctly classified ROIs (**Figure S1c**), indicating that these classification errors did not result from these neurons having noisier traces. Instead, as in the viral preparation, the misclassification of VR neurons arose because increases in aFWHM drove larger rises in F_0, ROI_ than in ΔF_ROI_ (**Figure 3g,h**; **Table S5**), leading to a drop in somatic ΔF/F_0_ **(Figure 3i**; **Table S5)**.

As in virally transduced preparations, F_0, neuropil_ and ΔF_neuropil_ in transgenic mice increased to a similar extent with aFWHM **(Figure 3j,k; Figure S1d**). Consequently, neuropil ΔF/F_0_ remained largely unchanged with aFWHM (**Figure 3l**). As somatic ΔF/F_0_ decreased with aFWHM, inadequate neuropil contamination removal at large aFWHMs led to a reduction in the orientation selectivity of the population, with smaller median gOSI values at large aFWHMs (e.g., medians of 0.54 at 3.6-μm and 0.37 at 21.0-μm aFWHM, respectively; **Figure 3m,n**). This reduction was more pronounced than in the viral preparation (cf., **Figure 2m,n**), likely because this transgenic mouse line expresses the sensor in the axons and dendrites of all excitatory neurons. Since the neuropil signal represents the averaged response across many more excitatory neurons, it exhibited lower orientation selectivity than viral expression, where only a subset of neurons was labeled (**Figure 3o,p**; cf., **Figure 2o,p**). At 21.0-μm aFWHM, the axially expanded focal volume captured more dendritic trees or somata of OS neurons, causing neuropil responses to become more orientation tuned and leading to an increase in gOSI (**Figure 3o,p**). At lower resolutions, inadequate neuropil contamination removal resulted in shifts in the preferred orientation of OS neurons toward that of the neuropil, as visualized in the PO maps (**Figure 3q**_;_ all FOVs, **Figure S3**), and an increase of PO angle change for OS neurons (**Figure 3r).**

In summary, transgenic pan-excitatory GCaMP6s expression exhibited resolution-dependent artifacts similar to, and in some respects more pronounced than those observed with virally transduced GCaMP6s expression.

### Soma-targeted expression of jGCaMP8s remains susceptible to artifacts at reduced axial resolution

Strategies have been developed to restrict GCaMP expression to neuronal somata to reduce neuropil contamination^80,91,92^. We next investigated whether restricting sensor expression primarily to neuronal somata could confer resilience to artifacts caused by low-resolution imaging, by conducting the same experiments and analyses using a soma-targeted calcium sensor expressed in V1 L2/3 neurons via intracortical viral injection of *AAV2/PHP.eB-hSyn-RiboL1-jGCaMP8s* (**Table S4, Movie S5**)^80^.

As expected, fluorophore expression in the neuropil was reduced compared to cytosolic sensors. At high resolution, time-averaged images and movies revealed mostly isolated somata (**Figure 4a, Movie S6**). At 3.6-µm aFWHM (3 mice, 7 FOVs, N = 2,810 cells), the ratio of basal fluorescence between neuropil and soma of each ROI, F_0, Neuropil_/F_0, ROI_, was 0.55 ± 0.21 (mean ± s.d.) for RiboL1-jGCaMP8s, smaller than the values of 0.89 ± 0.18 and 0.92 ± 0.15 for virally transduced and transgenically expressed cytosolic GCaMP6s, respectively. However, neuropil fluorescence and their calcium transients remained detectable (**Figure 4b,c**) and thus could still cause neuropil contamination. Furthermore, after normalization by excitation power, the averaged fluorescence brightness of jGCaMP8s-expressing cell bodies was 10.0- and 9.7-fold smaller than that of virally transduced and transgenically expressed GCaMP6s, respectively. Our observations were consistent with previous reports that soma-targeting yields lower sensor expression in somata and does not fully prevent neuropil expression^80,91,92^.

**Figure 4.**
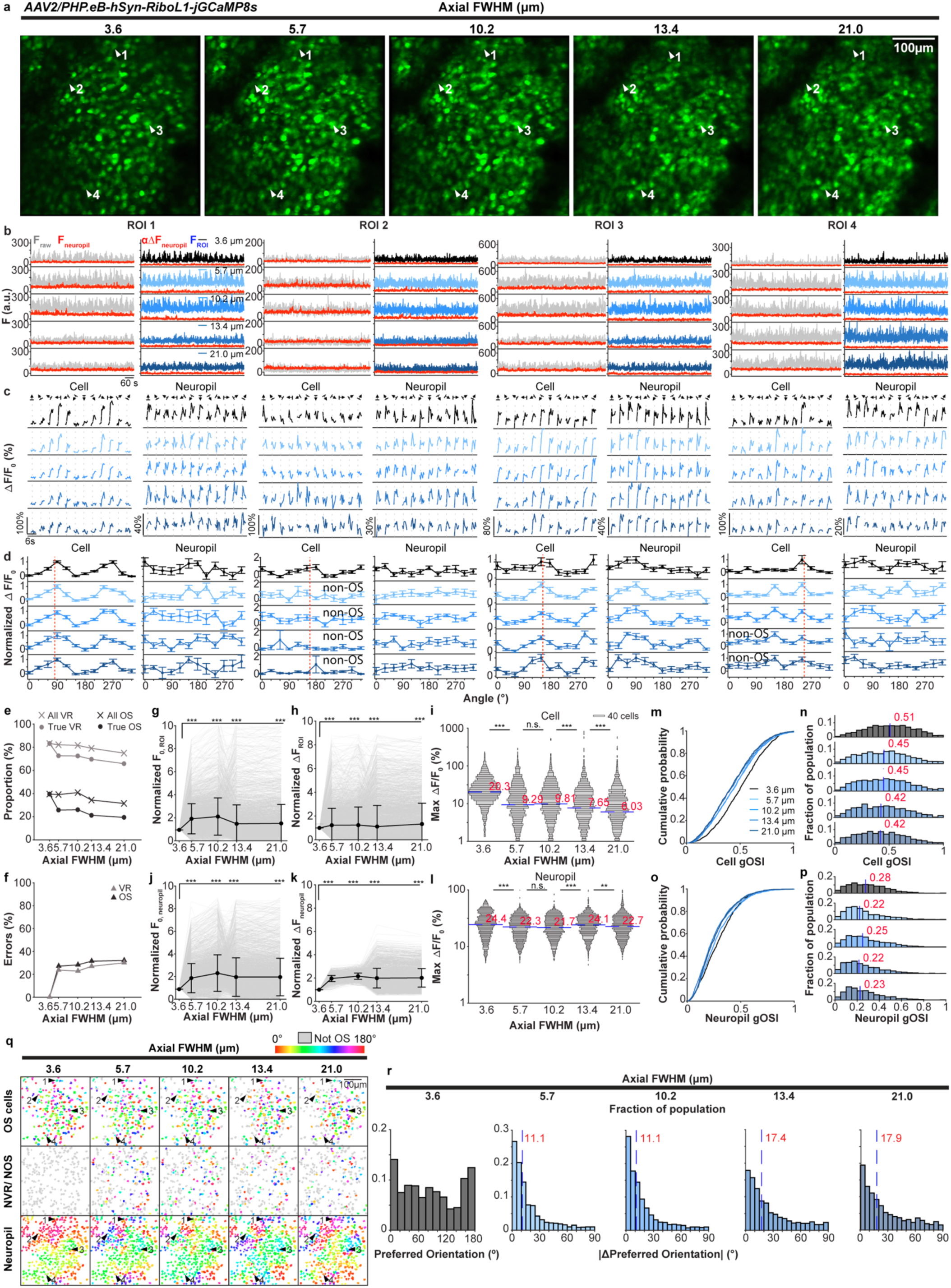
Soma-targeted expression of jGCaMP8s remains susceptible to artifacts at reduced axial resolution. **(a)** 2P fluorescence images of an example FOV of mouse V1 L2/3 neurons labeled with AAV-hSyn-RiboL1-jGCaMP8s acquired at five aFWHM values of 3.6, 5.7, 10.2, 13.4, and 21.0 µm. Arrowheads, example neurons: Cell 1 is classified as VR and OS under all conditions; Cell 2 is misclassified as not VR as it becomes obscured by a blood vessel shadow at large aFWHMs; Cell 3 exhibits changes in tuning curve and a shift in preferred orientation at large aFWHMs; Cell 4 exhibits changes in tuning curve and is misclassified as not OS as aFWHM elongates. **(b)** Representative temporal traces of F_raw_ (gray), F_neuropil_ (red), αΔF_neuropil_ (red), and F_ROI_ (blue to black, color-coded by aFWHM) for Cell 1-4. α = 0.7. **(c)** Trial-averaged ΔF/F_0_ traces for Cell 1-4 and their neuropil. Gray dotted lines: onset of grating stimuli. **(d)** Tuning curves based on trial- and time-averaged ΔF/F_0_ during grating presentation for Cell 1-4 and their neuropil. Normalized individually to maximal ΔF/F_0_ values. Red dashed lines: preferred grating angles. **(e)** Out of 2,810 neurons, fraction of cells classified to be VR (gray cross) or OS (black cross) from images acquired with different aFWHM values; fraction of cells that are truly VR (gray dot) or OS (black dot). **(f)** Fraction of neurons that are misclassified in terms of VR (gray triangle) and OS (black triangle) properties. **(g)** F_0_, _ROI_ and **(h)** ΔF_ROI_ of 2,810 neurons for all aFWHM conditions, normalized to the value at 3.6 µm aFWHM (gray lines). Black dots: mean; error bars: s.d. Two-sample KS test: ***p < 0.001. **(i)** For 2,341 neurons classified to be VR at 3.6-µm aFWHM, the distributions of their maximal trial-averaged ΔF/F_0_ across conditions. Red numbers and blue lines: median values. One-sided WRS test against previous aFWHM conditions: ***p < 0.001. **(j)** F0, neuropil and **(k)** ΔF_neuropil_ of neuropil associated with the 2,810 neurons. (l) Distributions of maximal trial-averaged ΔF/F_0_ for neuropil associated with 2,341 VR neurons. ***p < 0.001. (m,n) For 1,040 neurons classified to be OS at 3.6-µm aFWHM, the cumulative distributions and histograms of their gOSI values; **(o,p)** Distributions of gOSI for their corresponding neuropil. Red numbers and blue lines: median gOSI values. **(q)** Color-coded preferred orientation (PO) maps for VR neurons in the example FOV in **(a).** Top and middle rows: maps for cells classified to be OS and not OS at 3.6-µm aFWHM, respectively, color-coded by preferred orientation (gray for not OS). Bottom row: map for corresponding neuropil. **(r)** Leftmost histogram: distribution of preferred orientations of neurons classified to be OS at 3.6-µm aFWHM. Other histograms: distributions of changes in preferred orientation angles of these cells, relative to their preferred orientation angles at 3.6-µm aFWHM, at larger aFWHM values. Red numbers and blue lines: medians.

At large aFWHMs, somata at different axial locations started to overlap (**Figure 4a**). As a purely optical effect caused by the elongation of the excitation volume, somatic overlap also occurred with cytosolic sensors but was obscured by the strong neuropil fluorescence. Here, soma-targeted expression made the effects of overlapping somata on the accuracy of calcium imaging more readily detectable.

As with cytosolic sensors, some neurons remained mostly unchanged in their tuning properties across aFWHM conditions (e.g., ROI 1; **Figure 4a-d**), while others were obscured by blood vessel shadows (e.g., ROI 2, an OS neuron that was misclassified as non-OS at large aFWHMs; **Figure 4a-d**). As expected, we observed altered tuning properties for neurons at large aFWHMs (e.g., a change in preferred orientation, ROI 3; a loss of orientation selectivity, ROI 2,4; **Figure 4a-d**).

At the population level, residual neuropil expression and somata crosstalk caused the fractions of misclassified VR (2,341 VR cells at 3.6-µm aFWHM) and OS neurons (1,040 OS cells at 3.6-µm aFWHM) to increase with aFWHM, reaching levels comparable to those observed with cytosolic sensors (**Figure 4e,f**). Correctly and incorrectly classified neurons, and their surrounding neuropil, had similar baseline brightness distributions (**Figure S1e,f**), indicating that these errors did not arise from the dimmer fluorescence of the soma-targeted sensor. Consistent with previous observations, increasing aFWHM led to a decrease in the maximum trial-averaged somatic ΔF/F_0_ and median gOSI for ROIs (**Figure 4g,h,i,m,n**), while neuropil ΔF/F_0_ remained largely unchanged (**Figure 4j,k,l,o,p**). However, soma-targeted sensors showed some improvement over cytosolic sensors: PO maps revealed a weaker correlation between the preferred orientations of spuriously classified OS neurons and their surrounding neuropil (**Figure 4q**; all FOVs, **Figure S4**). The changes in PO angles at large aFWHMs were also less pronounced than those observed with cytosolic sensors (median |ΔPO| of 17.9° at 21.0 µm aFWHM, **Figure 4r**; *cf.* 24.8° in **Figure 2r** and 23.8° in **Figure 3r**), indicating a moderate improvement attributable to reduced neuropil contamination.

In summary, soma-targeted jGCaMP8s has much lower brightness and does not fully mitigate the artifacts that affect cytosolic sensors at low spatial resolution. Crosstalk from cells that were out of focus at 3.6-µm aFWHM, together with residual contamination from neuropil still led to errors in the functional characterization of neurons when resolution was compromised.

### Functional classification errors from low axial resolution persist across analysis pipelines

Up to this point, our analyses were based on hand-segmented ROIs with neuropil subtraction at a fixed ⍺ = 0.7, which does not adequately remove neuropil contamination. We therefore evaluated whether alternative methods could better compensate for the loss of axial resolution, including: (1) cell-specific ⍺ neuropil subtraction (i.e., “Variable ⍺” and “Allen SDK”), (2) automatic ROI detection with neuropil subtraction (i.e., Suite2p with its default ⍺ = 0.7), and (3) automated ROI detection with demixing of overlapping activity sources (CNMF). Calcium imaging datasets acquired at different aFWHM conditions for each FOV were processed independently by Suite2p (Python implementation, version 0.13.2.dev19) and CNMF (Python package in CaImAn^66^, version 1.9.11). For both methods, we systematically varied key parameters (“threshold_scaling” for Suite2p, “min_SNR” and “nlter” for CNMF) using ranges employed in the literature^93–96^ on an example FOV with transgenic GCaMP6s expression acquired at 3.6-µm aFWHM. We observed only minor changes in the numbers of ROIs detected and the maximal trial-averaged ΔF/F_0_ values (**Figure S5**). Therefore, default parameters were used unless otherwise noted (**Methods**). With these pipelines and parameter settings, we then asked whether any of these analysis pipelines could mitigate the resolution-dependent classification errors documented above.

#### SNR thresholding and advanced neuropil subtraction do not correct artifacts from reduced axial resolution

To enable fair comparisons with Suite2p and CNMF, both of which apply SNR thresholding to identify valid ROIs, we rejected hand-segmented ROIs with SNR ≤ 3.5 for the “⍺ = 0.7”, “Variable ⍺”, and “Allen SDK” analyses below (**Methods**). For the “⍺ = 0.7” analysis, SNR thresholding preferentially removed cells with smaller calcium transients (**Figure S6a,b**) and reduced the number of ROIs by up to 17% for the 3.6-µm datasets and by up to 26% for the 21.0-µm datasets (**Table 1**). As a result, the medians of the max ΔF/F_0_ distributions increased after thresholding across all aFWHM and labeling conditions (**Figure S6c,d**, **Table S6,7**). The median values rose by up to 19% for the 3.6-µm datasets, whereas increases of up to 3.5-fold were observed for the 21.0-µm datasets, indicating substantial SNR degradation caused by lower-resolution image acquisition.

**Table 1.**
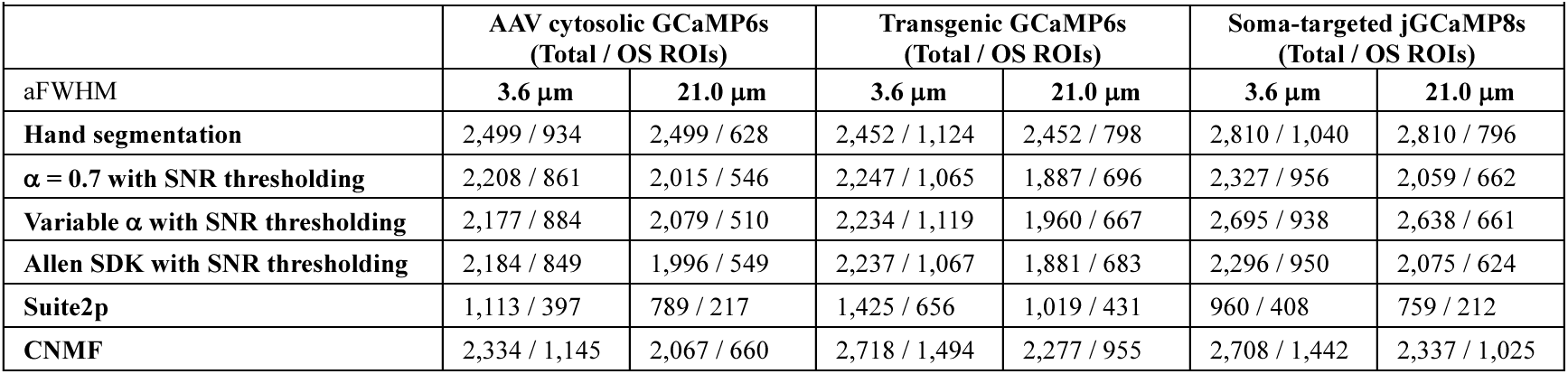
Total number of ROIs detected and the corresponding number of OS ROIs by different pipelines.

|  |  |  |  |  |  |  |
| --- | --- | --- | --- | --- | --- | --- |
|  | AAV cytosolic GCaMP6s<br>(Total / OS ROIs) |  | Transgenic GCaMP6s<br>(Total / OS ROIs) |  | Soma-targeted jGCaMP8s<br>(Total / OS ROIs) |  |
| aFWHM | 3.6 μm | 21.0 μm | 3.6 μm | 21.0 μm | 3.6 μm | 21.0 μm |
| Hand segmentation | 2,499 / 934 | 2,499 / 628 | 2,452 / 1,124 | 2,452 / 798 | 2,810 / 1,040 | 2,810 / 796 |
| $\alpha = 0.7$ with SNR thresholding | 2,208 / 861 | 2,015 / 546 | 2,247 / 1,065 | 1,887 / 696 | 2,327 / 956 | 2,059 / 662 |
| Variable $\alpha$ with SNR thresholding | 2,177 / 884 | 2,079 / 510 | 2,234 / 1,119 | 1,960 / 667 | 2,695 / 938 | 2,638 / 661 |
| Allen SDK with SNR thresholding | 2,184 / 849 | 1,996 / 549 | 2,237 / 1,067 | 1,881 / 683 | 2,296 / 950 | 2,075 / 624 |
| Suite2p | 1,113 / 397 | 789 / 217 | 1,425 / 656 | 1,019 / 431 | 960 / 408 | 759 / 212 |
| CNMF | 2,334 / 1,145 | 2,067 / 660 | 2,718 / 1,494 | 2,277 / 955 | 2,708 / 1,442 | 2,337 / 1,025 |

Despite removing ROIs with noisier traces, the fractions of ROIs erroneously classified in terms of visual responsiveness and orientation selectivity remained high (c.f., **Figure 5a,b**; **Methods**). This is consistent with the observation that correctly and incorrectly classified ROIs had similar baseline brightness (**Figure S1**) and thus indicates that SNR-based thresholding cannot effectively improve functional classification accuracy.

**Figure 5.**
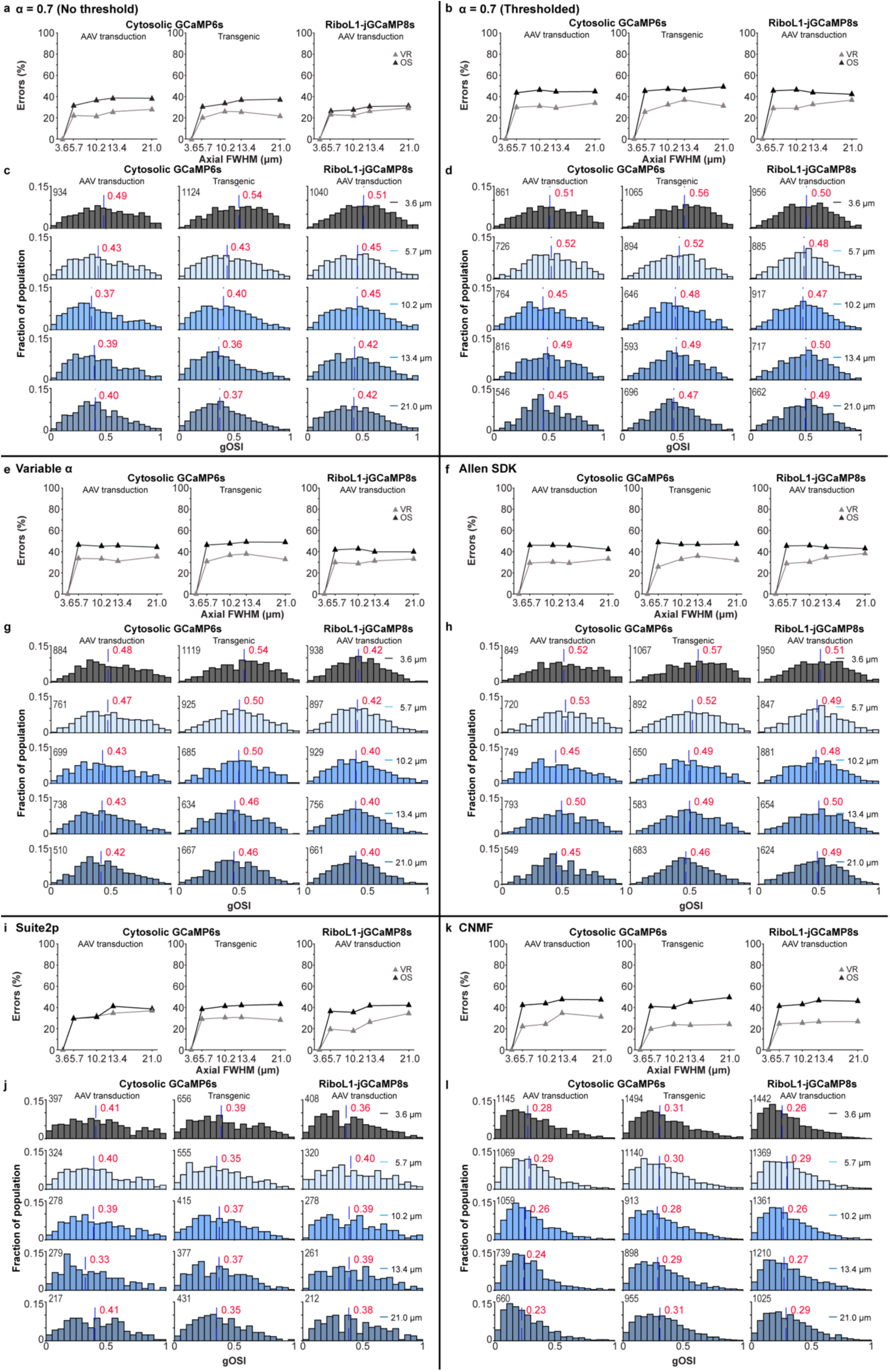
Classification and tuning characteristics errors persist across analysis pipelines. **(a)** Fraction of neurons that are misclassified in terms of VR (gray triangle) and OS (black triangle) properties for the “α = 0.7” subtraction method (same data as Figs. 2,3,4f). **(b)** Same as (a) but for neurons with activity traces that pass thresholding with SNR > 3.5. At 3.6-µm aFWHM: 1,876 VR and 861 OS neurons of 2,208 manually segmented cells expressing GCaMP6s via viral transduction; 1,934 VR and 1,065 OS of 2,247 manually segmented cells expressing GCaMP6s transgenically; 2,006 VR and 956 OS of 2,327 manually segmented cells expressing RiboL1-jGCaMP8s via viral transduction. **(c)** Histograms of gOSI for neurons classified as OS by “α = 0.7” method without SNR thresholding from the dataset acquired at 3.6-µm aFWHM (same data as Figs. 2,3,4n). Red numbers and blue dashed lines here and below: median values. Color-coded by aFWHM here and below: black to blue. Gray numbers near the top of the y axis here and below: number of OS neurons; values are the same across aFWHM conditions for this panel. **(d)** Histograms of gOSI for neurons in (c) but also with SNR > 3.5. **(e)** Fraction of neurons that are misclassified for “Variable α”. At 3.6-µm aFWHM: 1,783 VR and 884 OS of 2,177 manually segmented cells passing SNR thresholding and expressing GCaMP6s via viral transduction; 1,878 VR and 1,119 OS of 2,234 manually segmented cells passing SNR thresholding and expressing GCaMP6s transgenically; 2,263 VR and 938 OS of 2,695 manually segmented cells passing SNR thresholding and expressing RiboL1-jGCaMP8s via viral transduction. **(f)** Fraction of neurons that are misclassified for “Allen SDK”. At 3.6-µm aFWHM: 1,861 VR and 849 OS of 2,184 manually segmented cells passing SNR thresholding and expressing GCaMP6s via viral transduction; 1,926 VR and 1,067 OS of 2,237 manually segmented cells passing SNR thresholding and expressing GCaMP6s transgenically; Soma-targeted: 1,978 VR and 950 OS of 2,296 manually segmented cells passing SNR thresholding and expressing RiboL1-jGCaMP8s via viral transduction. **(g,h)** Histograms of gOSI for neurons classified as OS by “Variable α” and “Allen SDK” for datasets acquired at each aFWHM condition, respectively. **(i)** Fraction of neurons that are misclassified for Suite2p. At 3.6-µm aFWHM: 833 VR and 397 OS of 1,113 auto-detected cells expressing GCaMP6s via viral transduction; 1,138 VR and 656 OS of 1,425 auto-detected cells expressing GCaMP6s transgenically; 830 VR and 408 OS of 960 auto-detected cells expressing RiboL1-jGCaMP8s via viral transduction. **(j)** Histograms of gOSI for neurons classified as OS by Suite2p for datasets acquired at each aFWHM condition. **(k)** Fraction of neurons that are misclassified for CNMF. At 3.6-µm aFWHM: 1,998 VR and 1,145 OS of 2,334 auto-detected cells expressing GCaMP6s via viral transduction; 2,399 VR and 1,494 OS of 2,718 auto-detected cells expressing GCaMP6s transgenically; 2,334 VR and 1,442 OS of 2,708 auto-detected cells expressing RiboL1-jGCaMP8s via viral transduction. **(l)** Histograms of gOSI for neurons classified as OS by CNMF for datasets acquired at each aFWHM condition.

SNR thresholding did increase the gOSI values for OS ROIs under larger aFWHM conditions (c.f., **Figure 5c,d**; **Table S8**). However, gOSI still decreased for cytosolic sensors at lower resolution. For neurons with soma-targeted sensor, gOSI distributions were similar across all aFWHM conditions, with median values close to the ground truth. However, this improvement came at the cost of losing OS cells: from 3.6-µm to 21.0-µm aFWHM, the number of OS neurons was reduced by 30% to 37% (see cell numbers at the top left corners of the histograms in **Figure 5d**).

To examine whether cell-specific ⍺ values would improve performance, we applied the “Variable ⍺” and “Allen SDK” methods to the F_raw_ and F_neuropil_ traces extracted from the hand-segmented ROIs described above. Neuropil subtraction, F_ROI_(t)=F_raw_(t) - ⍺ΔF_neuropil_(t), was then performed for each ROI using its cell-specific ⍺ value, from which ΔF/F_0_ trace was calculated and ROIs with SNR ≤ 3.5 were excluded.

For virally transduced and transgenic GCaMP6s data, the “Variable ⍺” and “Allen SDK” methods returned median ⍺ values ranging from 0.82 to 0.93 (**Figure S6e,f**), resulting in stronger neuropil subtraction than with ⍺ = 0.7. However, the two methods produced max ΔF/F_0_ distributions similar to those of “⍺ = 0.7” (**Figure S6g,h**), and classification errors also persisted at similar levels (**Figure 5e,f**). A decrease in the gOSI median values for cytosolic sensors at large aFWHMs **(Figure 5g,h**) further indicates that neuropil contamination was not fully removed.

Overall, the three neuropil subtraction methods, when applied to hand-segmented ROIs with SNR thresholding, produced highly consistent results. They led to highly overlapping sets of ROIs (see the example FOV in **Figure S7a**), as well as comparable numbers of neurons passing the SNR threshold, similar VR and OS neuron numbers and fractions across all imaging and labeling conditions (**Figure S7b,c**), and similar gOSI and preferred orientation distributions (**Figure S7d,e**). For datasets acquired at lower resolution, all three methods yielded comparable, substantial errors in the functional classification and orientation-tuning characterization of V1 L2/3 neurons.

#### Suite2p and CNMF do not resolve artifacts from reduced axial resolution

Compared with hand segmentation, Suite2p detected substantially fewer ROIs, with counts approximately 30-50% of those obtained by hand segmentation (**Figure S7a,b**, **Table 1**), consistent with a previous report^97^. We used Suite2p’s default neuropil subtraction formula F_ROI_(t)=F_raw_(t) - 0.7 ∗ F_neuropil_(t). Subtracting F_neuropil_, instead of ΔF_neuropil_ as performed above on hand-segmented ROIs, yielded smaller F_0_ and consequently larger ΔF/F_0_ values (**Figure S6i**). Accordingly, the max ΔF/F_0_ values for VR cells were approximately double those of the hand-segmented ROIs (**Table S6**).

Despite these methodological differences, the fractions of VR and OS neurons from Suite2p analysis (**Figure S7c**), as well as the preferred orientation distributions of OS neurons (**Figure S7f**), closely tracked those from hand-segmented ROIs. Likewise, functional response characteristics derived from Suite2p analysis remained susceptible to artifacts in data acquired at reduced axial resolution. Classification errors for both VR and OS neurons remained substantial and mostly increased with aFWHM **(Figure 5i**).

The gOSI distributions showed much less dependence on aFWHM, with consistently similar median values (**Figure 5j**); However, with medians ranging from 0.33 to 0.42, they were much smaller than the ground-truth values of ∼0.50. At all aFWHM conditions, Suite2p’s gOSI distributions had smaller median values than those from hand-segmented ROI analysis. We speculate that these differences may arise from Suite2p weighting the pixel values in the ROI masks when calculating F_raw_(t).

CNMF detected similar numbers of ROIs to or more than hand segmentation (**Figure S7a,b**; **Table 1**), likely because it can identify spatially overlapping sources of activity. The ΔF/F_0_ values from CNMF analysis were closer to those obtained from hand-segmented ROIs than were those from Suite2p (**Figure S6j; Table S6**). As with all previous methods, classification errors for both VR and OS neurons remained high and worsened at larger aFWHMs **(Figure 5k**).

As with Suite2p, the gOSI distributions from CNMF showed less dependence on aFWHM. However, with medians ranging from 0.23 to 0.31, they were even lower than those from Suite2p, even for the highest-resolution data acquired at 3.6-µm aFWHM (**Figure S7e, Figure 5l, Table S8**). The reduction in gOSI associated with CNMF analysis has also been reported in previous publications^98,99^.

Taken together, these results demonstrate that the functional classification errors caused by reduced axial resolution persist across different labeling strategies and analysis pipelines. Despite the substantial methodological differences among the approaches evaluated – including SNR thresholding, cell-specific neuropil subtraction, automated ROI detection, and source demixing – all methods yielded comparable misclassification rates for VR and OS neurons that increased with aFWHM. While some approaches differed in the absolute number of detected ROIs, ΔF/F₀ magnitudes, or gOSI values, none corrected the underlying resolution-dependent artifacts. These findings indicate that the errors in classification and tuning characterization described here are an inherent consequence of reduced axial resolution rather than a limitation of any particular analysis methods, and that high axial resolution at the time of image acquisition remains the most effective means of preventing them.

### Low axial resolution leads to errors in orientation tuning correlation

Having established that imaging at low spatial resolution leads to errors in functional classification and tuning characterization of individual neurons that even advanced analysis pipelines cannot correct, we next examined how it affects measurements of functional correlation within cortical circuits – properties fundamental to understanding circuit organization.

Since the seminal calcium imaging experiment that reported a “salt-and-pepper” arrangement of orientation tuning preference in the rat V1, in contrast to the orientation columns observed in the cat V1^6^, calcium imaging has been extensively used to investigate activity correlation and population encoding in the rodent, and most commonly, the mouse V1. However, whether nearby neurons in the mouse V1 are more likely to share similar orientation preferences, a question that one would expect calcium imaging to answer definitely, remains unsettled. Whereas one study reported broad clustering with a correlation length over hundreds of micrometers^100^, another found narrow clustering limited to tens of micrometers^101^.

We speculated that these disparate results may have stemmed from differences in neuropil correction. Given that developing accurate cortical models requires knowing not only whether such clustering exists but also its precise spatial scale, in a separate study^102^, our group set out to resolve this controversy. We first acquired calcium imaging data from mouse V1 L2/3 neurons expressing GCaMP6 cytosolically at high spatial resolution. We hand-segmented the neurons and found that, by varying the neuropil subtraction coefficient ⍺ from 0 to 1, the spatial correlation of orientation tuning narrowed from hundreds to tens of micrometers. Given the lack of a principled way to perform neuropil contamination correction, we then expressed nuclear-targeted GCaMP6 (H2B-GCaMP6) in L2/3 neurons. With GCaMP6s restricted to the nucleus, neuropil contamination was completely eliminated (**Figure 6a**). We then determined the preferred orientation of these neurons (**Figure 6b**) and found a positive orientation tuning correlation for OS neurons within ∼20 µm, which decayed to a baseline of zero correlation at greater pairwise distances between somata **(Figure 6c**), confirming a fine-scale spatial correlation and micro-clustering of orientation tuning in the mouse V1^102^.

**Figure 6.**
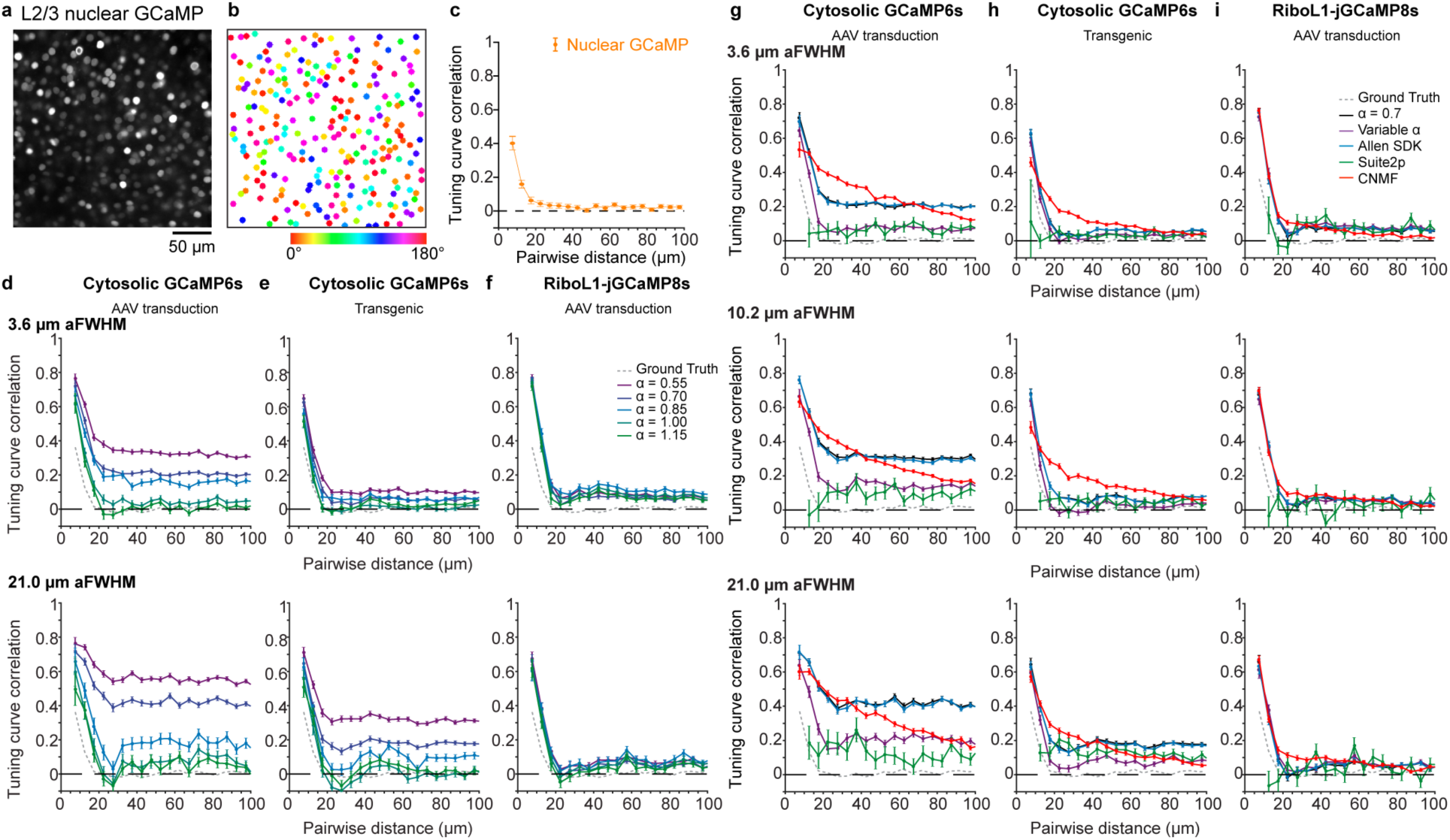
Accuracy of orientation tuning correlation over pairwise distance depends strongly on labeling strategy and analysis pipeline, with more artifacts observed at lower axial resolution. **(a)** An example FOV of V1 L2/3 neurons labeled with nuclear-targeted GCaMP6s and **(b)** its color-coded preferred orientation (PO) map. **(c)** Orientation tuning correlation for OS neuron pairs vs. distance for 582 OS neurons labeled with nuclear-targeted GCaMP6s from 2 mice, 7 FOVs. Data in (a-c) adapted from Ref. (104). **(d,e,f)** Orientation tuning correlations for manually segmented OS cells processed with neuropil subtraction at a constant α coefficients of 0.55, 0.7, 0.85, 1.0, or 1.15 for datasets acquired at 3.6-μm and 21.0-μm aFWHM for V1 L2/3 neurons with virally transduced GCaMP6s, transgenically expressed GCaMP6s, and virally transduced RiboL1-jGCaMP8s expression, respectively. Gray dotted traces: ground truth from (c). **(g,h,i)** Orientation tuning correlation for OS cells determined through “α = 0.7” (black), “Variable α” (purple), “Allen SDK” (blue), “Suite2p” (green), “CNMF” (red) for datasets acquired at 3.6-μm, 10.2-μm, and 21.0-μm aFWHM for V1 L2/3 neurons with virally transduced GCaMP6s, transgenically expressed GCaMP6s, and virally transduced RiboL1-jGCaMP8s expression, respectively. Error bars: s.e.m.

Having established the ground truth for the spatial scale of orientation tuning correlation, we calculated the population tuning curve correlation coefficient over pairwise distance for OS neurons from calcium imaging data acquired at different aFWHMs and analyzed by different pipelines (**Methods**).

For the hand-segmented ROIs of cytosolic data, we performed neuropil subtraction using constant ⍺ value of 0.55, 0.7, 0.85, 1.0, or 1.15 on datasets acquired at 3.6- or 21.0-µm aFWHM and applied SNR thresholding to the ΔF/F_0_ traces. We then identified OS neurons for each aFWHM condition and calculated their orientation tuning correlation with respect to pairwise distance between their somata. Depending on the ⍺ value used, we could reproduce either the long-range or short-range correlations (**Figure 6d,e**), consistent with our previous results^102^. The artifactual long-range correlations became more severe for datasets acquired at lower resolution (c.f., 3.6- and 21.0-µm aFWHM results in **Figure 6d,e**). Soma-targeted jGCaMP8s data showed much less long-range correlation even at 21.0 µm aFWHM (**Figure 6f**), suggesting that these long-range correlation artifacts indeed arose from neuropil contamination.

For AAV-transduced cytosolic expression of GCaMP6s, even at the high axial resolution of 3.6 µm aFWHM (top panel, **Figure 6g**), none of the analysis pipelines reproduced the ground truth from nuclear GCaMP6s. For hand-segmented ROIs, “⍺ = 0.7” and “Allen SDK” generated highly similar results, with high correlation at distances < 20 µm but an artifactual correlation coefficient of ∼0.2 that persisted over 100 µm; “Variable ⍺” produced a correlation curve most closely resembling the ground truth, yet still exhibited a nonzero baseline correlation of 0.06. Suite2p yielded a similarly small baseline correlation but failed to capture the high correlation at distances < 20 µm, likely due to the lower density of automatically detected ROIs (approximately half that of the other methods; **Table 1**).

Consistent with CNMF producing gOSI distributions that deviated most from the known properties of L2/3 neurons, CNMF yielded a highly artifactual trace characterized by slowly decreasing positive correlations over distance. Analysis on data acquired at lower axial resolution revealed more severe errors, with heightened artifactual correlations (middle and bottom panels for 10.2- and 21.0-µm aFWHM, respectively, **Figure 6g**; **Figure S8a**): compared with the high-resolution data, the erroneous correlations at large cortical distances increased by 50% at 10.2-µm aFWHM and doubled at 21.0-µm aFWHM for methods using neuropil subtraction; CNMF exhibited a smaller relative increase, though its correlation trace was already highly artifactual at the highest resolution.

All the hand-segmented ROI methods performed similarly well for transgenically expressed GCaMP6s data acquired at 3.6 µm aFWHM (top panel, **Figure 6h**), with artifactual correlations at larger distances approaching zero. As above, Suite2p produced low correlations over large distances but missed the high correlation at distances < 20 µm. For these methods, as aFWHM increased, tuning curve correlations had progressively elevated baselines (middle and bottom panels for 10.2- and 21.0-µm aFWHM, respectively, **Figure 6h**; **Figure S8b**). For these transgenic datasets, CNMF produced smaller artifactual correlation coefficients at larger pairwise distances than for the AAV-transduced datasets, but still deviated the most from the ground truth.

The best performance was, unsurprisingly, observed with the soma-targeted jGCaMP8s data (**Figure 6i**; **Figure S8c**). Except for Suite2p, which failed to capture the short-range correlation from dataset acquired at 10.2- and 21.0-μm aFWHMs, all other methods accurately captured the clustering of neurons with similar orientation preferences at distances < 20 µm; correlation coefficients at large distances, although non-zero, remained below 0.1. CNMF overestimated correlations at distances between 20 and 35 µm compared with other methods, but overall performed much better than with cytosolic data.

In summary, across all datasets, the accuracy of orientation tuning correlation coefficients over distance depended strongly on both imaging resolution and analysis pipeline. Artifactual long-range correlations were most severe for cytosolic GCaMP6s data and worsened at lower axial resolutions, with CNMF consistently producing the most artifactual results. Suite2p minimized spurious long-range correlations but failed to capture genuine short-range clustering across all conditions. Transgenic expression modestly improved results over AAV-transduced cytosolic data, while soma-targeted jGCaMP8s data yielded the best overall performance, indicating that neuropil contamination is the primary driver of artifactual correlations in cytosolic imaging data.

### Pairwise activity correlations are sensitive to imaging resolution and analysis pipeline

Having examined orientation tuning curve correlations in OS neurons, we next assessed pairwise activity correlations of all ROIs as a function of distance between their somata to determine whether artifacts caused by neuropil contamination and low axial resolution extended to other network-level analyses. We calculated the ground-truth pairwise activity correlation coefficients using ΔF/F_0_ traces of L2/3 neurons expressing nuclear-GCaMP6s^102^, and, for the population, averaged the coefficients across all pairs of neurons within similar pairwise distances (with a bin width of 30 μm, gray dashed lines, **Figure 7**; **Methods**). We observed a weak correlation averaging around 0.1 over 200 μm pairwise distance, which slightly increased to 0.12 for neurons within 30 μm of each other.

**Figure 7.**
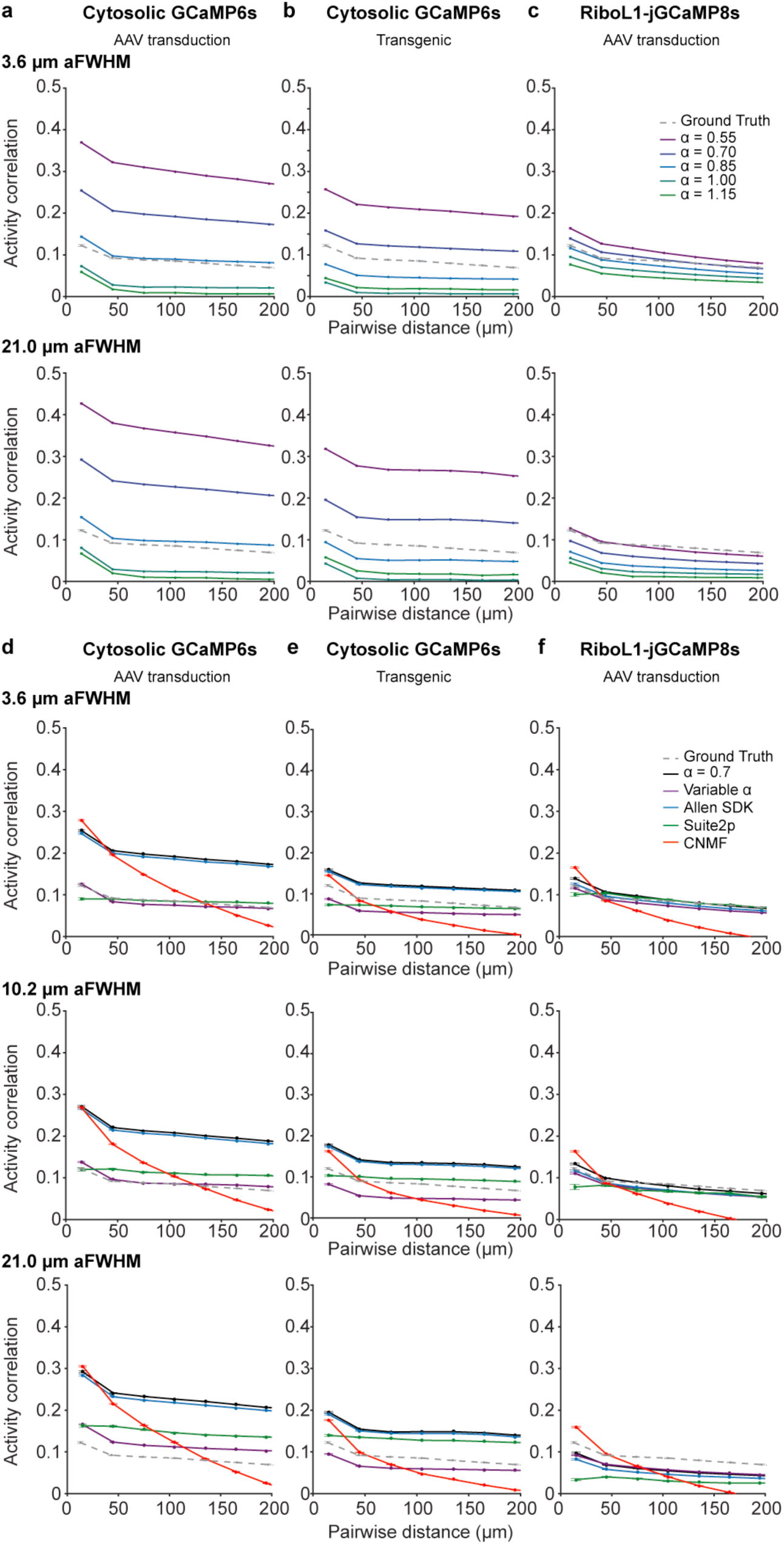
Pairwise activity correlations are sensitive to neuropil subtraction method and imaging resolution. (a,b,c) Pairwise calcium activity correlation for manually segmented ROIs processed with neuropil subtraction at a constant α coefficients of 0.55, 0.7, 0.85, 1.0, or 1.15, for datasets acquired at 3.6-μm and 21.0-μm aFWHM for V1 L2/3 neurons with virally transduced GCaMP6s, transgenically expressed GCaMP6s, and virally transduced RiboL1-jGCaMP8s expression, respectively. Gray dashed line: pairwise activity correlation vs. distance for 1,489 neurons labeled with nuclear-targeted GCaMP6s from 2 mice, 7 FOVs; data from Ref. (104). **(d,e,f)** Pairwise activity correlation for ROIs from “α = 0.7” (black), “Variable α” (purple), “Allen SDK” (blue), “Suite2p” (green), “CNMF” (red) analysis pipelines for datasets acquired at 3.6-μm, 10.2-μm, and 21.0-μm aFWHM for V1 L2/3 neurons with virally transduced GCaMP6s, transgenically expressed GCaMP6s, and virally transduced RiboL1-jGCaMP8s expression, respectively. Error bars: s.e.m.

Similar to orientation tuning correlation, for cytosolic sensors, we observed wildly different correlation behaviors of hand-segmented ROIs depending on the value used for fixed-⍺ neuropil subtraction (⍺ values of 0.55, 0.7, 0.85, 1.0, and 1.15; **Figure 7a,b**). Compared to the ground-truth correlation, α ≤ 0.7 and α ≥ 1.0 consistently produced correlations that were too high and too low, respectively. While α = 1.0 generated correlation curves most like the ground truth for neurons expressing virally transduced GCaMP6s (**Figure 7a**), it led to correlations for neurons expressing GCaMP6s transgenically that were too low (**Figure 7b**). Activity correlations of neurons expressing soma-targeted jGCaMP8s exhibited a more moderate dependence on the value of α, but higher α values still yielded smaller correlation coefficients (**Figure 7c**). Across the animal preparations and aFWHMs, no single α value reproduced the ground-truth correlation.

For cytosolic sensors (**Figure 7d,e**), “⍺ = 0.7” and “Allen SDK” produced highly similar traces with the strongest activity correlations across all aFWHM conditions. “Variable ⍺” and Suite2p yielded closely matched long-range correlation coefficients at approximately half the values of those from “⍺ = 0.7” and “Allen SDK”; they also yielded correlation traces that most closely resemble the ground truth. However, Suite2p again failed to capture the elevated correlations at distances < 50 µm. CNMF diverged most substantially from the other methods, generating large short-range correlations that decayed to zero at ∼180 µm.

The most consistent and accurate results across analysis pipelines were obtained from the soma-targeted jGCaMP8s dataset acquired at 3.6-µm aFWHM: all methods except CNMF yielded correlation coefficients peaking at ∼0.10 – 0.14 at short distances and decreasing gradually to ∼0.07 at 200 µm (top panel, **Figure 7f**). At larger aFWHMs, activity correlation traces showed increasing deviations from the ground truth (bottom panels, **Figure 7f**).

Across both cytosolic and soma-targeted data, pairwise activity correlations depended strongly on aFWHM and analysis pipeline (**Figure 7d-f**, **Figure S9**), confirming that sensor-expression-, resolution-, and pipeline-dependent artifacts extend beyond tuning correlation to network-level activity measurements more broadly.

## Discussion

In this study, we systematically characterized how axial resolution, sensor expression strategy, and analysis pipeline jointly determine the accuracy of in vivo 2P calcium imaging. By acquiring calcium imaging datasets from the same L2/3 neurons in V1 of awake mice at five aFWHM values (3.6 to 21.0 µm) representative of 2PFM systems used in neuroscience, including recently developed lower-resolution systems, and by analyzing the same datasets through five pipelines, we found that reducing axial resolution introduces errors that no analysis pipeline could fully correct.

### Resolution-dependent artifacts in characterizing the functional and network properties of V1 neurons

As aFWHM increased, fluorescence from individual ROIs became increasingly mixed with fluorescence from surrounding neuropil and neighboring cell bodies. This leads to two sources of errors. The contaminating fluorescence reduces the amplitude of the ROI’s ΔF/F_0_; for neurons with low firing rates and thus small calcium transients, this reduction can cause genuinely visually responsive cells to be misclassified as non-responsive. The calcium activity from neighboring cell bodies or surrounding neuropils also introduces false transients, converting non-responsive neurons into apparently responsive ones and distorting the orientation-selectivity characterization of visually responsive cells. Across all sensor expression strategies with cytosolic GCaMP, this contamination drove a stereotyped pattern of artifacts that were increasingly severe at large aFWHMs: somatic ΔF/F₀ magnitudes were attenuated, neurons were misclassified in terms of visual responsiveness and orientation selectivity, and network-level measurements such as pairwise tuning correlations and activity correlations deviated from the ground truths. These artifacts persisted across hand-segmented and automatically detected ROIs, fixed and cell-specific neuropil subtraction coefficients, and subtraction-based and demixing-based pipelines.

In V1, neuropil is highly visually responsive. Because the neuropil within the excitation focus often consists of axons and dendrites of many neurons, its visual response is less orientation-selective than that of individual L2/3 somata. The progressive failure to remove neuropil contamination at larger aFWHMs was therefore reflected in the systematic decrease of gOSI values across the V1 population. Failure to distinguish signals of ROIs of interest from surrounding neuropil or nearby cell bodies led to classification errors in terms of visual responsiveness and orientation selectivity.

These single-cell artifacts propagated to network-level analyses. Neuropil activity, often modulated by behavioral or brain state, is highly correlated in space^72,103–105^. Failure to fully remove neuropil contributions introduces a common signal across neurons within the same FOV. This shared neuropil contamination inflated correlations between neurons artifactually, as we observed for most analysis pipelines on cytosolic GCaMP6s data; the effect grew more severe with increasing aFWHM. Because correlated neuropil activity is widespread in the brain, failure of the analysis pipelines tested here to remove neuropil contamination would skew interpretations of population dynamics in brain regions beyond V1.

### Limitations of current analysis pipelines in addressing neuropil contamination

We found that the choice of analysis pipeline can dominate the apparent properties of calcium imaging datasets. The five analysis pipelines tested here, from neuropil subtraction with a fixed (⍺ = 0.7) or cell-specific ⍺ (Variable ⍺, Allen SDK) on hand-segmented ROIs, to automated ROI detection with neuropil subtraction (Suite2p) or demixing with constrained non-negative matrix factorization (CNMF), generated varied and often artifactual classification and correlation analysis results from the same calcium imaging datasets.

For cytosolic GCaMP6s expression, this occurred even for datasets acquired at the highest axial resolution: varying the neuropil subtraction coefficient ⍺ on hand-segmented ROIs reproduced the range of quantitatively divergent results on the spatial scale of orientation tuning organization in mouse V1 reported in the literature. “Variable ⍺”and Suite2p generated correlations that were most similar to the ground truths, but neither matched the ground truths across all preparations. Suite2p captured the lower correlations at larger distances but missed the heightened correlation for nearby neurons, due to its smaller number of detected ROIs. With its more complex processing steps – including spatial filtering, temporal denoising, baseline estimation, and signal demixing – CNMF consistently generated results most distinct from the methods based on neuropil subtraction, reporting much slower decay of correlation over pairwise distance. Given that ROIs detected by CNMF have areas on average 1.40-fold larger than hand-segmentation ROIs and 1.94-fold larger than those detected by Suite2p (example FOVs, **Figure S10**; CNMF detecting bigger ROIs were also reported previously^106^), we speculated that CNMF failed to spatially separate local neuropil from somata in densely labeled cytosolic data, and, by expanding the somatic spatial components into surrounding neuropil pixels, artifactually reduced the gOSI values and inflated activity correlations.

Our finding that pipeline choice can affect quantitative outcomes even at high resolution is consistent with prior work of calcium imaging in the mouse brain, where the same underlying limitation of commonly used pipelines – their difficulty in demixing and removing contaminating activity – produces a related failure mode. For calcium imaging of CA1 neurons acquired at high resolution (i.e., with a 0.8 NA objective^107^), CNMF and Suite2p failed to reject false transients arising from overlapping neurons or dendrites, which accounted for 15-20% of all transients^108^. For one dataset, the contamination led to an 18% overestimation of place cells and shifted 19% of place fields by more than 10 cm.

Together, our studies indicate that uncorrected contamination is a problem even for data acquired at high resolution and can influence quantitative outcomes. As aFWHM increased, the contamination became even more severe and all pipelines exhibited increasing deviation from the ground truths for both cytosolic and soma-targeted sensors. The failure of these analysis pipelines to correct resolution-dependent artifacts is, in retrospect, not surprising. Subtraction-based methods rely on a scaled neuropil estimate, but the appropriate value of the scaling coefficient ⍺ depends on the volumes and positions of the soma and surrounding neuropil within the excitation focus, as well as on the local sensor concentrations – parameters that existing methods cannot capture accurately. Moreover, these methods assume that the contaminating signal from neuropil structures above and below the ROI can be represented by the signal in the neuropil mask, which originates from structures lateral to the ROI. With the excitation focus encompassing larger volumes at larger aFWHMs – including other cell bodies – signal in the neuropil mask may become increasingly distinct from the actual contamination. For these reasons, cell-specific ⍺ estimation methods (Variable ⍺, Allen SDK) improved performance only marginally. Demixing-based methods such as CNMF can in principle separate overlapping sources, but the core assumptions in the matrix factorization (sparse spatial footprints, low-rank background) become increasingly violated as the focal volume expands and more neurons and neuropil contribute to each pixel. As a result, none of these methods can recover information that has been irretrievably mixed during image acquisition.

### Soma-targeted sensors provide partial protection against resolution-dependent artifacts

Soma-targeted sensors provided partial, but incomplete, protection against these artifacts. Targeting jGCaMP8s expression to the somatic compartment reduced neuropil fluorescence, enabling accurate correlation measurements for the highest-resolution datasets across all pipelines except CNMF. Soma-targeted data also fared best at low resolution. However, the residual neuropil expression and false transients from overlapping cell bodies were sufficient to generate classification errors at large aFWHMs. Soma-targeted expression also yielded substantially dimmer cell bodies than cytosolic sensors, consistent with reports on soma-targeted sensors in which much higher laser power was used for 2P excitation^80,92^. A newer construct for soma-targeted ribo-jGCaMP8s appears to require less excitation power^109^, suggesting that continued engineering efforts may offer an optimal solution combining brightness with high somatic restriction. Similarly, nuclear-targeted sensors offer a potential alternative, provided their slower kinetics and lower sensitivity suit the biological question^91,102,110–112^.

### Implications for microscope design

Our findings have direct implications for the growing class of 2PFM systems that trade axial resolution for capabilities such as larger fields of view, faster volumetric coverage, or miniaturized geometries for freely moving recordings. These systems enable experiments that higher-resolution microscopes cannot perform, and our results should not be read as an argument against their use. Rather, our results indicate that when the analysis methods tested above are applied to datasets acquired with such systems, quantitative claims about both the properties of individual neurons and network-level statistics such as spatial correlation need to be treated with caution. For studies whose biological conclusions depend on such measurements, restricting sensor expression to neuronal somata or nuclei, combined with careful choice of analysis pipeline and tests of the robustness of those conclusions against pipeline and parameter choices, provides some mitigation. Whenever possible, however, new imaging systems should treat high axial resolution as a design priority.

### Limitations of this study

Several limitations of this study deserve mention.

First, our analyses were restricted to V1 L2/3 neurons imaged through a 170-µm-thick cranial window in quietly awake mice; the magnitude of the artifacts we describe may differ in deeper cortical layers, through thicker windows or prism implants, in other brain regions, in actively behaving mice, or in different species. Sample-induced aberrations and scattering, which worsen with thicker windows and imaging depth, would further degrade resolution and likely amplify the trends we report. At larger depths, out-of-focus excitation of more superficial sensor-expressing structures would become an additional source of contamination^113^.

Second, we focused our evaluation on pipelines in widespread use, so that our results are representative of what most neurobiologists obtain in practice. Although we varied selected parameters for Suite2p and CNMF, we did not exhaustively explore the full parameter space, and we cannot exclude the possibility that alternative parameter combinations may mitigate some of the artifacts described here. Nevertheless, because our analyses largely employed the default parameters of each pipeline, the results we report are likely representative of the outcomes obtained by typical users. Other advanced approaches may perform better than the five pipelines tested here: EXTRACT^67,68^ uses the theory of robust statistics to generate a loss function that minimizes the influence of contaminating sources such as neuropil or overlapping neurons; SEUDO^108^ models contaminating cell bodies and neurites as a superposition of independent Gaussian kernels with exponentially distributed amplitudes and can remove false transients from individual frames. We therefore provide our raw datasets to the community for further evaluation (**Supplementary Data**).

Third, our characterization focused on visual responsiveness, orientation selectivity, and pairwise correlations. Whether and how neuropil contamination and reduced axial resolution affect other measurements of neuronal and population properties, such as dimensionality, decoding accuracy, or noise correlation, remains to be determined.

### Best practices and recommendations

Based on our findings, we make the following recommendations for maintaining the accuracy of calcium imaging experiments:

#### Imaging

Whenever possible, acquire calcium imaging data at high resolution with an excitation NA ≥ 0.8, a condition that is satisfied by most commercial 2PFM systems.

#### Sensor expression

When high axial resolution cannot be achieved, as is often the case with large-FOV systems or miniaturized microscopes, soma-targeted sensors should be used to reduce, though not eliminate, neuropil artifacts. Nuclear-targeted sensors can fully remove neuropil contamination and are appropriate when high temporal resolution is not required. When cytosolic sensors must be used, sparse labeling of the population of interest may reduce neuropil contamination and limit artifacts.

#### Analysis

Different analysis pipelines yield different quantitative results even at optimal resolution. For hand-segmented ROI methods, cells with high structural contrast should be selected, and SNR thresholding together with a cell-specific α should be incorporated to improve performance. Pipelines that are more conservative in ROI detection, and consequently yield ROIs of lower density (e.g., Suite2p), may miss short-range activity correlations. When applying CNMF to cytosolic data, one may need to use background models larger than the default rank-1 per FOV to more accurately capture neuropil contamination.

#### Validation

Biological conclusions should be verified for robustness against the choice of analysis pipeline and parameters (for example, Refs. 102,114). In particular, conclusions should not depend on the value of the neuropil subtraction coefficient *α*, nor on the method used to determine it. When such dependence is observed, additional scrutiny is required before biological conclusions are drawn: soma- or nuclear-targeted sensors should be used if possible and, whenever feasible, results should be validated against electrophysiological ground truths.

Beyond cross-pipeline comparison, post-processing validation provides a complementary and practically accessible safeguard^115^. Because no single method fully certifies the extracted signals, several should be used in combination. First, frame-by-frame visual inspection of ROI spatial profiles in the raw calcium movie during purported transients can reveal false transients arising from overlapping cell bodies^108^ or neuropil bleed-through. Second, self-consistency metrics test whether the extracted spatial and temporal components jointly account for local pixel variance without residual structured activity. Third, cross-validation of population responses against simultaneously recorded behavioral or stimulus events can offer further diagnostic power^115^.

#### Cross-study comparison

These considerations also bear on the comparison of results across studies and laboratories. Microscopes differ between labs. Even for the same model of microscope, underfilling the microscope objective leads to different excitation NA and thus axial resolution. A thicker cranial window, a tilted brain, or a misuse of objective correction collar further increases aFWHM. Furthermore, as we demonstrated here, because pipelines differ in how they detect ROIs, weight pixel contributions, and remove neuropil signal, different pipelines can yield substantively different single-neuron properties and network-level statistics even when the underlying imaging data are identical. Therefore, to increase reproducibility and help resolve contradictory results, it is important to report microscope resolution (measurable using fluorescent beads) as well as details on sample preparation, pipeline choices, and parameter settings.

### Conclusion

In summary, for accurate quantitative population calcium imaging, high axial resolution during image acquisition is the most reliable safeguard against the artifacts we describe; for systems in which resolution is necessarily compromised, soma-targeted sensor expression, combined with careful analysis choices, can mitigate—but does not eliminate—these artifacts. Our findings underscore that the increasing popularity of 2PFM systems of lower axial resolution brings with it a necessity to characterize the failure modes of the resulting data, and to interpret quantitative claims about neuronal and network activity from these imaging systems with caution. Given that different pipelines produce different quantitative results even for cytosolic data acquired at the highest resolution, new analysis pipelines that can accurately separate the contributions of somata from those of their neuropil remain urgently needed to make calcium imaging a quantitatively accurate method for neuronal activity measurement.

## Author Contribution

N.J. conceived the project; H.Y.A.Y. performed surgery, acquired all *in vivo* imaging data, and analyzed all data; U.A. acquired and analyzed bead data; G.F. contributed to figure preparation; A.S.C. provided computational simulation tools; H.Y.A.Y. and N.J. wrote the manuscript with input from all authors.

## Acknowledgement

This work was supported by the National Institutes of Health BRAIN Initiative UF1NS107696 (N.J.), 5U19NS107613 (N.J.), 1R01NS109553 (N.J.), U01NS118300 (N.J.), and NIBIB R01EB037653 (A.S.C.). Supplementary Data were deposited to the BioImage Archive^116^ and are publicly accessible under accession number S-BIAD3802.

## Supplementary Codes

Analysis codes in MATLAB® and Python are available at: https://github.com/JiLabUCBerkeley/Neuropil. All *in vivo* calcium imaging data are available at BioImage Archive^116^: https://www.ebi.ac.uk/biostudies/bioimages/studies/S-BIAD3802.

## Methods

### Animal use

All animal experiments followed National Institutes of Health guidelines for animal research. Procedures and protocols on mice were approved by the Animal Care and Use Committee at the University of California, Berkeley (AUP-2020-06-13343).Wild-type (C57BL/6J, JAX 000664) and transgenic (Slc17a7-IRES2-Cre×Ai162D (+/+)^89^, JAX 023527×JAX 031562) mice of both sexes, older than 2 months, were used for *in vivo* imaging. Mice were housed under 12-hour light/dark cycle, with ambient temperature between 20 and 26 °C and humidity between 40 and 60%.

### Craniotomy and virus injection in the visual cortex

Cranial window implantation and virus injection procedures were performed as previously described^85^. Mice were anesthetized with 1-2% isoflurane (by volume) in O_2_ and given the analgesic buprenorphine subcutaneously (0.3 mg per kg of body weight). Mice were head-fixed in a stereotaxic apparatus (Kopf instruments). A 3-mm craniotomy was created over the left V1, centered 2.5 mm lateral and 1 mm anterior to lambda. For wild-type mice, virus injection was performed using a glass pipette beveled at 45° with a 15-20-µm opening, back-filled with mineral oil. A fitted plunger controlled by a hydraulic manipulator (Narashige, MO-10) was inserted into the pipette to inject viral solution into the cortex at 250 µm below the pia. At each injection site, 30 nL viral vectors encoding GCaMP6s^74^ or RiboL1-jGCaMP8s^80^ (AAV2/1-hSyn-GCaMP6s with 1.76×10^13^ titer, AAV2/PHP.eB-hSyn-RiboL1-jGCaMP8s with 3.40×10^13^ titer; both diluted to 1.3×10^13^ titer) were injected over 1 min. The pipette was kept in the brain for 3 min after injection before being withdrawn. 12-18 injections were made across the exposed cortex with an average spacing of 500 µm. The cranial window consisted of a 3.5-mm-diameter 170-µm-thick glass disk and a glass ring with 3 mm inner diameter, concentrically attached with a UV-cured optical adhesive (Norland NOA 61). The cranial window was embedded in the craniotomy and sealed with 3M Vetbond tissue adhesive. An aluminum head-bar was then attached to the skull with cyanoacrylate glue and dental acrylic. Mice were allowed to recover for at least 3 weeks before experiments.

### Visual stimulation

Visual stimuli were generated using the Psychophysics Toolbox^117^ and delivered via a blue LED light source (450–495 nm, SugarCUBE) and back-projected onto a Teflon® screen (McMaster-Carr) using a custom-modified projector^118^. The screen was positioned 15 cm from the right eye and oriented at 55° to the long axis of the animal, covering 80° × 80° of visual space. Drifting grating stimuli with 100% contrast, a spatial frequency of 0.041 cycles per degree, and a temporal frequency of 2 Hz were presented to the animal. Each trial consisted of 12 gratings drifting in one of 12 directions (0°–330° in 30° increments) for 3 s, and presented in pseudorandom sequences with an interleaved black screen lasting 3 s. With 10 trials, each data acquisition session lasted 12 min.

### Fluorescent bead sample preparation, imaging, and analysis

Bead samples were prepared using two methods depending on the bead size. 0.5- and 2-μm-diameter carboxylate-modified FluoSpheres® (yellow-green fluorescent 505/515 nm, Thermo Fisher Scientific F8813 and F8827) were immobilized onto poly-L-lysine coated coverslips via electrostatic attraction as follows. A circle was drawn on a coverslip using a Sharpie, and its hydrophobic ink then confined poly-L-lysine solution (20 μL, 10 mg/mL) within the marked area. After the poly-L-lysine solution had dried completely, a diluted fluorescent beads solution (40 μL, 1:1000 in deionized water) was pipetted to the poly-lysine coated region and allowed to dry completely.

The 10- and 15-μm-diameter polystyrene FluoSpheres® (yellow-green fluorescent 505/515 nm, Thermo Fisher Scientific F8836 and F8844, respectively) were embedded in agarose as follows: A 4% agar solution was microwaved for 15 seconds and pipetted into a small petri dish. As the agar began to solidify, the fluorescent beads were evenly spread across the surface to prevent clumping.

Bead samples were imaged with a homebuilt 2P fluorescence microscope^70^ with a 16×0.8NA objective (Nikon, CFI75 LWD 16X W) under different axial resolution by underfilling using a motorized beam reducer (Special Optics, 1-4×motorized beam expander) set at 0.25×, 0.33×, 0.5×, or 1×, respectively. 3D stacks were acquired at pixel sizes ranging from 0.15 µm to 2 µm at a frame rate of 1 or 2.3 Hz and a post-objective power of 1.8 – 15.7 mW. At each z position, 10 frames were acquired and their average was used for further analysis.

Beads were first detected on the maximum-intensity projection (MIP) of the z stack acquired at the 1× beam reducer setting using Laplacian-of-Gaussian blob detection (σ range: 0.8*σ*_exp_ to 2*σ*_exp_, 14 levels, 5% threshold; 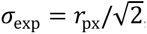, where *r*_px_ is the nominal bead radius in pixels) implemented in scikit-image (Python package, v0.21.0). Detection was then run independently in Z stacks acquired for the same beads at more underfilled conditions. The detected blobs were then matched to the centroids from the 1× dataset by nearest-neighbor search using pairwise Euclidean distances (tolerance 20 pixels; scipy.spatial.distance.cdist, SciPy v1.11.1). Only beads successfully detected in all five beam sizes were retained.

From the 3D stack of detected bead, a one-dimensional axial intensity profile was extracted by averaging pixel values over a 3 × 3 pixel region centered on the bead at each z slice. A one-dimensional Gaussian function

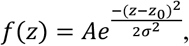

where *A* is the amplitude above baseline, *z₀* is the axial center, and *σ* is the width parameter, was fit to the baseline-subtracted profile by minimizing the weighted sum of squared residuals (scipy.optimize.curve_fit, SciPy, Python package, v1.11.1), 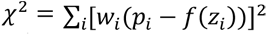, where *i* indexes each z slice in the profile, *p_i_* is the baseline-subtracted intensity, *f*(*z_i_*) is the Gaussian model evaluated at depth *z_i_*. Here *w_i_*, the weight assigned to each slice *i*, is proportional to its normalized intensity:

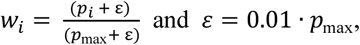

where *p*_max_ is the maximal intensity of the extracted profile after baseline subtraction. The axial FWHM was calculated from the fitted *σ* as:

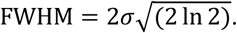

The fitted Gaussian amplitude *A* was divided by the measured post-objective laser power squared to account for the quadratic power dependence of two-photon fluorescence signal:

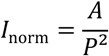

where *P* is the laser power measured at the sample (mW). Values were then normalized for each bead to its value under the 1× beam expansion condition:

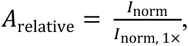

with their average and standard-error-of-the-mean values for all beads plotted in **Fig. 1b**.

### In vivo 2P imaging at variable axial resolution

Awake mice were habituated to head fixation for 15 min per day over three days before imaging. A 3D resin-printed head cone was attached to the headbar with Vetbond and dental cement to block stray light during imaging. The headbar was rigidly secured to a custom stage, and the animal body was gently restrained within a cylindrical tube to prevent running and minimize motion artifacts during imaging. A homebuilt 2P fluorescence microscope^70^ was used for *in vivo* imaging with either a 16× 0.8NA (Nikon, CFI75 LWD 16X W) or a 10× 0.6NA (Olympus, XLPLN10XSVMP) objective lens and the 920 nm output of a femtosecond laser system (Spectral Physics, InSight DeepSee). Images over a 400 × 400 µm FOV were acquired at 2 Hz with 1 µm pixel size. Axial resolution was adjusted by underfilling either objective using a motorized beam reducer (Special Optics, 1-4×motorized beam expander). For *in vivo* imaging, the 16× 0.8NA objective was used for axial FWHMs of 3.6 µm, 5.7 µm, 10.2 µm and the 10× 0.6NA objective was used for axial FWHMs of 13.4 µm and 21.0 µm. Bead images were acquired with the 16×0.8NA objective.

19 FOVs were imaged across 10 animals (wild-type mice expressing GCaMP6s: 4 animals, 6 FOVs; Slc17a7-IRES2-Cre×Ai162D: 3 animals, 6 FOVs; wildtype mice expressing RiboL1-jGCaMP8s: 3 animals, 7 FOVs). Each FOV was imaged five times at distinct axial FWHMs. Post-objective power ranged from 78 to 160 mW for virally transduced GCaMP6s expression, 60 to 137 mW for transgenically expressed GCaMP6s, and 96 to 201 mW for virally transduced RiboL1-jGCaMP8s. Exact post-objective powers used for each *in vivo* imaging dataset are listed in **Table S9**.

### Image registration

Imaging data were processed using Fiji^119^ and custom MATLAB^®^ and Python scripts (**Supplementary Code**). Frames acquired from the same FOV were registered using the non-rigid motion correction algorithm NoRMCorre^81^, with datasets acquired at 3.6-µm aFWHM as the reference. The motion-registered frames were used for all subsequent analyses.

### Analysis for functional classification of manually segmented ROIs

ROIs were manually segmented using time-averaged images acquired at 3.6 µm aFWHMs. For each ROI, F_raw_(t) was extracted by averaging the pixel values within the ROI from each frame, and F_neuropil_(t) was extracted by averaging the pixels in the surrounding neuropil mask (within 30 µm from the centroid of each ROI, excluding pixels within ROI masks) (e.g., “F_raw_” and “F_neuropil_” traces in **Figure 2b**). For each 6-s-long trace acquired during a 3-s-long dark screen and then a 3-s-long drifting grating stimulus, we calculated the neuropil baseline fluorescence value F_0, neuropil_ as the mean of F_neuropil_(t) during the 3 s of dark screen preceding the grating. We then calculated ΔF_neuropil_(t) = F_neuropil_(t) − F_0, neuropil_ for this specific 6-s-long grating period, scaled it by ⍺, and obtained F_ROI_(t)=F_raw_(t) − ⍺⋅ΔF_neuropil_(t) (e.g., “⍺⋅ΔF_neuropil_” and “F_ROI_” traces in **Figure 2b**, for all 120 6-s-long grating periods of each data acquisition session).

The neuropil subtraction coefficient (⍺) was either fixed for all ROIs (“⍺ = 0.7”, or another specified value) or variable for each ROI (“Variable ⍺” and “Allen SDK”). For “Variable ⍺”, “estimateNeuropil” function (provided by Dr. Dipoppa^73^) was used to calculate ⍺ for each ROI from its F_raw_(t) and F_neuropil_(t). For “Allen SDK”^75,79^, “allensdk.brain_observatory.r_neuropil.estimate_contamination_ratios” function from AllenSDK (version 2.16.2) was used to calculate ⍺ for each ROI from F_raw_(t) and F_neuropil_(t).

For each ROI, we then calculated F_0,ROI_ as the mean F_ROI_(t) across 3 s of dark screen, and then ΔF_ROI_(t)/F_0,ROI_ for the 6-s-long trace with ΔF_ROI_(t)=F_ROI_(t) − F_0,ROI_. Concatenating 120 such 6-s-long ΔF/F_0_ traces in the order of their acquisition gave us the full 720-s-long ΔF/F trace (not shown). Averaging 10 trials for each grating stimulus, we obtained the trial-averaged ΔF/F_0_ traces for all 12 grating stimuli types (e.g. traces during the 1-s dark screen before grating onset and the 3-s-long grating presentation, **Figure 2c**).

Averaging the data points of the trial-averaged ΔF/F_0_ within the 3-s-long grating presentation, we obtained a ΔF/F_0_ value for each of the 12 grating types, defined as R(*θ*), with *θ* being the orientation of the grating from 0 to 330° at 30° interval. Normalizing to the maximum ΔF/F_0_ value among the 12 values, we obtained a tuning curve for this ROI (e.g., **Figure 2d**).

To determine whether an ROI responded to visual stimulation, we calculated the time averaged ΔF/F_0_ value within each of the ten 3-s-long presentations of the same grating stimulus and tested this distribution with a two-sided Student’s t-test to determine whether it was significantly different from zero. An ROI was defined as visually responsive (VR) if its response to at least one of the 12 grating stimuli was significantly different from zero (p < 0.05).

For VR neurons, we then compared the 12 sets of ΔF/F_0_ values for each grating stimulus (10 values per set, as each grating stimulus was repeated 10 times). Neurons were defined as orientation-selective (OS) if the sets were significantly different (one-way ANOVA, p < 0.05).

On the population level, for neurons within the same FOV imaged at different aFWHMs, we compared the distributions of the baseline fluorescence F_0,ROI_ and transient magnitude ΔF_ROI_. For each ROI, we averaged the 120 F_0, ROI_ values (from 120 3-s-long dark screen periods) to obtain the F_0, ROI_ value for each aFWHM condition and normalized to the value measured at 3.6-µm aFWHM (e.g., **Figure 2g**). The same averaging and normalization were performed for transient magnitude ΔF_ROI_ (e.g., **Figure 2h**). Two-sample Kolmogorov-Smirnov tests were performed against the distributions of datasets acquired at 3.6-µm aFWHM.

For each VR neuron, we identified the grating stimulus that gave rise to the largest *R*(*θ*) and tested whether the distributions of these maximal *R*(*θ*) values (e.g., “Max ΔF/F_0_”, **Figure 2i**) decreased relative to those at the previous, smaller aFWHM condition using one-sided Wilcoxon rank sum tests.

For each OS neuron, we computed its global orientation selectivity index (gOSI^120^; e.g. **Figure 2n**) as 1 – circular variance and the preferred orientation (PO; e.g. **Figure 2q,r**) using the trial- and time-averaged responses (defined above as *R*(*θ*)), with

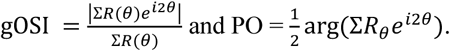

Perfect orientation selectivity is indicated with gOSI of 1, whereas a gOSI of 0 indicates no orientation selectivity.

The same approaches were applied to F_neuropil_ traces to determine the baseline brightness of the neuropil F_0,neuropil_ (e.g., **Figure 2j**), the neuropil transient magnitude ΔF_neuropil_ (e.g., **Figure 2k**), the maximal neuropil response (e.g., “Max ΔF/F_0_”, **Figure 2l**), and the gOSI (e.g., **Figure 2o,p**) and PO of the neuropil (e.g., **Figure 2q**).

### SNR thresholding for manually segmented ROIs

For comparison across analysis pipelines, we excluded manually segmented ROIs with poor SNR by applying an SNR filter adapted from EXTRACT^67,68^. For each ROI, the signal was defined as the maximal ΔF/F_0_ value of its full 720-s-long ΔF/F_0_ trace (i.e., max(ΔF/F_0_)). The noise *σ*_noise_ was calculated in the frequency domain using a power spectral density (PSD) approach: the ΔF/F_0_ trace (1,440 time points from 2 Hz × 6 s × 120 grating stimulation) was Fourier transformed, and the PSD was computed for first half of the output (the first 720 values of the FFT output, since the FFT produces a symmetrical spectrum where the second half is a redundant complex conjugate mirror of the first) and normalized to reflect single-sided power spectrum. The mean of the PSD values at high-frequency components P_high_ (> 0.5ƒ_Nyquist_, corresponding to the last 360 points of the single-sided PSD) was used to calculate the noise variance *σ*_noise_ as:

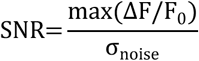

For each ROI, SNR was then defined as the ratio of max(ΔF/F_0_) and *σ*_noise_:

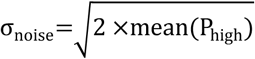

Only ROIs with an SNR > 3.5 were retained for analysis.

### Automatic ROI detection and neuropil subtraction with Suite2p

Motion-registered images were used as input to Suite2p (Python package, version 0.13.2.dev19). The Suite2p GUI was used to inspect ROI detection, and we output the ROI masks, neuropil masks, F_raw_(t), F_neuropil_(t), and F_ROI_(t) (with neuropil subtraction using F_neuropil_(t) and ⍺ = 0.7) from Suite2p for further analyses. From F_ROI_(t) of each automatically detected ROI, we calculated its full ΔF/F_0_ trace, from which we determined the visual responsiveness of the ROI; for VR ROIs, we identified their largest *R*(*θ*) values (“Max ΔF/F_0_”, **Figure S6i**); for OS ROIs, we calculated their gOSI values (**Figure 5j**).

Default parameters^65,93^ were used during processing with the following exceptions. The sampling rate (“fs”) was set to 2 Hz, the frame rate of our data acquisition. For “threshold*_*scaling”, a parameter that controls ROI detection sensitivity, we tested values of 0.5, 1, and 2 on an example FOV with cytosolic GCaMP6 expression (**Figure S5b**) and chose the default value of 1 for all cytosolic GCaMP6s data. For soma-targeted jGCaMP8s, we set “threshold*_*scaling” to 0.5 to detect more ROIs from these dimmer images.

### Analysis with CNMF

We used the CaImAn implementation^66^ (Python package, version 1.9.11) of CNMF and motion-registered images as input. We used CaImAn’s Jupyter notebooks to inspect ROI detection and exported the spatial mask (A), temporal traces (C), and full ΔF/F_0_ trace for each ROI, as well as the temporal trace (f) and spatial mask (b) of the low-rank background (B). From the full ΔF/F_0_ trace, we determined the visual responsiveness of the ROI; for VR ROIs, we identified their largest R(*θ*) values (“Max ΔF/F_0_”, **Figure S6j**); for OS ROIs, we calculated their gOSI values (**Figure 5l**).

Default parameter values were used with the following exceptions. The frame rate (“fr”) was set to 2 (default = 30; in frames per second), and “K”, the number of components/ROIs to be found within the FOV, was set to 600 (default = 30). For each detected ROI, CNMF computes the peak SNR of its temporal trace averaged over the duration of a typical transient, and an ROI is rejected if the computed SNR is below “min_SNR”. We varied “min_SNR” across values of 0.5, 1, 1.5, 2, 2.5, and 3 on an example FOV to test the effect on the number of ROIs (**Figure S5c**) and chose its default value of 2.5. We also varied “nIter” across 1, 5, and 10, which defines the number of rank-1 refinement iterations for ROI initialization (**Figure S5c**) and chose its default value of 5.

### Detection of shared ROIs across all pipelines

To identify ROIs that were detected across all processing pipelines (“common” column, **Figure S7a**), we calculated the centroid x and y coordinates of manually segmented ROIs as well as those of ROIs detected automatically by CNMF and Suite2p. An automatically detected ROI was considered to represent the same cell as a manually segmented ROI if its centroid coordinates fell within ±5 pixels of those of the manually segmented ROIs. These spatial tolerances were chosen to account for slight variations in ROI boundary detection between pipelines.

### Methods for nuclear-GCaMP6s dataset

Details on sample preparation, data acquisition, and image analyses were described previously^102^. Briefly, adult wild-type mice underwent craniotomy and virus injection (AAV2/1-Syn-H2B-GCaMP6s, 5 × 10^12^ GC/mL, 30 nL/injection site) as described above. *In vivo* 2P imaging was performed using a Thorlabs Bergamo II multiphoton microscope with excitation at 920 nm and a Nikon 16×, 0.8NA objective. Visual stimuli consisted of gratings drifting in 12 directions at 30° increments in a pseudorandom sequence, with 10 repetitions per direction. Each stimulus lasted 10 s: a uniform blue screen was shown for 5 s, followed by a drifting grating for another 5 s. Calcium imaging data were motion-registered using NoRMCorre^81^, and ROIs were segmented using Suite2p. Because the dataset used nuclear-localized GCaMP6s, neuropil subtraction was not performed.

### Calculation of orientation tuning correlation over pairwise distance

For each pair of OS neurons within the same FOV, we calculated the Pearson correlation coefficient of their tuning curves (e.g., **Figure 2d**) and their pairwise distance as the physical distance between the neurons’ centroids. Each cell’s mean trial response vectors across stimulus directions were then fitted with a double-Gaussian, as described in the analysis and statistics section. Correlation coefficients from all FOVs with pairwise distances falling within the same 5-µm bins (bins centered from 7.5 to 97.5 µm with increments of 5 µm) were averaged to obtain the orientation tuning correlation versus pairwise distance (**Figure 6, Figure S8**). The ground truth trace was obtained from the nuclear-targeted GCaMP6s dataset acquired previously^102^.

### Calculation of pairwise activity correlation over distance

Pairwise activity correlation analysis was performed on all ROIs. For every pair of ROIs within the same FOV, we calculated the Pearson correlation coefficient of their full ΔF/F_0_ traces and the pairwise distance as the physical distance between the neurons’ centroids. Correlation coefficients from all FOVs with pairwise distances falling within the same 30-µm bins (bins centered from 15 to 195 µm with increments of 30 µm) were averaged to obtain the pairwise activity correlation versus distance (**Figure 7, Figure S9**). The ground truth pairwise activity correlation trace was calculated using ΔF/F_0_ traces extracted from the nuclear-targeted GCaMP6s dataset.

### Calculation of error percentages in VR and OS neurons

For “⍺ = 0.7” analysis (**Fig. 2f**, **3f**, **4f**, **5a**, **5b**), both false negatives (VR or OS neurons from 3.6-µm datasets that became not VR or OS at larger FWHMs) and false positives (neurons that were not VR or OS from 3.6-µm datasets that became VR or OS at larger FWHMs) were included in error percentage calculation.

For all the other analysis methods, only false negatives were included in error percentage calculation as detailed below.

To quantify classification errors in VR and OS neurons for the “Variable ⍺” and “Allen SDK” methods, VR and OS neurons identified from the 3.6-µm aFWHM dataset were used as the reference population. Neurons from this population were considered misclassified under lower resolution conditions if they either failed to pass the SNR threshold, or no longer satisfied the criteria for VR or OS classification.

To quantify classification errors in VR and OS neurons for Suite2p analyses, VR and OS neurons identified under the 3.6-µm aFWHM condition were used as the reference population. Neurons from this population were considered misclassified under lower resolution conditions if they either were no longer detected or no longer satisfied the criteria for VR or OS classification at lower resolutions. Because Suite2p automatically detects ROIs independently for each dataset, we implemented a proximity-based matching method to identify ROIs corresponding to the same cells across different aFWHM conditions. For each ROI detected under the 3.6 µm aFWHM condition, we calculated the centroid coordinates and searched for corresponding ROIs in other aFWHM conditions. ROIs were considered to represent the same cell if their x and y coordinates fell within ±5 pixels of each other in both dimensions.

To quantify classification errors in VR and OS neurons for CNMF analyses, VR and OS neurons identified under the 3.6-µm aFWHM condition were used as the reference population. Neurons from this population were considered misclassified under lower resolution conditions if they either were no longer detected or no longer satisfied the criteria for VR or OS classification at lower resolutions. The same proximity-based method as above was used to identify ROIs representing the same neurons across different aFWHM conditions.

