## Supplemental Materials for "High Axial Resolution Is Necessary for Quantitative Two-Photon Calcium Imaging of Neuronal Populations"

**High Spatial Resolution is Essential for Accurate In Vivo Calcium Imaging of Neuronal Populations**

**Movie S1. Motion-registered virally transduced GCaMP6s time series across varying aFWHM conditions.** This movie shows a 144-s-long motion-registered functional images from AAV-expressed cytosolic GCaMP6s under all aFWHM conditions. Images were acquired at 2 frames/s, and the video is rendered at 20 frames/s.

**Movie S2. Trial-averaged GCaMP6s time series across varying aFWHM conditions.** This movie shows the 72-s-long trial-averaged functional images from AAV-expressed cytosolic GCaMP6s under all aFWHM conditions. Images were acquired at 2 frames/s, and the video is rendered at 10 frames/s.

**Movie S3. Motion-registered transgenic GCaMP6s time series across varying aFWHM conditions.** This movie shows a 144-s-long motion-registered functional images from transgenic GCaMP6s under all aFWHM conditions. Images were acquired at 2 frames/s, and the video is rendered at 20 frames/s.

**Movie S4. Trial-averaged transgenic GCaMP6s time series across varying aFWHM conditions.** This movie shows the 72-s-long trial-averaged functional images from transgenic GCaMP6s under all aFWHM conditions. Images were acquired at 2 frames/s, and the video is rendered at 10 frames/s.

**Movie S5. Motion-registered soma-targeted jGCaMP8s time series across varying aFWHM conditions.** This movie shows a 144-s-long motion-registered functional images from AAV-expressed soma-targeted jGCaMP8s under all aFWHM conditions. Images were acquired at 2 frames/s, and the video is rendered at 20 frames/s.

**Movie S6. Trial-averaged soma-targeted jGCaMP8s time series across varying aFWHM conditions.** This movie shows the 72-s-long trial-averaged functional images from AAV-expressed soma-targeted jGCaMP8s under all aFWHM conditions. Images were acquired at 2 frames/s, and the video is rendered at 10 frames/s.

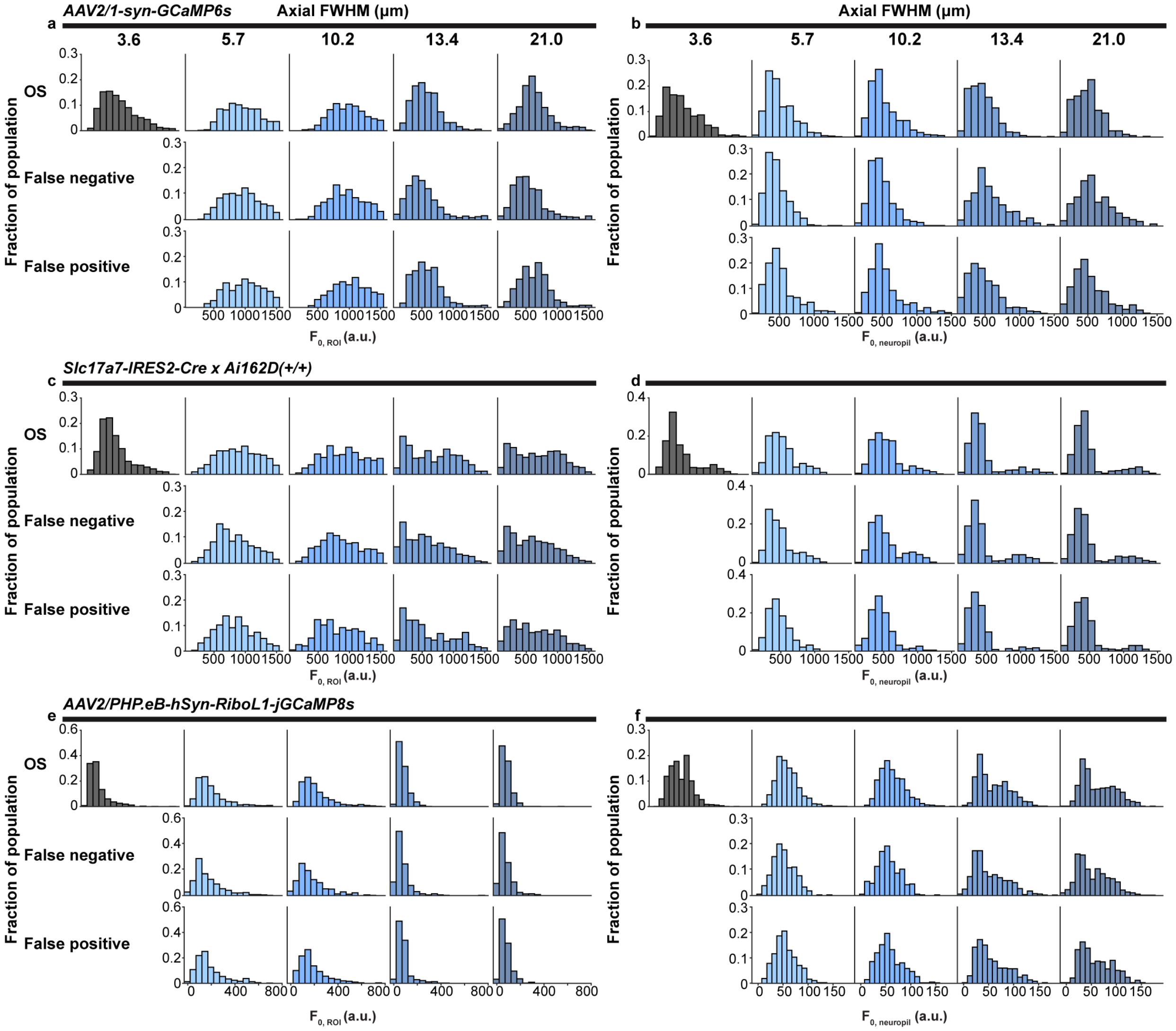


**Figure S1. Histogram distributions of baseline fluorescence ($F_0$) for neurons and their associated neuropils measured with varying axial FWHM. (a,b)** For V1 L2/3 neurons expressing GCaMP6s through viral transduction, distribution of $F_{0,ROI}$ and $F_{0,neuropil}$ for ROIs and their associated neuropil, respectively: (Top row, from left to right) ROIs classified to be OS at 3.6-µm aFWHM, and among these true OS ROIs, the ROIs that are also classified as OS at larger aFWHMs; (Middle row) ROIs that are classified to be VR/OS at 3.6-µm aFWHM but are classified to be not VR (NVR) /not OS (NOS) at larger aFWHMs (i.e., "False negative" neurons); (bottom row) ROIs that are classified to be NVR/NOS at 3.6-µm aFWHM but are determined to be VR/OS at larger aFWHMs (i.e., "False positive" neurons). (**c,d**) Same as (**a,b**) but for V1 L2/3 neurons expressing GCaMP6s transgenically. (**e,f**) Same as (**a,b**) but for V1 L2/3 neurons expressing RiboL1-jGCaMP6s via viral transduction.

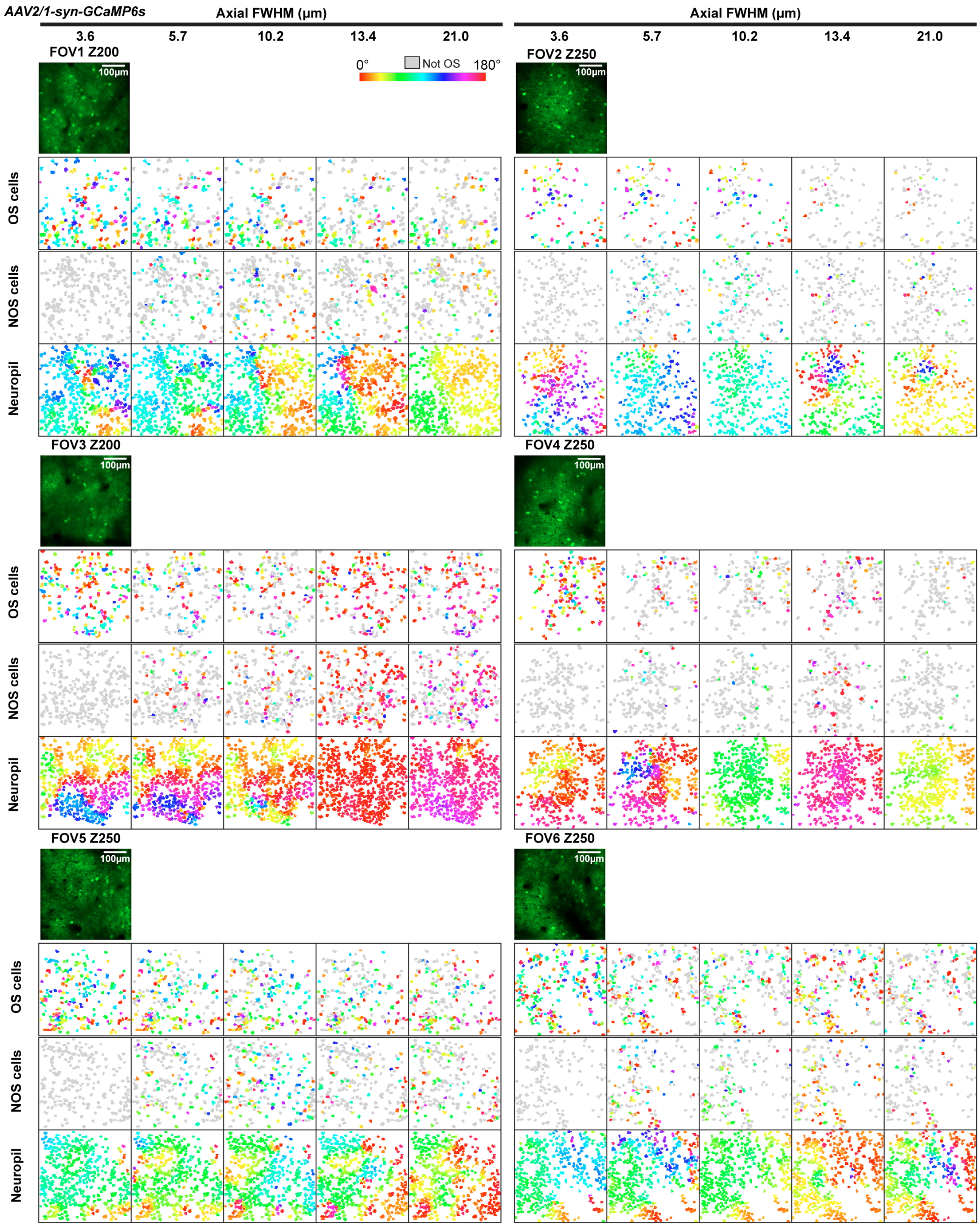


**Figure S2. Preferred orientation map of cells expressing GCaMP6s via viral transduction.** 2P fluorescence images of six FOVs and their color-coded preferred orientation (PO) maps for VR neurons in each FOV. Top and middle rows: maps for cells classified to be OS and not OS at 3.6-µm aFWHM, respectively, color-coded by preferred orientation (gray for not OS). Bottom row: map for corresponding neuropil. FOV 1 is the same FOV as Fig. 2a.

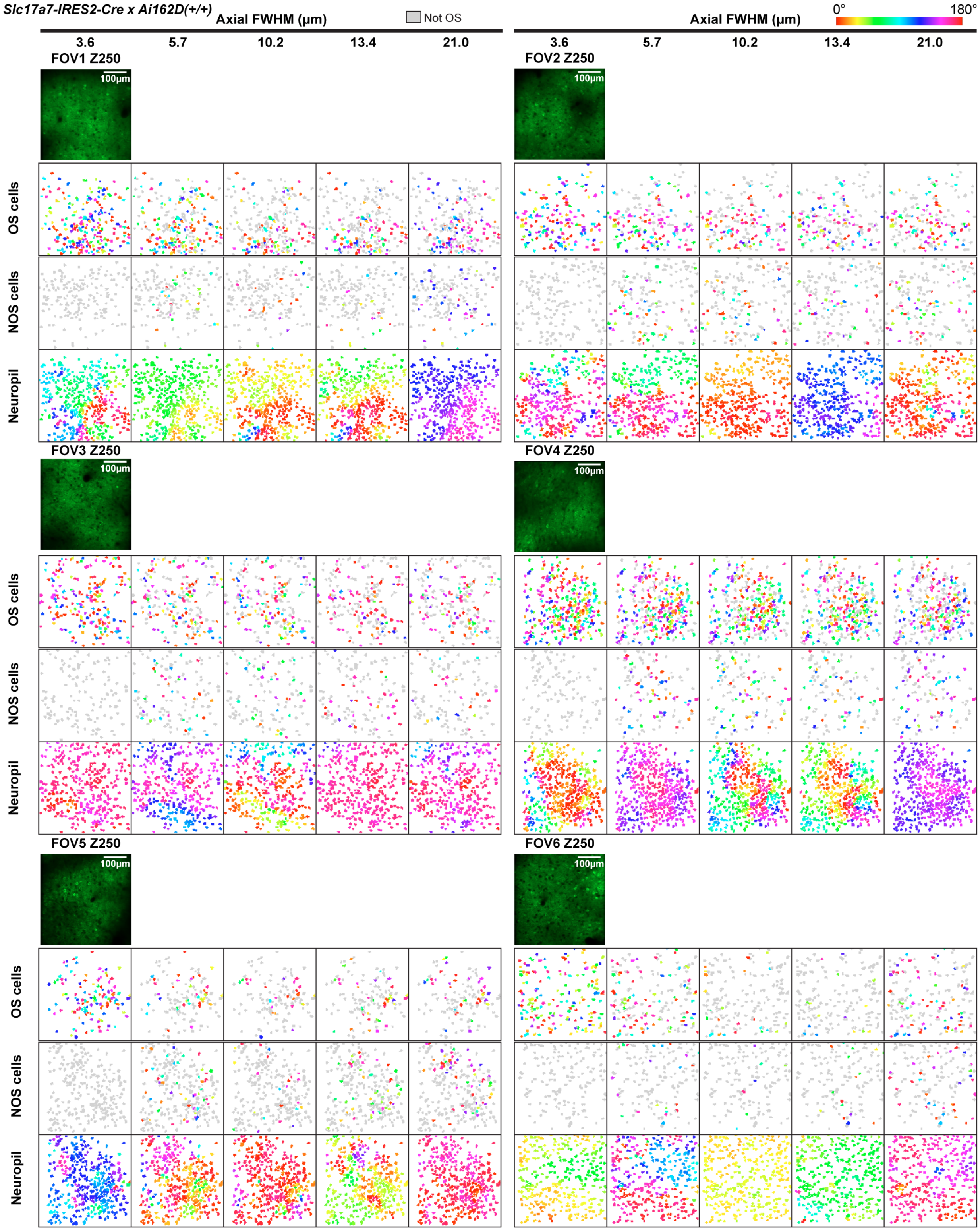


**Figure S3. Preferred orientation map of cells expressing GCaMP6s transgenically.** 2P fluorescence images of six FOVs and their color-coded preferred orientation (PO) maps for VR neurons in each FOV. Top and middle rows: maps for cells classified to be OS and not OS at 3.6-µm aFWHM, respectively, color-coded by preferred orientation (gray for not OS). Bottom row: map for corresponding neuropil. FOV1 is the same FOV as Fig. 3a.

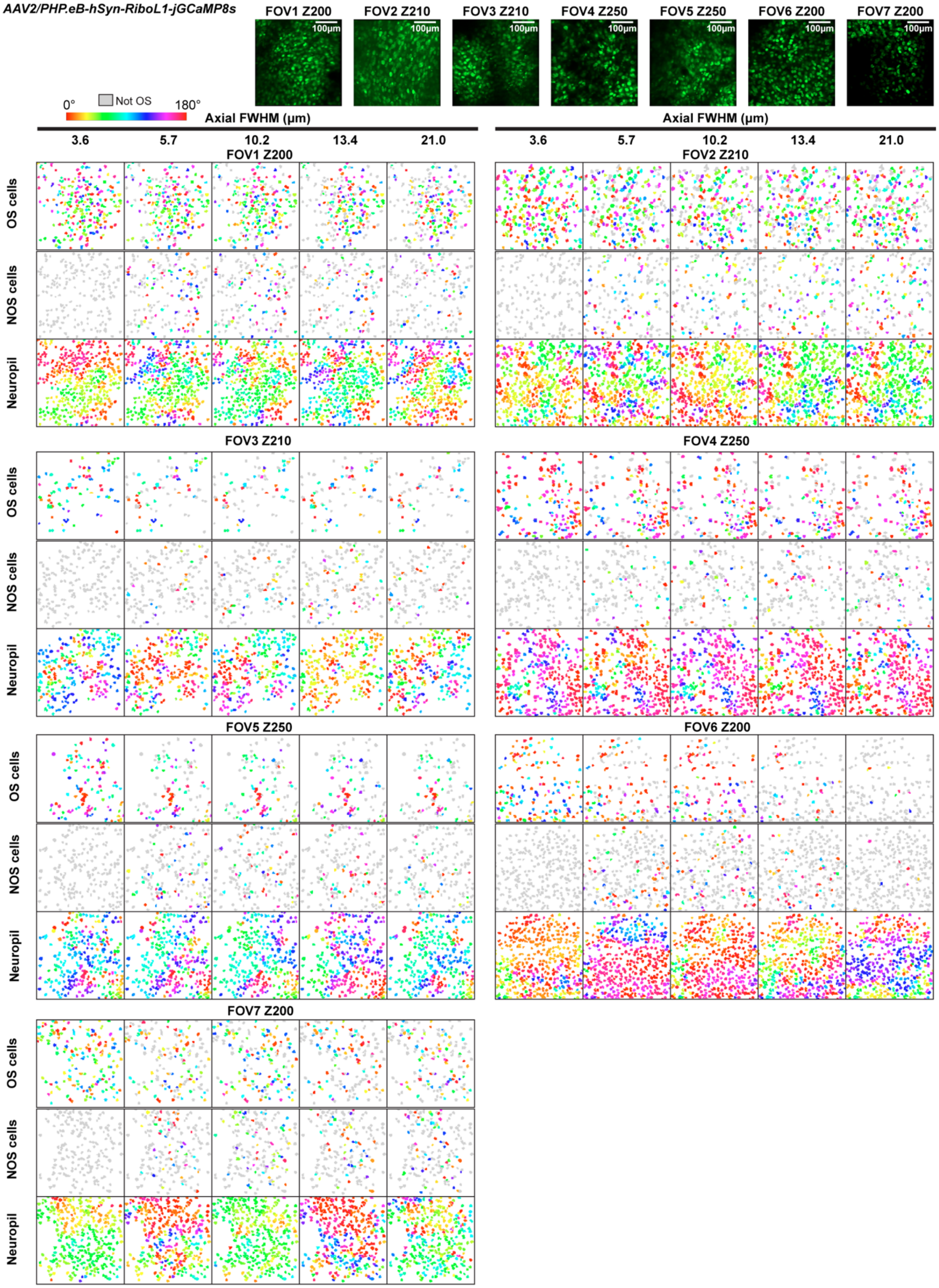


**Figure S4. Preferred orientation map of cells expressing soma-targeted RiboL1-jGCaMP8s via viral transduction.** 2P fluorescence images of six FOVs and their color-coded preferred orientation (PO) maps for VR neurons in each FOV. Top and middle rows: maps for cells classified to be OS and not OS at 3.6-µm aFWHM, respectively, color-coded by preferred orientation (gray for not OS). Bottom row: map for corresponding neuropil. FOV1 is the same FOV as Fig. 4a.

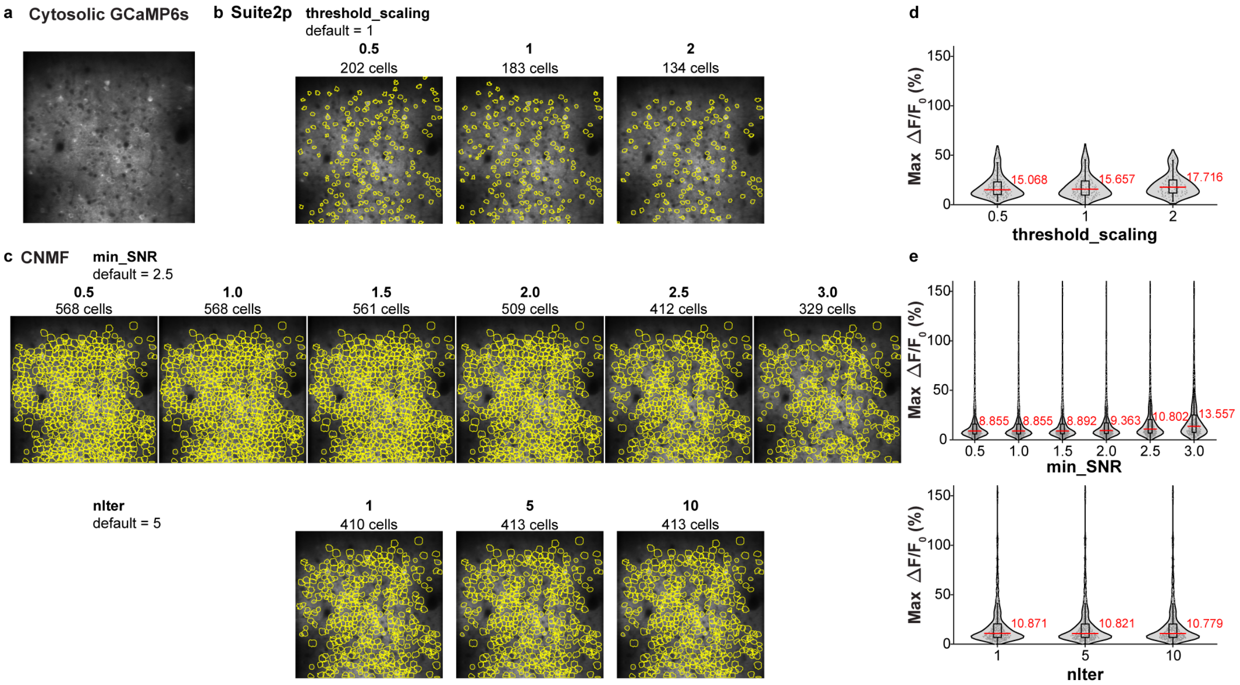


**Figure S5. ROI detection and maximal $\Delta F/F_0$ distribution vary moderately on parameter selection in Suite2p and CNMF. (a)** Example FOV of neurons expressing GCaMP6s transgenically. **(b,c)** Detected ROIs (yellow circles) for Suite2p and CNMF, respectively, at different values of "threshold_scaling" (for Suite2p), "min_SNR" and "nIter" (for CNMF). **(d,e)** Violin plots showing the maximal trial-averaged $\Delta F/F_0$ as a function of the parameter values for Suite2p and CNMF, respectively. Red lines and numbers: medians.

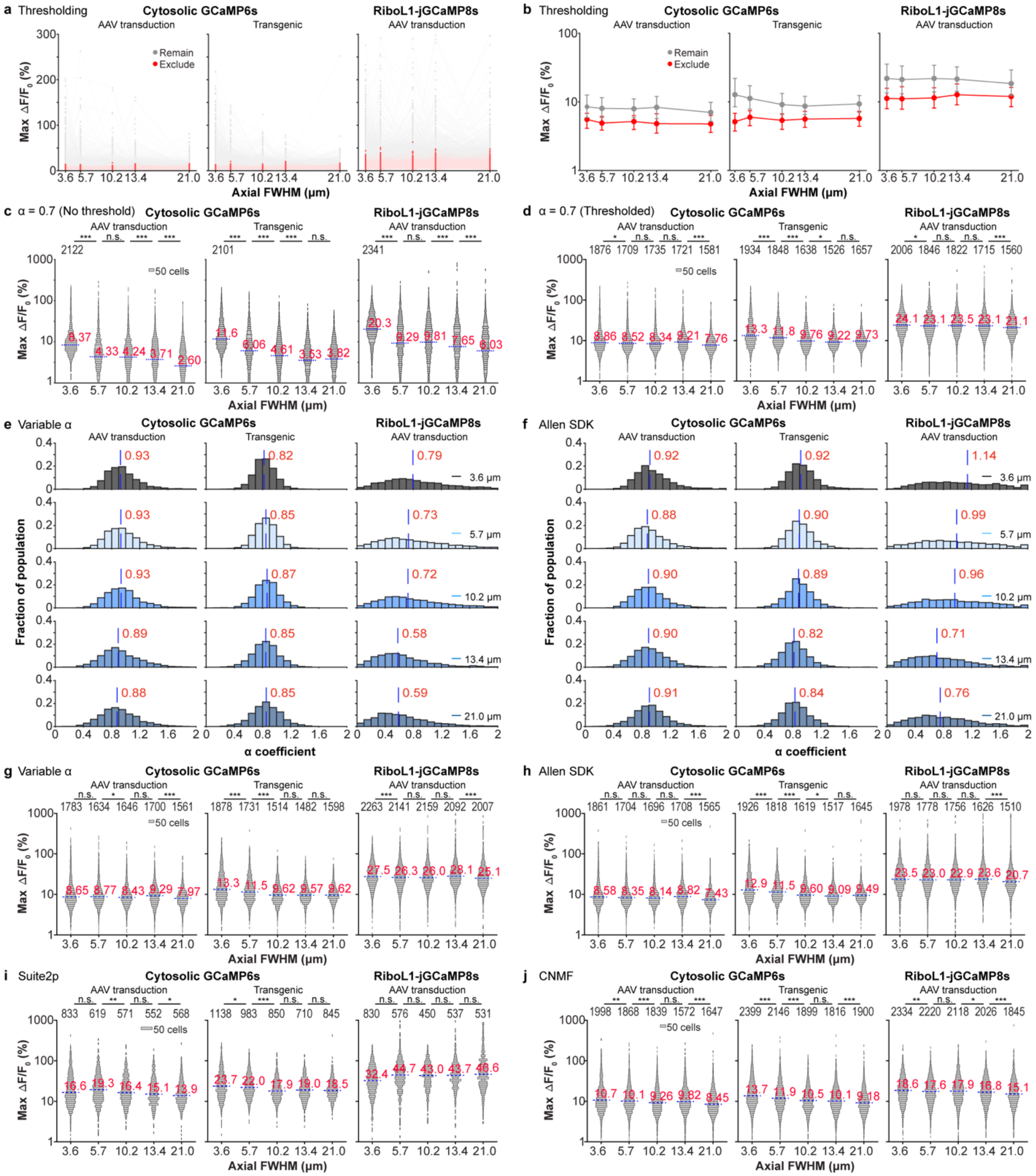


**Figure S6. Additional results for more advanced analysis pipelines. (a-d)** Consequences of SNR thresholding on 2,499, 2,452, and 2,810 hand-segmented ROIs and "α = 0.7" analysis pipeline for V1 L2/3 neurons with virally transduced or transgenic GCaMP6s expression, and virally transduced RiboL1-jGCaMP8s expression. **(a,b)** Scatter plots and population statistics, respectively, of the maximal trial-averaged $\Delta F/F_0$ values for manually segmented ROIs with SNR > 3.5 (gray symbols; passing SNR thresholding) or SNR ≤ 3.5 (red symbols). Dots and whiskers in **b**: median values with interquartiles. **(c,d)** Distributions of maximal trial-averaged $\Delta F/F_0$ values for neurons classified as VR before (using datasets acquired at 3.6-μm aFWHM; same data as **Figs. 2i,3i,4i**) and after SNR thresholding (classified as VR for each aFWHM condition independently), respectively. Here and below, one-sided WRS test against previous aFWHM conditions: n.s., not significant, $^*p < 0.05$, $^{**}p < 0.01$, and $^{***}p < 0.001$. Number of ROIs are labeled above each aFWHM condition here and below. **(e,f)** Histogram distributions of cell-specific α coefficients from "Variable α" and "Allen SDK", respectively. Blue lines and red numbers: medians. **(g,h)** Distributions of maximal trial-averaged $\Delta F/F_0$ values for neurons classified as VR (for each aFWHM condition independently) for "Variable α" and "Allen SDK", respectively. **(i,j)** Distributions of maximal trial-averaged $\Delta F/F_0$ values for neurons classified as VR (for each aFWHM condition independently) for "Suite2p" and "CNMF", respectively.

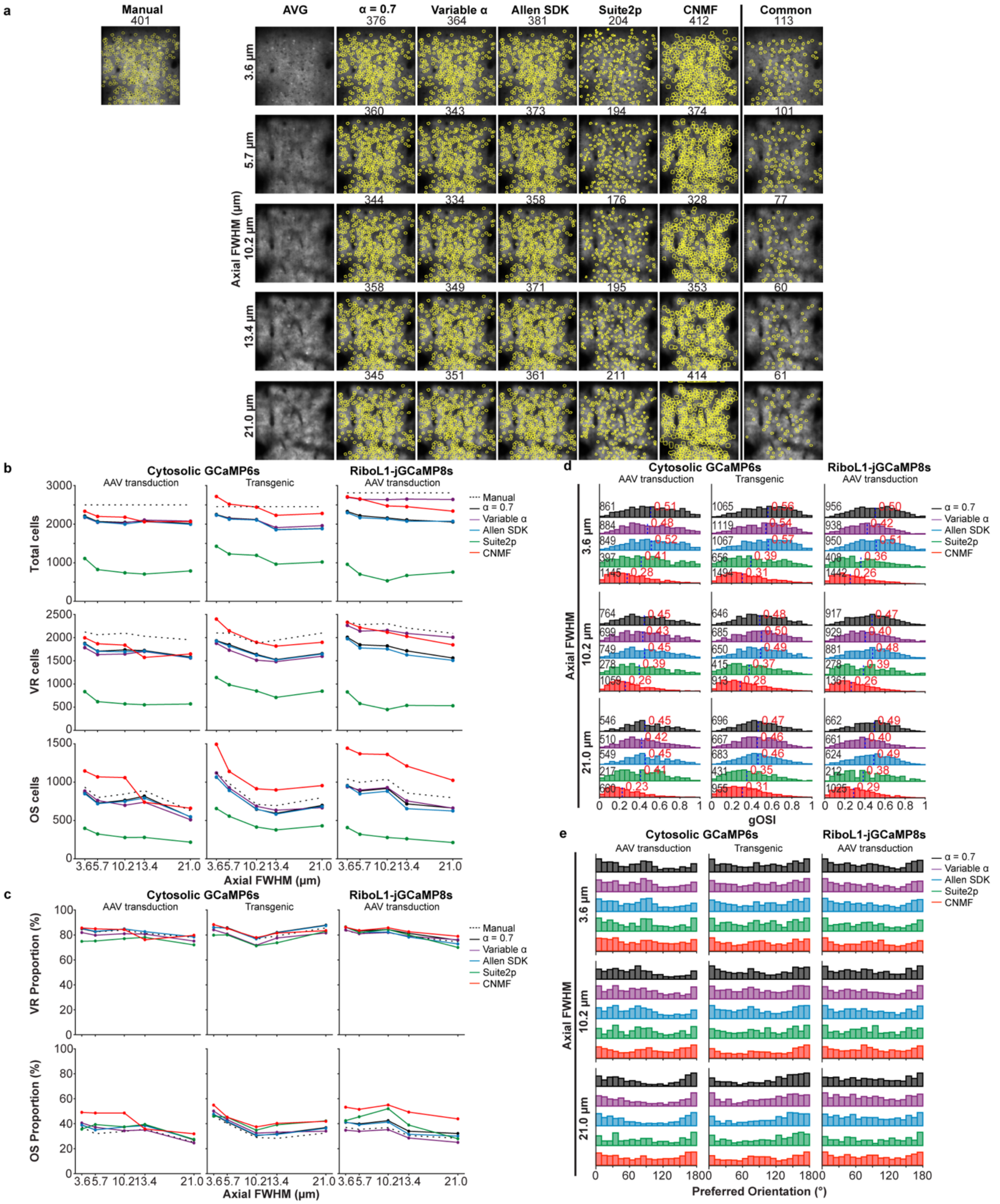

**Figure S7. Additional comparison of analysis pipelines.** **(a)** Left to right: 2P fluorescence images of an example FOV (same as **Fig. 3a**), the hand-segmented ROIs that pass SNR thresholding for “α=0.7”, “Variable α”, and “Allen SDK”, the automatically detected ROIs by Suite2p and CNMF, the ROIs that are common to all the preceding methods, and the ROIs hand-segmented from the dataset acquired at 3.6-µm aFWHM. Rows: all aFWHM conditions. **(b)** Top to bottom: total number of ROIs from all FOVs and animals, number of VR cells, number of OS cells versus aFWHM for all analysis pipelines and labeling strategies. **(c)** Proportions of VR and OS neurons versus aFWHM, calculated from data in **b**. **(d)** Histogram distributions of gOSI. Numbers at the left end of histograms: total number of cells. Blue lines and red numbers: medians. **(e)** Histogram distributions of preferred orientation of OS cells. (Pipeline key: black, α = 0.7; purple, Variable α; blue, Allen SDK; green, Suite2p; red, CNMF).

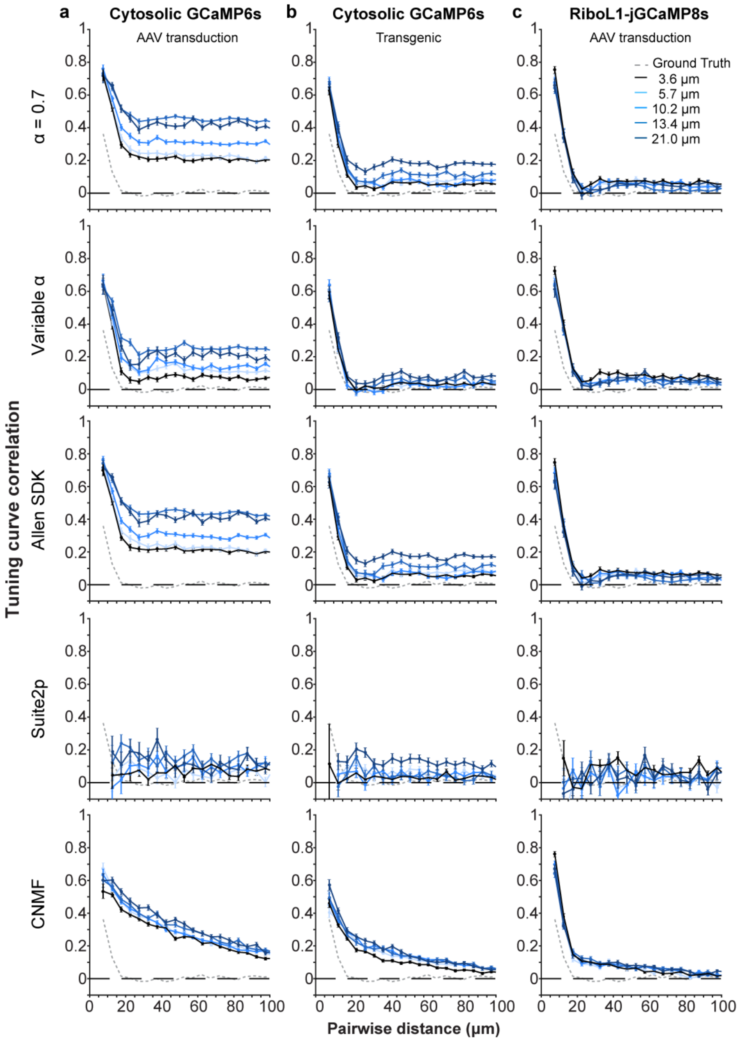


**Figure S8. Orientation tuning correlation for OS cells from all analysis pipelines and aFWHM conditions. (a)** Top to bottom: orientation tuning correlation vs. distance for datasets acquired at aFWHMs of 3.6, 5.7, 10.2, 13.4, and 21.0 µm and analyzed with "α = 0.7", "Variable α", "Allen SDK", Suite2p, and CNMF, respectively, for V1 L2/3 OS neurons with virally transduced GCaMP6s expression. **(b,c)** Same as **(a)**, but for neurons with transgenic GCaMP6s expression and virally transduced RiboL1-jGCaMP8s expression, respectively. Gray dotted traces: ground truth from data acquired using nuclear-targeted GCaMP6s. Error bars: s.e.m.

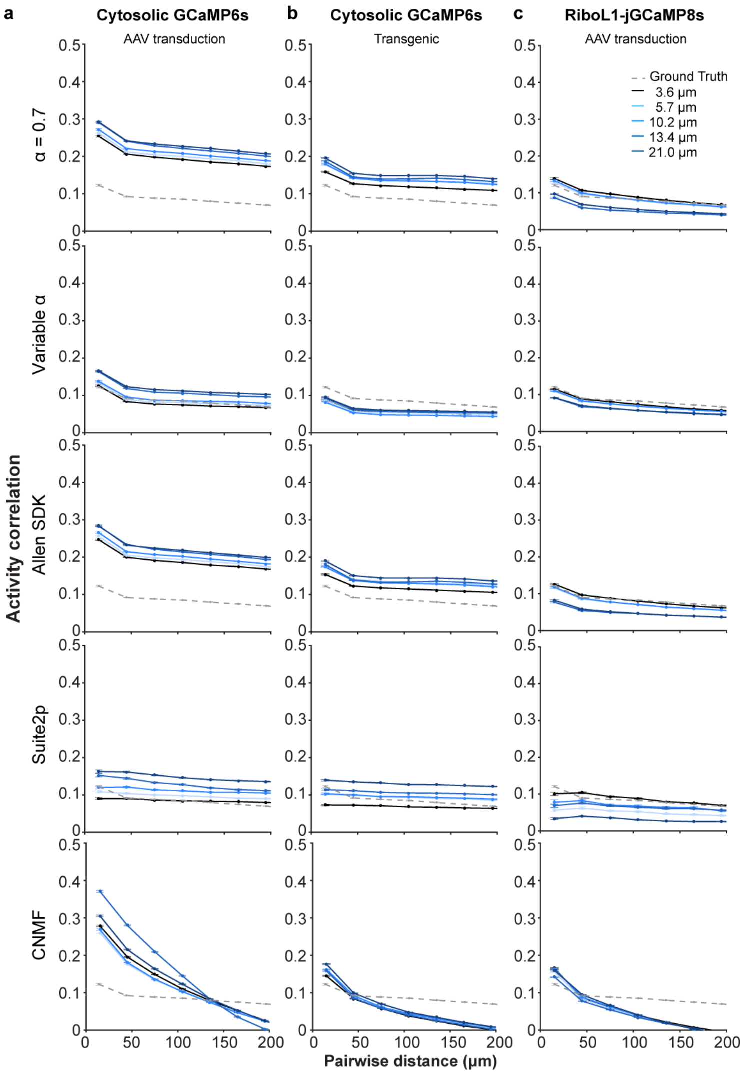


**Figure S9. Pairwise activity correlation analysis for all analysis pipelines and aFWHM conditions. (a)** Top to bottom: pairwise activity correlation vs. distance for datasets acquired at aFWHMs of 3.6, 5.7, 10.2, 13.4, and 21.0 μm and analyzed with "α = 0.7", "Variable α", "Allen SDK", Suite2p, and CNMF, respectively, for V1 L2/3 OS neurons with virally transduced GCaMP6s expression. **(b,c)** Same as **(a)**, but for neurons with transgenic GCaMP6s expression and virally transduced RiboL1-jGCaMP8s expression, respectively. Error bars: s.e.m.

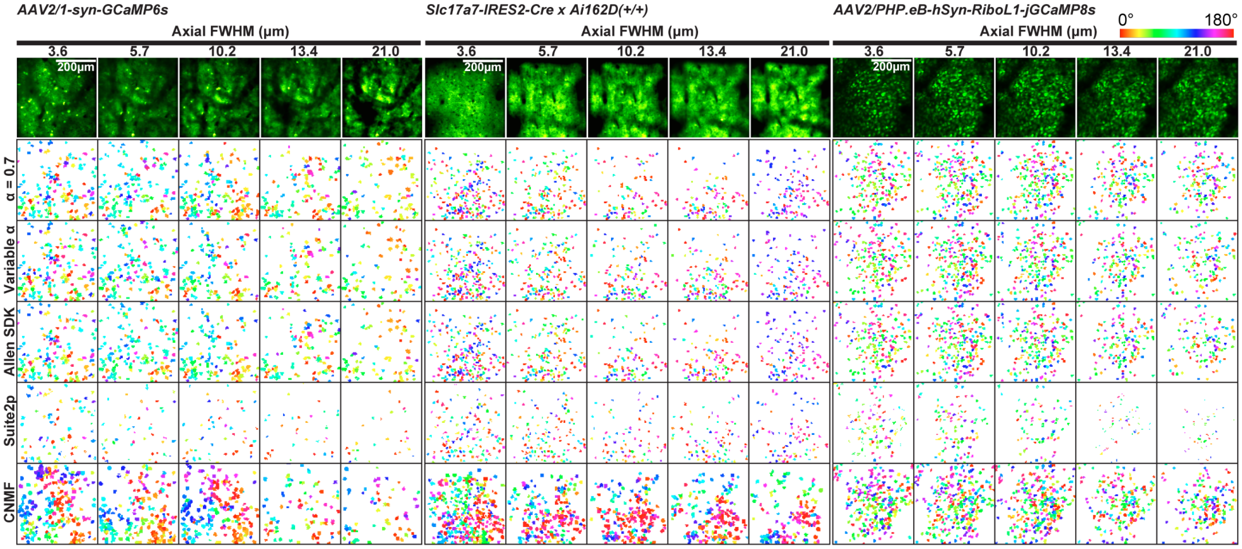


**Figure S10. ROI masks for OS neurons from different analysis pipelines**. Example preferred orientation maps for OS neurons in example FOVs with (from left to right) AAV-transduced cytosolic GCaMP6s, transgenic cytosolic GCaMP6s, and soma-targeted jGCaMP8s expression from imaging data acquired at different aFWHMs. Rows (top to bottom): 2P images, and preferred orientation maps from α = 0.7, Variable α, Allen SDK, Suite2p, and CNMF. OS cell masks are color-coded by their preferred orientation.

**Table S1. Evolution of two-photon microscopy for large-scale neural population imaging categorized by axial resolution.**

| Category | Author(s) | DOI | aFWHM (µm) | FOV Size ($mm^2$) | Analysis Pipeline (if used) |
|---|---|---|---|---|---|
| Large FOV | Tsai et al.<br>Opt. Express 2015 | 10.1364/OE.23.013833 | 16 - 33 | 8 x 10<br>80 $mm^2$ | - |
| Large FOV | Sofroniew et al.<br>eLife 2016 | 10.7554/eLife.14472 | 4.09 – 6.88 | Ø 5<br>19.6 $mm^2$ | Manual segmentation |
| Large FOV | Stirman et al.<br>Nat. Biotechnol. 2016 | 10.1038/nbt.3594 | 11.8 – 12.1 | Ø 3.5<br>9.6 $mm^2$ | Pixel-wise correlation or kurtosis map |
| Large FOV | Chen et al.<br>eLife 2016 | 10.7554/eLife.14679 | 4.5 – 10.5 | 1.7 x 1.7<br>2.9 $mm^2$ | Manual segmentation |
| Large FOV | Bumstead et al.<br>Neurophotonics 2018 | 10.1117/1.NPh.5.2.025001 | 28 | Ø 7<br>38.5 $mm^2$ | - |
| Large FOV | Terada et al.<br>Nat. Commun. 2018 | 10.1038/s41467-018-06058-8 | 9.96 | Ø 5.2 – 6.4<br>21.2 – 32.2 $mm^2$ | CNMF |
| Large FOV | Rumyantsev et al.<br>Nature 2020 | 10.1038/s41586-020-2130-2 | 8 | 2.0 x 2.0<br>4 $mm^2$ | PCA/ICA based sorting algorithm |
| Large FOV | Clough et al.<br>Nat. Commun. 2021 | 10.1038/s41467-021-26737-3 | 10.51 – 12.46 | Ø 3, Ø 4.8<br>7.1 – 18.1 $mm^2$ | Manual segmentation |
| Large FOV | Ota et al.<br>Neuron 2021 | 10.1016/j.neuron.2021.03.032 | 7.07 | 3.0 x 3.0<br>9 $mm^2$ | Custom code |
| Large FOV | Yu et al.<br>Nat. Commun. 2021 | 10.1038/s41467-021-26736-4 | ~8 | Ø 5<br>19.6 $mm^2$ | Suite2p |
| Large FOV | Yao et al.<br>Opt. Lett. 2022 | 10.1364/OL.450973 | 5.1 - 5.8 | Ø 3.46<br>9.4 $mm^2$ | Manual segmentation |
| Large FOV | Mok et al.<br>eLight 2024 | 10.1186/s43593-024-00076-4 | ~5 | 3.23 x 3.23<br>10.4 $mm^2$ | Manual segmentation, Suite2p |
| Large FOV | Yu et al.<br>Nat. Methods 2024 | 10.1038/s41592-023-02098-1 | 5.84 | 2.0 x 2.0<br>4 $mm^2$ | Manual segmentation |
| Large FOV | Agetsuma et al.<br>Cell Rep. Methods 2025 | 10.1016/j.crmeth.2025.101010 | 34 | 2.0 x 2.0<br>4 $mm^2$ | CNMF |
| Large FOV | Yang et al.<br>bioRxiv 2026 | 10.1101/2025.06.07.658419 | 10.6 – 13.1 | Ø 8<br>50.3 $mm^2$ | Suite2p |
| Fast axial scanning | Grewe et al.<br>Biomed. Opt. Express 2011 | 10.1364/BOE.2.002035 | 8 | 0.38 x 0.38<br>0.14 $mm^2$ | Manual segmentation |
| Fast axial scanning | Katona et al.<br>Nat. Methods 2012 | 10.1038/nmeth.1851 | 2.49 – 7.9 | 0.7 x 0.7 x 1.4 | Custom code |
| Fast axial scanning | Kong et al.<br>Nat. Methods 2015 | 10.1038/nmeth.3476 | 2.9 – 4.6<br>7.8 – 15.3 | 0.15 x 0.15 x .04<br>0.375 x 0.375 x .13 | Simple Neurite Tracer |
| Fast axial scanning | Prevedel et al.<br>Nat. Methods 2016 | 10.1038/nmeth.4040 | 10 | 0.5 x 0.5 | ICA based clustering |
| Fast axial scanning | Rupprecht et al.<br>Biomed. Opt. Express 2016 | 10.1364/BOE.7.001656 | 2.6 – 7 | - | - |
| Fast axial scanning | Liu et al.<br>Biomed. Opt. Express 2019 | 10.1364/BOE.10.005059 | 7 – 9.25 | 0.5 x 0.5 | Suite2p |
| Fast axial scanning | Weisenburger et al.<br>Cell 2019 | 10.1016/j.cell.2019.03.011 | 14.5 – 16.3 | 1 x 1 x 1.22 | CNMF (CaImAn) |
| Fast axial scanning | Chong et al.<br>Biomed. Opt. Express 2019 | 10.1364/BOE.10.000267 | 13.7 – 21.5 | 0.5 x 0.5 | - |
| Fast axial scanning | Beaulieu et al.<br>Nat. Methods 2020 | 10.1038/s41592-019-0728-9 | ~10 – 16 | - | Manual segmentation |
| Fast axial scanning | Chien et al.<br>Biomed. Opt. Express 2021 | 10.1364/BOE.405738 | 10.64 | 0.35 x 0.35 | Manual segmentation |
| Fast axial scanning | Demas et al.<br>Nat. Methods 2021 | 10.1038/s41592-021-01239-8 | 10 - 30 | 5.4 x 6 x 0.5 | CNMF (CaImAn) |
| Fast axial scanning | Janiak et al.<br>Nat. Commun. 2022 | 10.1038/s41467-022-28192-0 | 9.94 – 41.49 | 1.2 – 3.5<br>1.44 – 12.25 $mm^2$ | - |
| Miniaturized 2PFM | Göbel et al.<br>Opt. Lett. 2005 | 10.1364/OL.29.002521 | 20 | - | - |
| Miniaturized 2PFM | Flusberg et al.<br>Opt. Lett. 2005 | 10.1364/OL.30.002272 | 9.8 | Ø 0.145 - 0.215 | - |
| Miniaturized 2PFM | Engelbrecht et al.<br>Opt. Express 2008 | 10.1364/OE.16.005556 | 7.68 | Ø 0.20 | Custom correlation-based ICA algorithm |
| Miniaturized 2PFM | Hoy et al.<br>Opt. Express 2008 | 10.1364/OE.16.009996 | 16.4 | Ø 0.31 | - |
| Miniaturized 2PFM | Piyawattanametha et al.<br>Opt. Lett. 2009 | 10.1364/OL.34.002309 | 10.3 | 0.295 x 0.1 | Manual Segmentation |
| Miniaturized 2PFM | Ozbay et al.<br>Sci. Rep. 2018 | 10.1038/s41598-018-26326-3 | 10 | Ø 0.24 | Correlation-based clustering algorithm |
| Miniaturized 2PFM | Glas et al.<br>PLOS One 2019 | 10.1371/journal.pone.0214954 | 33.35 | 1 x 1 | Manual segmentation |
| Miniaturized 2PFM | Li et al.<br>Optica 2021 | 10.1364/OPTICA.422657 | 5.0<br>14.4 | Ø 0.14<br>Ø 0.3 | CNMF (CaImAn) |
| Miniaturized 2PFM | Zong et al.<br>Nat. Methods 2021 | 10.1038/s41592-020-01024-z | 12.18 | 0.42 x 0.42 | Manual segmentation |
| Miniaturized 2PFM | Zong et al.<br>Cell 2022 | 10.1016/j.cell.2022.02.017 | 12.8 – 17.8 | 0.42 x 0.42<br>0.5 x 0.5 | Suite2p |

| | | | | | |
|---|---|---|---|---|---|
| Miniaturized 2PFM | Accanto et al.<br>Neuron 2023 | 10.1016/j.neuron.2022.10.030 | 7-13 | Ø 0.25 | CNMF (CaImAn) |
| Miniaturized 2PFM | Wang et al.<br>Nano Letters 2023 | 10.1021/acs.nanolett.3c02439 | 18.08 | 0.045 x 0.045 | - |
| Miniaturized 2PFM | Zhao et al.<br>Opt. Express 2023 | 10.1364/OE.492674 | 24.64 | 1 x 0.788 | Manual segmentation |
| Miniaturized 2PFM | Futia et al.<br>bioRxiv 2024 | 10.1101/2024.10.21.619528 | 14 | 0.2 x 0.3 | CNMF (CaImAn) |
| Miniaturized 2PFM | Nardin et al.<br>Neurophotonics II 2024 | 10.1117/12.3022187 | 20 – 30 | 0.38 x 0.38 | - |
| Miniaturized 2PFM | Klioutchnikov et al.<br>bioRxiv 2025 | 10.1101/2025.08.01.668113 | 22.2 | 0.45 x 0.6 | Manual segmentation |
| Miniaturized 2PFM | Madruga et al.<br>Nat. Commun. 2025 | 10.1038/s41467-025-62534-y | 10.18 | 0.445 x 0.38 | Suite2p |
| Miniaturized 2PFM | Wu et al.<br>Nat. Methods 2025 | 10.1038/s41592-025-02780-6 | 3.73<br>7.04<br>23.68 | 0.6 x 0.26<br>0.5 x 0.425<br>1 x 0.8 | Manual segmentation |
| Miniaturized 2PFM | Zhang et al.<br>Cell Rep. Methods 2025 | 10.1016/j.crmeth.2025.101221 | 23.0 – 24.0 | 0.5 x 0.5 | CNMF |
| Miniaturized 2PFM | Blot et al.<br>Cell Rep. Methods 2026 | 10.1016/j.crmeth.2026.101305 | 9 - 13 | Ø 0.48 | - |

**Table S2. Normalized maximum pixel value and post objective power for different bead sizes measured across the various aFWHMs as shown in Figure 1b.**

| Bead size (μm) | Post objective power (mW) | | | |
|---|---|---|---|---|
| | aFWHM = 3.5 μm | aFWHM = 5.3 μm | aFWHM = 9.8 μm | aFWHM = 22.1 μm |
| 0.5 | 4.7 | 6.3 | 9.7 | 15.7 |
| 2 | 3.5 | 4.9 | 6.7 | 10.8 |
| 10 | 1.8 | 2.3 | 2.5 | 3.2 |
| 15 | 2.2 | 2.8 | 3.0 | 3.2 |

| Bead size (μm) | Signal ratios (mean ± standard error of mean) relative to 3.5-μm aFWHM | | | |
|---|---|---|---|---|
| | aFWHM = 3.5 μm | aFWHM = 5.3 μm | aFWHM = 9.8 μm | aFWHM = 22.1 μm |
| 0.5 | 1 | 0.414 ± 0.019 | 0.110 ± 0.020 | 0.018 ± 0.003 |
| 2 | 1 | 0.663 ± 0.049 | 0.217 ± 0.027 | 0.059 ± 0.015 |
| 10 | 1 | 0.866 ± 0.070 | 0.491 ± 0.063 | 0.208 ± 0.041 |
| 15 | 1 | 0.909 ± 0.054 | 0.581 ± 0.066 | 0.254 ± 0.042 |

**Table S3. Imaging acquisition parameters for all data collections.**

| Parameters | Figure 2 & 5, Figure S1-2 |
|---|---|
| Mouse line | Wildtype (C57BL/6J, male/female, aged 2-4 months) |
| Procedure | Virus injection (AAV2/1-syn-GCaMP6s::WPRE.SV40) and cranial window implantation |
| Post. obj. power (mW) | 78 - 160 |
| Excitation wavelength (nm) | 920 |
| Image size (μm x μm) | 400 x 400 |
| Pixel size (μm) | 1 |
| Frame rate (Hz) | 2 |

| Parameters | Figure 3 & 5, Figure S1 & 3 |
|---|---|
| Mouse line | Slc171a7-IRES2-Cre x Ai162D (+/+), JAX 023527 (male/female, aged 2-4 months) |
| Procedure | Cranial window implantation |
| Post. obj. power (mW) | 60 - 137 |
| Excitation wavelength (nm) | 920 |
| Image size (μm x μm) | 400 x 400 |
| Pixel size (μm) | 1 |
| Frame rate (Hz) | 2 |

| Parameters | Figure 4-5, Figure S1 & 4 |
|---|---|
| Mouse line | Wildtype (C57BL/6J, male/female, aged 2-4 months) |
| Procedure | Virus injection (AAV2/PEP.hB-hSyn-RiboL1-jGCaMP8s) and cranial window implantation |
| Post. obj. power (mW) | 96 - 201 |
| Excitation wavelength (nm) | 920 |
| Image size (μm x μm) | 400 x 400 |
| Pixel size (μm) | 1 |
| Frame rate (Hz) | 2 |

**Table S4. Statistical tests and p values for aFWHM condition comparisons for Figure 2, 3, and 4.**

AAV2/1-syn-GCaMP6s (**Figure 2**)

| Parameters | Test | p values | | | |
|---|---|---|---|---|---|
| | | aFWHM (μm) | | | |
| | | 5.7 | 10.2 | 13.4 | 21.0 |
| $F_{0,\ ROI}$ (**Figure 2g**) | Two-sample KS against 3.6-μm aFWHM | 0 | 0 | 0 | 0 |
| $\Delta F_{ROI}$ (**Figure 2h**) | Two-sample KS against 3.6-μm aFWHM | $2.00 \times 10^{-268}$ | $2.94 \times 10^{-257}$ | 0 | 0 |
| Maximal cell $\Delta F/F_0$ (**Figure 2i**) | One-sided WRS against previous aFWHM condition | $7.65 \times 10^{-168}$ | 0.176 | $2.35 \times 10^{-8}$ | $1.33 \times 10^{-9}$ |
| $F_{0,\ neuropil}$ (**Figure 2j**) | Two-sample KS against 3.6-μm aFWHM | 0 | 0 | 0 | 0 |
| $\Delta F_{neuropil}$ (**Figure 2k**) | Two-sample KS against 3.6-μm aFWHM | 0 | 0 | 0 | 0 |
| Maximal neuropil $\Delta F/F_0$ (**Figure 2l**) | One-sided WRS against previous aFWHM condition | 1 | $2.46 \times 10^{-94}$ | 1 | 1 |
| Cell gOSI (**Figure 2m,n**) | One-sided WRS against 3.6-μm aFWHM | $1.22 \times 10^{-5}$ | $8.43 \times 10^{-20}$ | $8.91 \times 10^{-13}$ | $2.26 \times 10^{-13}$ |
| Neuropil gOSI (**Figure 2o,p**) | One-sided WRS against 3.6-μm aFWHM | 0.993 | 0.405 | $9.61 \times 10^{-19}$ | $3.04 \times 10^{-55}$ |

Slc171a7-IRES2-Cre × Ai162D (+/+) (**Figure 3**)

| Parameters | Test | p values | | | |
|---|---|---|---|---|---|
| | | aFWHM (μm) | | | |
| | | 5.7 | 10.2 | 13.4 | 21.0 |
| $F_{0,\ ROI}$ (**Figure 3g**) | Two-sample KS against 3.6-μm aFWHM | 0 | 0 | 0 | 0 |
| $\Delta F_{ROI}$ (**Figure 3h**) | Two-sample KS against 3.6-μm aFWHM | $7.95 \times 10^{-270}$ | 0 | 0 | 0 |
| Maximal cell $\Delta F/F_0$ (**Figure 3i**) | One-sided WRS against previous aFWHM condition | $4.76 \times 10^{-140}$ | $1.71 \times 10^{-14}$ | $1.56 \times 10^{-13}$ | 0.80 |
| $F_{0,\ neuropil}$ (**Figure 3j**) | Two-sample KS against 3.6-μm aFWHM | 0 | 0 | 0 | 0 |
| $\Delta F_{neuropil}$ (**Figure 3k**) | Two-sample KS against 3.6-μm aFWHM | 0 | 0 | 0 | 0 |
| Maximal neuropil $\Delta F/F_0$ (**Figure 3l**) | One-sided WRS against previous aFWHM condition | $2.30 \times 10^{-16}$ | $1.37 \times 10^{-122}$ | $4.11 \times 10^{-29}$ | 0.013 |
| $F_{0,\ ROI}$ (**Figure 3g**) | One-sided WRS against 3.6-μm aFWHM | $4.65 \times 10^{-18}$ | $1.03 \times 10^{-36}$ | $2.08 \times 10^{-50}$ | $6.08 \times 10^{-50}$ |
| $\Delta F_{ROI}$ (**Figure 3h**) | One-sided WRS against 3.6-μm aFWHM | 0.173 | 0.981 | $1.63 \times 10^{-4}$ | $2.40 \times 10^{-29}$ |

AAV2/PEP.hB-hSyn-RiboL1-jGCaMP8s (**Figure 4**)

| **Parameters** | **Test** | **p values** | | | |
|---|---|---|---|---|---|
| | | **aFWHM (μm)** | | | |
| | | **5.7** | **10.2** | **13.4** | **21.0** |
| $F_{0,\,ROI}$ (**Figure 4g**) | Two-sample KS against 3.6-μm aFWHM | 0 | 0 | 0 | 0 |
| $\Delta F_{ROI}$ (**Figure 4h**) | Two-sample KS against 3.6-μm aFWHM | 0 | 0 | 0 | 0 |
| Maximal cell $\Delta F/F_0$ (**Figure 4i**) | One-sided WRS against previous aFWHM condition | $5.00 \times 10^{-135}$ | 0.927 | $1.26 \times 10^{-10}$ | $2.14 \times 10^{-5}$ |
| $F_{0,\,neuropil}$ (**Figure 4j**) | Two-sample KS against 3.6-μm aFWHM | 0 | 0 | 0 | 0 |
| $\Delta F_{neuropil}$ (**Figure 4k**) | Two-sample KS against 3.6-μm aFWHM | 0 | 0 | 0 | 0 |
| Maximal neuropil $\Delta F/F_0$ (**Figure 4l**) | One-sided WRS against previous aFWHM condition | $1.83 \times 10^{-13}$ | 0.555 | $2.58 \times 10^{-8}$ | 0.001 |
| $F_{0,\,ROI}$ (**Figure 4g**) | One-sided WRS against 3.6-μm aFWHM | $1.84 \times 10^{-9}$ | $4.98 \times 10^{-9}$ | $3.54 \times 10^{-15}$ | $5.08 \times 10^{-18}$ |
| $\Delta F_{ROI}$ (**Figure 4h**) | One-sided WRS against 3.6-μm aFWHM | 1 | 0.980 | 1 | 1 |

**Table S5. Statistical tests and p values for OS misclassifications as aFWHM increases for Figure S1.**

AAV2/1-syn-GCaMP6s

| Parameters | Test | Conditions | p values | | | |
|---|---|---|---|---|---|---|
| | | | aFWHM (μm) | | | |
| | | | 5.7 | 10.2 | 13.4 | 21.0 |
| $F_{0,\ ROI}$ (**Figure S1a**) | Two-sample KS against OS cases | False negative | 0.001 | $3.92 \times 10^{-5}$ | 0.315 | 0.040 |
| | | False positive | 0.204 | 0.050 | 0.002 | 0.003 |
| $F_{0,\ neuropil}$ (**Figure S1b**) | Two-sample KS against OS cases | False negative | 0.027 | $1.04 \times 10^{-4}$ | $1.57 \times 10^{-4}$ | 0.001 |
| | | False positive | 0.319 | 0.378 | $2.79 \times 10^{-4}$ | 0.005 |

Slc171a7-IRES2-Cre × Ai162D (+/+)

| Parameters | Test | Conditions | p values | | | |
|---|---|---|---|---|---|---|
| | | | aFWHM (μm) | | | |
| | | | 5.7 | 10.2 | 13.4 | 21.0 |
| $F_{0,\ ROI}$ (**Figure S1c**) | Two-sample KS against OS cases | False negative | $1.88 \times 10^{-5}$ | 0.041 | 0.002 | 0.004 |
| | | False positive | $2.24 \times 10^{-4}$ | $3.06 \times 10^{-4}$ | 0.024 | 0.510 |
| $F_{0,\ neuropil}$ (**Figure S1d**) | Two-sample KS against OS cases | False negative | 0.013 | 0.017 | 0.012 | 0.007 |
| | | False positive | 0.022 | $1.74 \times 10^{-4}$ | 0.003 | 0.320 |

AAV2/PEP.hB-hSyn-RiboL1-jGCaMP8s

| Parameters | Test | Conditions | p values | | | |
|---|---|---|---|---|---|---|
| | | | aFWHM (μm) | | | |
| | | | 5.7 | 10.2 | 13.4 | 21.0 |
| $F_{0,\ ROI}$ (**Figure S1e**) | Two-sample KS against OS cases | False negative | $3.72 \times 10^{-7}$ | $1.75 \times 10^{-7}$ | $2.89 \times 10^{-10}$ | $2.86 \times 10^{-7}$ |
| | | False positive | 0.034 | 0.002 | 0.052 | 0.123 |
| $F_{0,\ neuropil}$ (**Figure S1f**) | Two-sample KS against OS cases | False negative | $3.28 \times 10^{-5}$ | 0.002 | $1.54 \times 10^{-5}$ | $8.67 \times 10^{-7}$ |
| | | False positive | 0.053 | 0.004 | 0.110 | 0.103 |

**Table S6. Mean and standard error of maximal $\Delta F/F_0$ for VR cells, across pipelines and GECI strategies for Figure S6 (median plotted in the Figure S6).**

| | **AAV cytosolic GCaMP6s** | | **Transgenic GCaMP6s** | | **Soma-targeted jGCaMP8s** | |
|---|---|---|---|---|---|---|
| aFWHM | **3.6 μm** | **21.0 μm** | **3.6 μm** | **21.0 μm** | **3.6 μm** | **21.0 μm** |
| α = 0.7 (**Figure S6c**) | 13.1 ± 0.409 | 8.45 ± 0.135 | 19.6 ± 0.467 | 10.3 ± 0.174 | 28.7 ± 0.522 | 38.7 ± 8.62 |
| α = 0.7 (trace SNR threshold) (**Figure S6d**) | 14.1 ± 0.458 | 9.16 ± 0.160 | 20.8 ± 0.498 | 11.3 ± 0.211 | 31.3 ± 0.586 | 47.0 ± 11.6 |
| Variable α (trace SNR threshold) (**Figure S6g**) | 14.5 ± 0.522 | 9.46 ± 0.177 | 21.4 ± 0.547 | 11.2 ± 0.168 | 35.8 ± 0.611 | 41.6 ± 3.79 |
| Allen SDK (trace SNR threshold) (**Figure S6h**) | 13.6 ± 0.449 | 9.15 ± 0.327 | 21.7 ± 0.504 | 13.6 ± 0.800 | 31.2 ± 0.703 | 39.3 ± 3.08 |
| Suite2p (**Figure S6i**) | 28.4 ± 1.18 | 19.8 ± 0.822 | 33.9 ± 0.926 | 23.0 ± 0.672 | 42.3 ± 1.23 | 80.8 ± 5.40 |
| CNMF (**Figure S6j**) | 17.4 ± 0.502 | 12.2 ± 0.420 | 21.6 ± 0.485 | 12.5 ± 0.305 | 24.4 ± 0.433 | 21.1 ± 0.591 |

**Table S7. One-sided WRS statistical test against previous aFWHM conditions and resulting p values for trial-and-time-averaged $\Delta F/F_0$ distribution for VR cells, across pipelines and GECI strategies as shown in Figure S6.**

| Populations | Pipelines | p values | | | |
|---|---|---|---|---|---|
| | | aFWHM (μm) | | | |
| | | **5.7** | **10.2** | **13.4** | **21.0** |
| AAV2/1-syn-GCaMP6s | α = 0.7 (trace SNR threshold) (**Figure S6d**) | 0.043 | 0.062 | 0.996 | $1.69 \times 10^{-15}$ |
| | Variable α (trace SNR threshold) (**Figure S6g**) | 0.436 | 0.020 | 0.999 | $1.30 \times 10^{-13}$ |
| | Allen SDK (trace SNR threshold) (**Figure S6h**) | 0.100 | 0.074 | 0.988 | $3.65 \times 10^{-16}$ |
| | Suite2p (**Figure S6i**) | 0.988 | 0.004 | 0.080 | 0.015 |
| | CNMF (**Figure S6j**) | 0.008 | $5.15 \times 10^{-7}$ | 0.999 | $7.36 \times 10^{-10}$ |
| Slc171a7-IRES2-Cre × Ai162D (+/+) | α = 0.7 (trace SNR threshold) (**Figure S6d**) | $2.07 \times 10^{-10}$ | $4.46 \times 10^{-24}$ | 0.011 | 0.988 |
| | Variable α (trace SNR threshold) (**Figure S6g**) | $5.19 \times 10^{-11}$ | $5.15 \times 10^{-16}$ | 0.425 | 0.482 |
| | Allen SDK (trace SNR threshold) (**Figure S6h**) | $2.17 \times 10^{-9}$ | $9.25 \times 10^{-24}$ | 0.016 | 0.989 |
| | Suite2p (**Figure S6i**) | 0.017 | $1.54 \times 10^{-11}$ | 0.832 | 0.470 |
| | CNMF (**Figure S6j**) | $9.09 \times 10^{-12}$ | $5.40 \times 10^{-12}$ | 0.287 | $7.81 \times 10^{-7}$ |
| AAV2/PEP.hB-hSyn-RiboL1-jGCaMP8s | α = 0.7 (trace SNR threshold) (**Figure S6d**) | 0.013 | 0.900 | 0.300 | $6.96 \times 10^{-6}$ |
| | Variable α (trace SNR threshold) (**Figure S6g**) | $7.60 \times 10^{-4}$ | 0.659 | 0.999 | $9.99 \times 10^{-10}$ |
| | Allen SDK (trace SNR threshold) (**Figure S6h**) | 0.134 | 0.697 | 0.940 | $2.48 \times 10^{-7}$ |
| | Suite2p (**Figure S6i**) | 1 | 0.135 | 0.834 | 0.927 |
| | CNMF (**Figure S6j**) | 0.008 | 0.776 | 0.029 | $2.98 \times 10^{-4}$ |

**Table S8. Mean gOSI for OS cells, across pipelines and GECI strategies as shown in Figure 5.**

| | **AAV cytosolic GCaMP6s** | | **Transgenic GCaMP6s** | | **Soma-targeted jGCaMP8s** | |
|---|---|---|---|---|---|---|
| aFWHM | **3.6 µm** | **21.0 µm** | **3.6 µm** | **21.0 µm** | **3.6 µm** | **21.0 µm** |
| $\alpha = 0.7$ (**Figure 5c**) | 0.51 | 0.48 | 0.55 | 0.47 | 0.50 | 0.48 |
| $\alpha = 0.7$ (trace SNR threshold) (**Figure 5d**) | 0.52 | 0.49 | 0.55 | 0.48 | 0.51 | 0.48 |
| Variable $\alpha$ (trace SNR threshold) (**Figure 5g**) | 0.50 | 0.44 | 0.53 | 0.46 | 0.44 | 0.41 |
| Allen SDK (trace SNR threshold) (**Figure 5h**) | 0.53 | 0.49 | 0.55 | 0.48 | 0.51 | 0.48 |
| Suite2p (**Figure 5j**) | 0.44 | 0.42 | 0.43 | 0.38 | 0.38 | 0.42 |
| CNMF (**Figure 5l**) | 0.32 | 0.27 | 0.35 | 0.33 | 0.30 | 0.32 |

**Table S9. Post-objective power (mW) for all data collections.**

AAV2/1-syn-GCaMP6s

| **Figure S2** | **aFWHM conditions (µm)** | | | | |
|---|---|---|---|---|---|
| | **3.6** | **5.7** | **10.2** | **13.4** | **21.0** |
| FOV1 Z250 | 120 | 86 | 86 | 127 | 127 |
| FOV2 Z250 | 108 | 78.6 | 80.8 | 82.2 | 83.1 |
| FOV3 Z200 | 120 | 86 | 86 | 93 | 94 |
| FOV4 Z250 | 120 | 86 | 86 | 160 | 160 |
| FOV5 Z250 | 120 | 86 | 86 | 127 | 127 |
| FOV6 Z250 | 120 | 86 | 86 | 120 | 121 |

Slc171a7-IRES2-Cre × Ai162D (+/+)

| **Figure S3** | **aFWHM conditions (µm)** | | | | |
|---|---|---|---|---|---|
| | **3.6** | **5.7** | **10.2** | **13.4** | **21.0** |
| FOV1 Z250 | 120 | 102 | 100 | 135 | 136 |
| FOV2 Z250 | 74 | 60 | 60 | 89 | 90 |
| FOV3 Z250 | 90 | 68 | 68 | 101 | 103 |
| FOV4 Z250 | 137 | 101 | 98 | 89 | 90 |
| FOV5 Z250 | 137 | 101 | 98 | 89 | 90 |
| FOV6 Z250 | 106 | 79 | 78 | 89 | 90 |

AAV2/PEP.hB-hSyn-RiboL1-jGCaMP8s

| **Figure S4** | **aFWHM conditions (µm)** | | | | |
|---|---|---|---|---|---|
| | **3.6** | **5.7** | **10.2** | **13.4** | **21.0** |
| FOV1 Z200 | 145 | 112 | 109 | 126 | 127 |
| FOV2 Z210 | 145 | 112 | 109 | 174 | 176 |
| FOV3 Z210 | 150 | 112 | 109 | 174 | 176 |
| FOV4 Z250 | 145 | 112 | 109 | 199 | 201 |
| FOV5 Z250 | 145 | 112 | 109 | 126 | 127 |
| FOV6 Z200 | 145 | 101 | 98 | 138 | 139 |
| FOV7 Z200 | 145 | 105 | 102 | 96 | 98 |
